# ReSeq simulates realistic Illumina high-throughput sequencing data

**DOI:** 10.1101/2020.07.17.209072

**Authors:** Stephan Schmeing, Mark D. Robinson

## Abstract

In high-throughput sequencing data, performance comparisons between computational tools are essential for making informed decisions in the data processing from raw data to the scientific result. Simulations are a critical part of method comparisons, but for standard Illumina sequencing of genomic DNA, they are often oversimplified, which leads to optimistic results for most tools.

ReSeq improves the authenticity of synthetic data by extracting and reproducing key components from real data. Major advancements are the inclusion of systematic errors, a fragment-based coverage model and sampling-matrix estimates based on two-dimensional margins. These improvements lead to a better representation of the original k-mer spectrum and more faithful performance evaluations. ReSeq and all of its code are available at: https://github.com/schmeing/ReSeq

## Background

High-throughput sequencing has revolutionized biology and medicine since it allows a myriad of applications, such as studying entire genomes at base-pair resolution. The accuracy of the obtained results after applying computational methods heavily depends on collected data and the tools used to process it. This paper focuses on standard Illumina short-read sequencing of genomic DNA (gDNA) that is obtainable for almost every molecular biology lab. In order to fully capitalize on these datasets, it is important to know the best tools for a given task, the typical error modes of these tools and whether a result is robust to fluctuations in the data or changes in the analysis.

With the ever-growing number of computational tools, evaluating their performance across the various situations in which they are applied has become an essential part of bioinformatics [1, 2]. There are two fundamental ways of doing benchmarks and validations. On the one hand, results can be compared to an *estimated* ‘gold-standard’ ground truth derived from real data, which can be based on consensus or an independent dataset (e.g., technology). On the other hand, tools can be compared on synthetic data, which are simulated from a specified ground truth that defines the desired results. Both strategies can introduce biases, through deriving the ground truth or by failing to mimic the properties of real data, respectively. Ideally, a mix of assessments from real and synthetic data is used, where differences in the results can highlight biases and provide estimates of uncertainty.

On real data, performance evaluations require a ground truth and estimating it with methods similar or identical to evaluated methods can induce a bias. To reduce such a bias, evaluations often only take into account situations where a confident consensus can be determined or where an alternative technology delivers reliable results [3]. This limitation necessarily reduces the breadth of method comparisons. Sometimes, these shortcomings can be mitigated with deeper sequencing, by using multiple alternative technologies and by carefully selecting the set of methods that go into the estimation. For example, extensive effort went into generating datasets with ground truth to assess variant calling on human gDNA datasets [4], such as the Genome in a Bottle Consortium [5,6] and Platinum Genomes [7]. Their detailed variant truth sets were derived using a consensus from multiple aligner / variant caller combinations. Having multiple technologies or pedigree information further increases the confidence in their ground truth calls. Alternatively, as an example of consensus-free approaches, Li et al. used Pacific Biosciences SMRT sequencing (PacBio) to create two independent assemblies, each of a homozygous human cell line. The combined assemblies provide the ground truth for a synthetic diploid dataset [8]. The advantage is that no variant callers are used to estimate the truth, while the disadvantage is that PacBio-specific errors remain. Despite the effort that went into the three mentioned truth sets, their results do not agree on variant caller performance, neither by value (e.g. false-positive rates of single-nucleotide polymorphisms) nor by rank [8]. Due to an abundance of truth sets in this field, the uncertainty in the comparisons can at least be assessed with real data alone. In other areas of research, often no published datasets with an estimated ground truth are readily available. For example, assessing the influence of polyploidy on variant calls using real data is limited to concordance checks [9].

Simulated data are a cheap and orthogonal way to benchmark computational methods and can readily address the shortcomings of real data based evaluations (e.g., biases toward certain tools or against certain genomic regions). Additionally, robustness towards properties of data (e.g., error rates) can be easily assessed. However, accurate method assessments require that simulated data recapitulate the important features of real data and do not oversimplify or bias the challenge for tested methods. Despite many published simulators, research comparing simulations has so far neglected to test for these important features. Reviews include Escalona et al. [10], which did not show any benchmarks, and Alosaimi et al. [11], which based the performance report on sensitivity and precision of mapping (of simulated reads). Unfortunately, this metric says little about the quality of the simulation; for example, a simulator that samples from unique regions of the reference and does not include any errors would receive a perfect score. A proper benchmark requires in-depth testing across a range of use-cases, since the most important features to mimic from real data depend on the application. We assess here many aspects of real data and in particular whether the key features for assembly have been reproduced. Furthermore, we show using the example of mapping, how to evaluate the scope of simulations in a benchmark.

For Illumina gDNA datasets, the simulation frameworks ART [12], pIRS [13] and NEAT [14] represent the state-of-the-art. Additionally, BEAR [15] was included to evaluate the effect of their design choices that simplify metagenomic simulations. We show below that all current simulators do an unsatisfactory job of reproducing, e.g., the k-mer spectrum of real data, due to their incomplete models for coverage, quality and base calling. As a result, methods tested on simulated data score nearly perfectly [16], which has presumably encouraged the field to rely on real data alone for evaluations; for example, the Assemblathon 1 [17] used simulations, while GAGE [18] and Assemblathon 2 [19] in the following years used real data alone.

Table 1 lists the features and input for each of the simulators compared in this study. The two main simulation components are the coverage model and the quality and base-call model. Coverage in ART and BEAR is modelled uniformly, while pIRS and NEAT introduce a GC bias by comparing the coverage across the binned reference [20,21]. This procedure is close to the Loess model described by Benjamini and Speed [22], who show that their alternative model based on the GC of individual fragments results in superior predictions. Furthermore, the bias from the sequences *flanking* the start and end of fragments [23] are not taken into account by any simulator. Finally, ART, pIRS and BEAR draw DNA fragment lengths from a user-defined Gaussian distribution, while NEAT uses the empirical distribution from the input bam file.

**Table 1:** Overview of modelled features for the compared simulators.

|  | ART | pIRS | NEAT | BEAR | ReSeq |
| --- | --- | --- | --- | --- | --- |
| Coverage | Parameters | Parameters | Parameters | Parameters | Mapped reads/Parameters |
| GC bias | - | Mapped reads | Mapped reads | - | Mapped reads |
| Flanking bias | - | - | - | - | Mapped reads |
| Fragment length | Parameters | Parameters | Mapped reads | Parameters | Mapped reads |
| Reference-sequence bias | - | - | - | Hard-coded/(Reads) | Mapped reads |
| Base qualities | Reads | Mapped reads | Reads | Reads | Mapped reads |
| Sequence qualities | - | - | - | - | Mapped reads |
| Substitution errors | Hard-coded | Mapped reads | Hard-coded | Reads | Mapped reads |
| Systematic errors | - | - | - | - | Mapped reads |
| InDel errors | Parameters | Mapped reads | Hard-coded | Reads | Mapped reads |
| Variants | - | Two genomes | Vcf | - | Vcf |

Parameters: Manually selected parameters. Reads: Learned from raw reads. Mapped reads: Learned from mapped reads. Hard-coded: Cannot be changed. BEAR’s reference-sequence bias estimation from reads is design for metagenomics.

For the qualities and base-calls, ART draws from empirical distributions of position-dependent qualities and introduces substitution errors [24–26] according to the probability given by the quality values. InDels are inserted based on four user-specified rates for insertion/deletion in the first/second read. In contrast, pIRS, NEAT and BEAR draw quality values from a non-homogeneous Markov chain, where the quality depends on the last quality and the position in the read. pIRS then chooses a base call from a learned distribution depending on the quality, position and reference base, while the inserted InDels depend only on the position in the read. NEAT instead follows a decision tree, where the occurrences of errors (Substituition and InDels) only depends on the quality and the substituted nucleotide only on the reference base, while the length and nucleotides of InDels have constant probabilities. It is worth mentioning that in the current version of NEAT, only qualities can be trained, but the base-call and InDel distributions rely on pretrained values. Specifically for metagenomics, BEAR’s error model is learned from duplicated reads using DRISEE [27] and therefore does not require a reference. Its parameters are obtained from exponential regression on the substitution rates by nucleotide and position and on the InDel rates by position. A peculiar design choice of BEAR is to replace quality values at error positions with qualities generated by the error model instead of simply having error rates depend on the quality values. Notably, neither systematic, sequence-specific errors [28, 29], nor the relationship between quality and fragment length [30] are included by any simulator.

The ability for users to train simulators on real data is another important feature, because profiles require constant updates to changes in sequencers, chemistries, etc. and even without technology changes, developer-provided profiles are not always accurate for a given use case due to differences in genome contexts or fragmentation method (see Results). None of the simulators mentioned can be fully trained on real data, instead relying at least partially on user-provided parameters or hard-coded models (Table 1).

**Re**al **Seq**uence Reproducer provides well-tested functionality to estimate the necessary parameters from real data mapped to a reference. Based on these estimates, it produces synthetic data with a k-mer spectrum matching real data without ever directly using k-mer information (see below). Requiring a reference is not a big constraint, since one is needed for the simulation anyways and furthermore, with a modest penalty in accuracy, the ReSeq parameters can be estimated from a *de novo* assembly generated from the reads.

We show that ReSeq outperforms all competitors in terms of delivering a realistic simulation and therefore lays the methodological groundwork for accurate benchmarking of genomics tools.

## Results

ReSeq consists of three parts: statistics calculation, probability estimation, simulation (Fig. 1). By default, all three steps are run as a pipeline, but they can also be called individually. Statistics calculation extracts the necessary information from the mapped reads and the corresponding reference. Afterwards, probability estimation combines the extracted matrices into distributions. Finally, the simulation step draws from those distributions.

**Figure 1:**
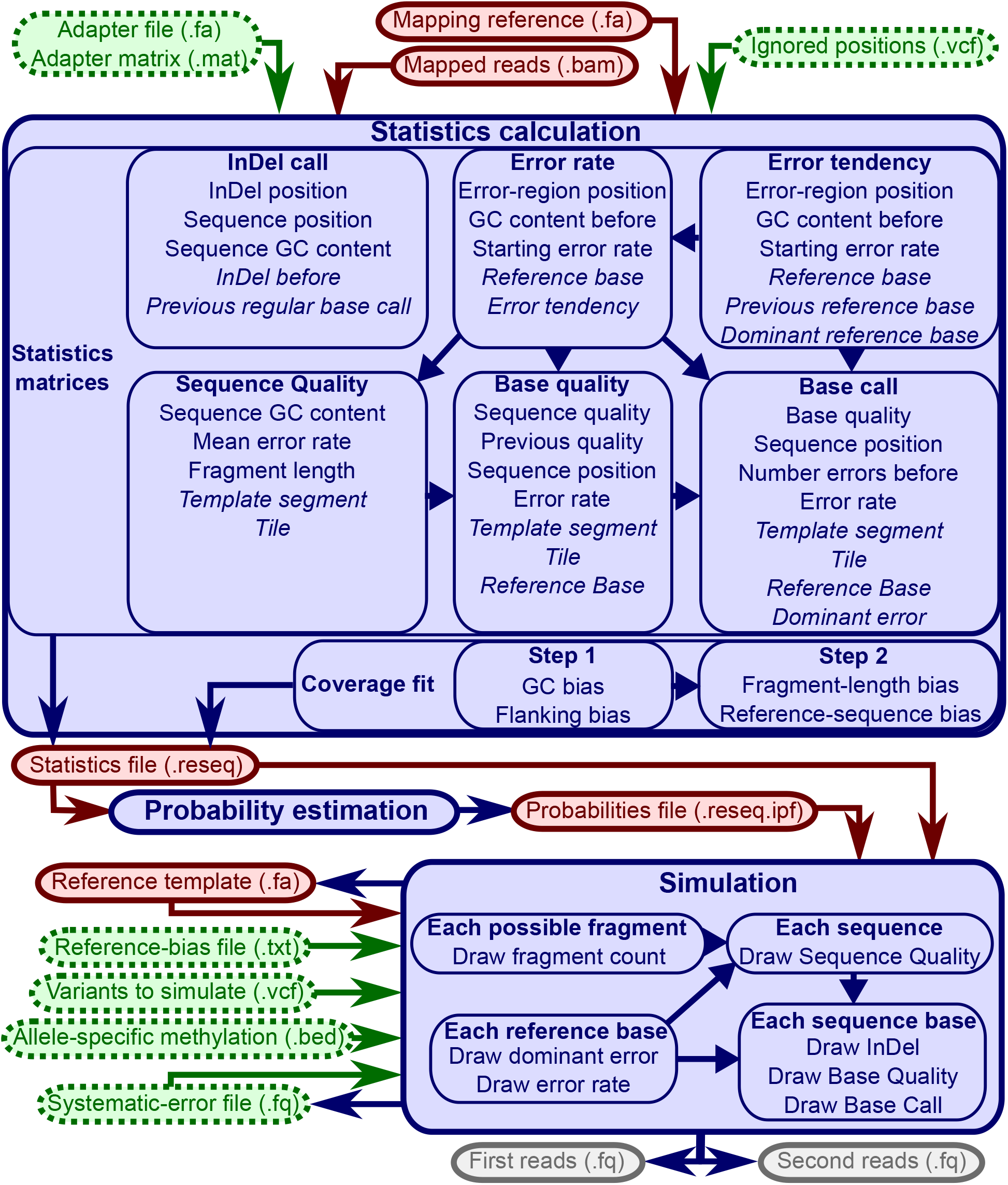
ReSeq overview. The three parts of ReSeq (solid, blue) with its mandatory (solid, red) and optional (dotted, green) inputs. The italic entries for the statistics matrices are independent dimensions in the matrix, while the normal entries are reduced to twodimensional interactions.

The statistics calculation has two components: estimating the bias parameters in the coverage model and populating matrices with statistics. A file with variants can be specified, such that their positions in the reference are excluded from the statistics. The bias estimation and statistics matrices will be discussed in detail in the next subsections.

The simulation produces synthetic data matching the calculated statistics and estimated biases. The reference provided can but does not need to be the same as the one used during the statistics calculation. To impose a clear separation, we will refer to the reference used during the simulation as the template. To simulate single-end reads, the second read file can simply be ignored, however paired-end Illumina data are still required for the statistics calculation. To properly handle coverage variations for sex chromosomes, mitochondria or metagenomics, the simulation optionally takes a reference-bias file (not necessary if template and reference are identical). To simulate diploid and polyploid genomes or pooled sequencing, variants can be specified for up to 64 alleles. To simulate bisulfite sequencing, allele-specific methylation values can be defined in an extended bed graph format with multiple score columns. However, we focus here on monoploid and diploid genomes and will not include simulations of bisulfite sequencing. For comparisons, we use eight representative datasets from different species, different Illumina machines and different adapters (Table 2).

**Table 2:**
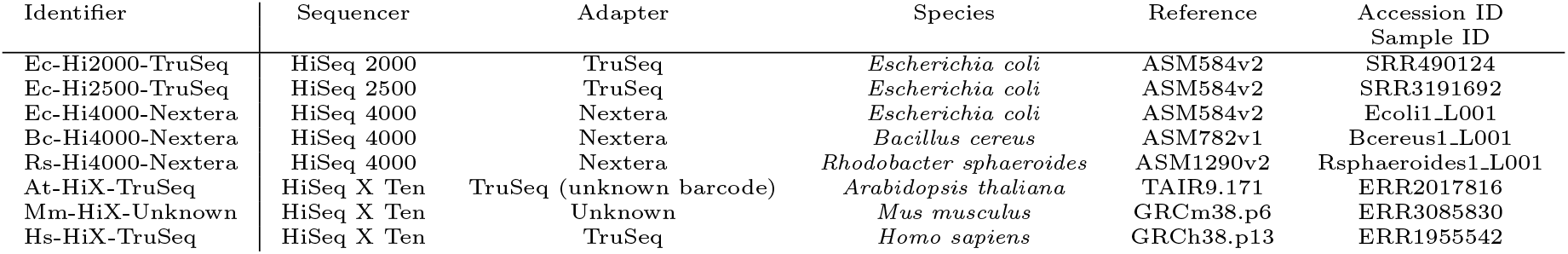
Used datasets.

Adapters labeled as unknown are not listed in the Illumina adapter overview [31].

### Quality and base calls

To simulate quality values and base calls, ReSeq fills six matrices during the statistics calculation (Fig. 1): insertions and deletions; systematic error rates at each reference position; systematic error tendencies at each reference position; sequence qualities; base qualities; and, base calls. The matrices are used to query the probability of the variable of interest conditional on all other variables in the matrix. For example, in a matrix containing the base quality *BQ*, previous quality *PQ* and sequence position *SP* we would query *p*(*BQ|PQ, SP*) for all *BQ*, which is a normalized slice of the matrix. These probabilities would then be used to draw the base quality.

However, due to the amount of variables included in each of these statistics, we cannot directly use large (sparse) matrices. For example, storing the quality values of Ec-Hi2000-TruSeq would require a matrix with 4.6 · 10^9^ entries. Therefore, ReSeq only retains twodimensional margins of the matrices. For the base-quality values, this means storing 10 two-dimensional margins for each template segment, tile and reference base combination: BQ - sequence quality (SQ), BQ - PQ, BQ - SP, BQ - error rate(ER), SQ - PQ, SQ - SP and so forth. This removes the higher-dimensional (3+) effects from the distributions, yet still provides a reasonable approximation (Fig. S1). The new set of marginal matrices has only around 3.0 · 10^5^ combined entries. This saves considerable computer memory, but more importantly prevents sparsity, because sufficient observations are required to sample accurate probability distributions in the absence of an analytical description. Using the full matrix, the conditional probability of a base quality *p*(*BQ|SQ, PQ, SP, ER*) would require many observations for every variable combination. In the reduced representation, the sampling only requires many observations for every two-dimensional combination of variables (i.e. SQ - PQ, SQ - SP, PQ - SP, etc.). Thus, the method requires much smaller input datasets.

To use the marginal matrices as a probability distribution during the simulation, they need to be converted back to a single matrix. This is done during the probability estimation step by combining the iterative proportional fitting procedure [32] with a data reduction technique described by Fienberg [33] (see Methods).

The most important new feature introduced by ReSeq is the representation of systematic errors. In older HiSeq2000 data, the systematic behaviour of errors is very pronounced (Fig. 2a,b), but also on the newer HiSeq4000 systematic errors are still present (Fig. 2c,d). Errors appear bundled at some positions and are strand dependent, which rules out variants as the cause. Ideally, systematic errors could be predicted on the read level. However, no model exists that predicts sequence-specific errors for a given position in a read based on all nucleotides in the same read preceding this position. Therefore, ReSeq distributes systematic errors randomly over the template by drawing error tendencies and rates for each template base on each strand. The five possible values for the error tendency are the four nucleotides and no tendency (i.e. random error).

**Figure 2:**
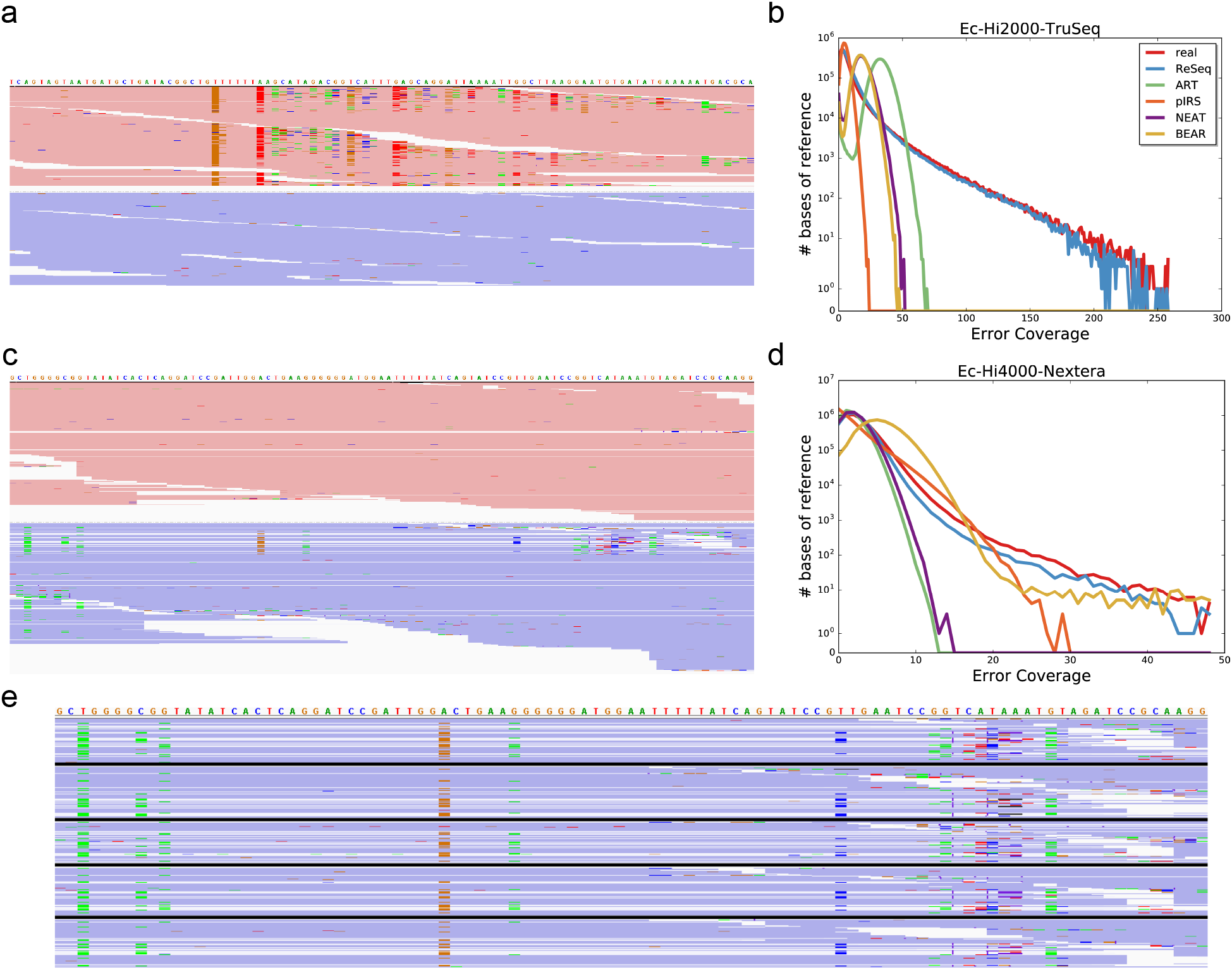
Systematic errors in Illumina data. a,c) Screenshots from the Integrative Genomics Viewer [34,35] for Ec-Hi2000-TruSeq(a) and Ec-Hi4000-Nextera(c). The forward strand is colored red and the reverse strand blue. The bright colors mark substitution errors. b,d) Amount of errors for all positions in the reference. e) Section from c with the same section from four technical replicates of Ec-Hi4000-Nextera (Ecoli1_L001): Ecoli1_L002, Ecoli1_L003, Ecoli2_L001, Ecoli3_L001. The datasets are separated by thick black lines.

To learn the distribution of systematic errors in the genome, they need to be distinguished from non-systematic (i.e. random) errors first. Since systematic-errors are strand specific, we assign a P-value to each reference position on each strand. The P-value is based on the number of occurrences of the most abundant error at this position (see Methods). The null hypothesis models random errors using a Poisson, which assumes equal error rates for every base in non-overlapping windows of the genome and conversion of bases with equal probability to the three possible erroneous nucleotides. We rejected the null hypothesis for adjusted P-values below 0.05.

The simulation starts by drawing the systematic error tendencies and associated rates. They can be stored and loaded to conserve the errors between multiple simulation runs, because real systematic errors are also very conserved, as highlighted in Figure 2e by comparing Ec-Hi4000-Nextera (Ecoli1_L001) with technical replicates from the same library run on different lanes (Datasets Ecoli1_L002 and Ecoli1_L003) and from separately prepared libraries sequenced on the same lane (Datasets Ecoli2_L001 and Ecoli3_L001). After setting the systematic errors, ReSeq loops through the template and distributes reads according to the bias model described in the next subsection. For each read, it starts by selecting a sequence quality. Then, each position in a read is handled by drawing an insertion, deletion or regular call. If a regular call was drawn, a quality value and base call are assigned. For insertions, ReSeq only draws quality values, because the nucleotides are already defined by the insertions themselves. The probabilities of all drawn values are based on many variables that are listed in Figure 1 and explained in more detail in the Methods section.

### Coverage model

The coverage model defines the probabilities of where simulated reads start and end in the simulated genome (template). It works on fragment sites, which are defined by their reference sequence, start position, fragment length and strand. Sites have been shown to be a good choice to estimate GC bias [22]. Since in our model duplications are fragments falling on the same site, the strand was added to make duplications strand-specific. During the statistics calculation, ReSeq determines all possible sites in a genome and counts the read pairs mapping to them. Sites, where the true counts are likely to deviate from the observed counts, e.g. repeat regions, are removed by looking at clusters of low mapping quality (see Methods).

The model takes four different sources of bias into account: GC and flanking sequence in a first step and fragment length and reference sequence in a second step. The flanking bias arises from the nucleotides *flanking* the start and end of the fragment as a result of the fragmentation process. We observed that their effect on the coverage is especially pronounced if enzyme digestion is used (Fig. S2). The GC bias is due to PCR [21], while the fragmentlength bias results from the fragmentation and size selection. Lastly, the reference-sequence bias represents the original abundance of the sequence in the sample, e.g. one half for male sex chromosomes or the microbial abundance for metagenomics.

In the first step of the coverage estimation, the GC and flanking bias are fit to the sites and their counts. To limit the memory requirements, sites are separated into combinations of reference sequence and fragment length. From all possible combinations, ReSeq selects only a small fraction and performs one fit each. Since modelling the counts at each site with a Poisson would only account for statistical duplications, we use a negative binomial to additionally account for PCR and optical duplications. This leads to the full likelihood over all sites *n*:

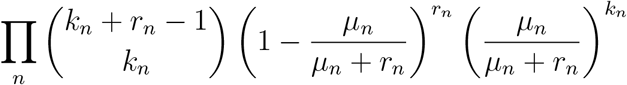

where *k_n_* is the observed count, *μ_n_* is the mean and *r_n_* is the dispersion at site *n*. Since coverage biases arise from different processes, we assume independence and write the mean as a product:

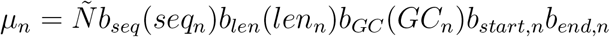

with 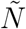 as the genome-wide normalization and the different *b* as the biases for this site. We cannot determine *b_seq_* and *b_len_*, because they are identical for all sites *n* within a fit. Thus, those biases are absorbed into the normalization:

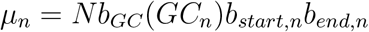

This model could alternatively be described as a generalized linear model with a logarithm link. However, the sheer size of the necessary matrix of explanatory variables makes it computationally prohibitive. Furthermore, the flanking bias uses the logit link function internally.

The normalization parameter is of no further interest after the fit, leaving 128 relevant parameters for the negative binomial. The GC bias *b_GC_* (Fig. 3a) is binned by percent into 101 bins that are described by a natural cubic spline with six knots, i.e. six degrees of freedom.

**Figure 3:**
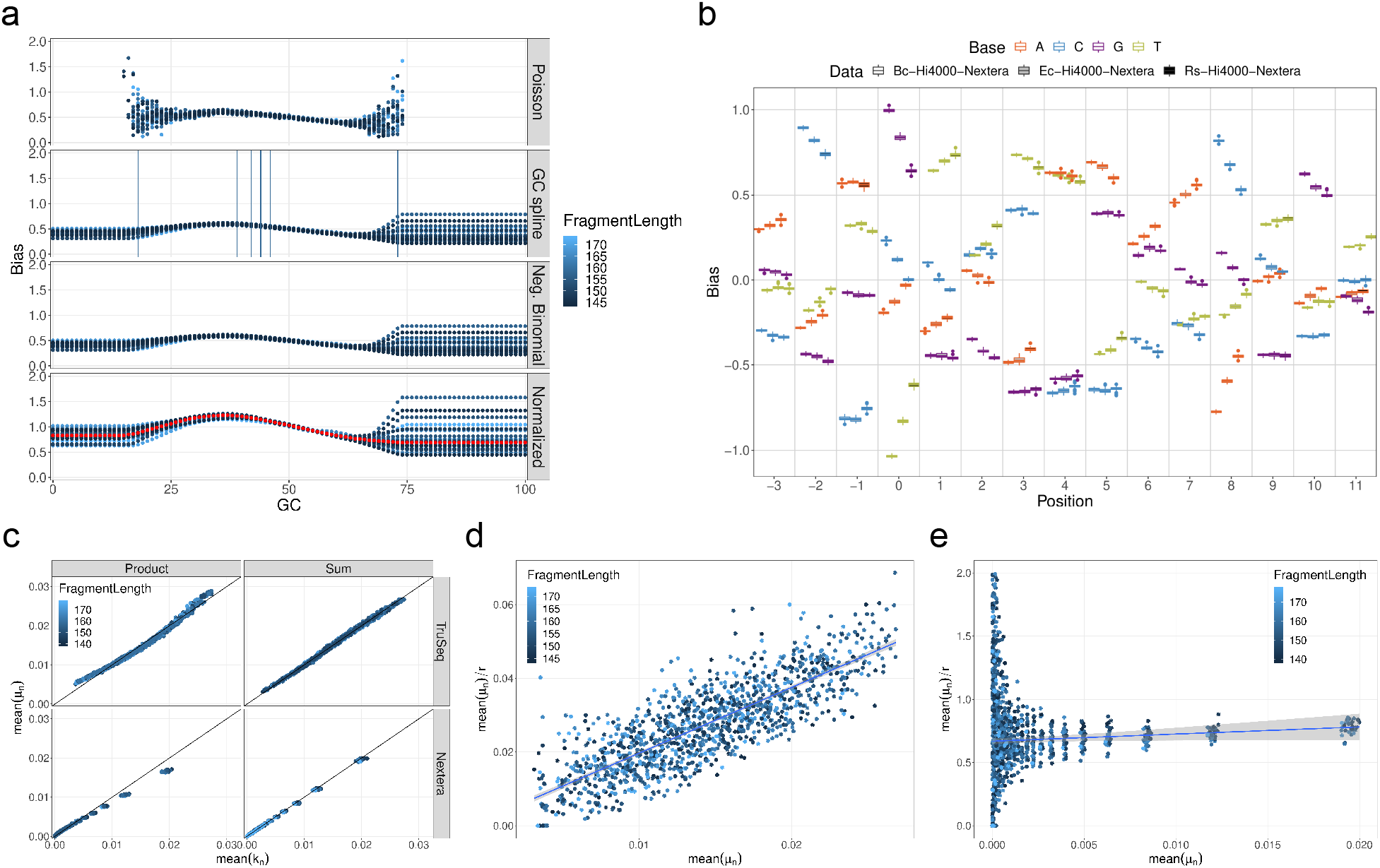
Coverage model. a) GC bias for Ec-Hi2000-TruSeq at the four steps of the bias fit for 30 fits with different fragment lengths. The red dots in the Normalized panel are the median value and represent the final result. The horizontal lines in the GC spline panel are the chosen knots for one example fragment length. b) Flanking bias. The effect of nucleotides in the genome relative to the fragment start or end with position 0 being the start/end. Negative positions are outside of the fragment. Only position −3 to 11 are shown from the total model that includes position −10 to 19. Each box summarises 30 to 40 fits with different fragment lengths. The boxes are arranged around their true position for improved readability. The three datasets are all created with Nextera adapters. c) Comparison of combining the different positions in the flanking bias by a product or a sum. Each dot is one bin of 2 · 10^5^ fragment sites for one of 30 fits with different fragment length. The fragment sites are ordered by their predicted mean counts *μ_n_* before binning. The x-axis is the mean of observed counts in the bin. The y-axis is the mean of predicted mean counts. For the sum the dots scatter around the identity, while for the product a curve is visible. d,e) The observed counts *k_n_* for the bins defined in c are fitted with a negative binomial with constant dispersion *r* for Ec-Hi2000-TruSeq(d) and Ec-Hi4000-Nextera(e). While Ec-Hi2000-TruSeq shows a significant slope and nearly no y-intercept, Ec-Hi4000-Nextera shows the exact opposite.

The flanking bias (Fig. 3b) has 120 parameters: one for each nucleotide across each of thirty positions (10 bases before the fragment to 20 bases within). The flanking bias at the start and end of the fragment use the same parameters due to their similarity (Fig. S2), with the end reverse complemented to keep the meaning of positions relative to the fragment. Since the best way of combining biases from individual positions in the flank is *a priori* unknown, we tested the two simple options of a product and a sum (see Methods) on datasets fragmented using mechanical forces (Ec-Hi2000-TruSeq) and enzymes (Ec-Hi4000-Nextera). As seen in Figure 3c, the predictions from the summation model nicely fit to the observed count means, using 2 · 10^5^ sites per bin.

To properly account for duplications, the dispersion has two parameters:

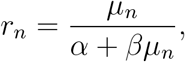

which leads to the following mean-variance relationship:

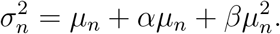

If we transform the dispersion equation to:

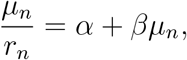

*α* is the y-intercept and *β* is the slope (see Figures 3d and e). This representation demonstrates that both parameters are needed to properly simulate all datasets and duplication types, even if single datasets need only one of the parameters (Fig. S3a).

The log-likelihoods are maximized with the limited-memory Broyden-Fletcher-Goldfarb-Shanno algorithm (L-BFGS) [36, 37] implemented in the NLopt package [38] due to its low number of required likelihood calculations. To further improve convergence and speed, ReSeq performs the fit in three steps (Fig. 3a). It starts by fitting a Poisson, which allows to analytically calculate the optimal GC bias for each bin. In a second step, the individual GC bins are connected using the cubic natural spline, while the spline’s knots are greedily adjusted to achieve the highest maximum likelihood. In the last step, ReSeq maximizes the likelihood for the full negative-binomial model with the knots fixed to the greedily-determined values. Afterwards, the normalized biases and dispersion from all converged reference sequences, fragment length combinations are merged using a median to be robust against outliers. Details on individual steps can be found in the Methods section.

To test the biases obtained by ReSeq, we use Bc-Hi4000-Nextera, Ec-Hi4000-Nextera and Rs-Hi4000-Nextera, which are all prepared using Nextera adapters. The flanking biases do not vary much in those three datasets (Fig. 3b), despite the different median GC content of the underlying genomes (35%, 51% and 69%). Furthermore, we clearly reproduce previous findings for Nextera adapters [23], where the biases between a nucleotide and its complement are very similar if mirrored around position 4 (e.g., A at 5 and T at 3). The GC biases for the three datasets are compared in Figure S4, where the spread between different fragment lengths highlights the lower confidence in biases based on fewer sites. Bc-Hi4000-Nextera and Ec-Hi4000-Nextera look very similar, except for low-confidence, GC-rich fragments. Rs-Hi4000-Nextera looks somewhat different, but its high-confidence region is poorly accessible by the other two datasets. The reduced occurrence of AT-rich fragments (AT dropout) described in the literature [23] is also observed with the exception of Rs-Hi4000-Nextera, which has little power to detect AT dropout due to its high median GC content. As another test, the biases were estimated on simulated, uniformly distributed data, which results in the expected flat profile (Fig. S5). The minor deviations for GC biases at the edges are due to low amounts of available sites. Overall, the above tests show that this part of the coverage model delivers accurate and consistent results.

After the other biases have been fitted, ReSeq estimates the reference-sequence and fragment-length biases. It starts by summing flanking and GC biases over all sites for selected combinations of fragment length and reference sequence *b_sum,ref,len_*. Additionally, the counts of mapped pairs for each reference sequence, *C_ref_*, and each fragment length, *C_len_*, are obtained (see Methods). The reference-sequence bias *b_ref_* and the fragment-length bias *b_len_* are then iteratively adjusted until convergence to follow the equations:

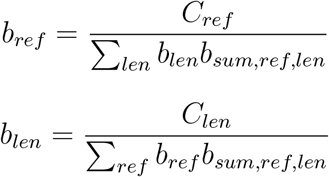

Figure 4 shows ReSeq’s improvements over the uniform distribution (ART and BEAR) or the sliding window approach (pIRS and NEAT), in terms of base coverage distributions and the number of duplicated read pairs.

**Figure 4:**
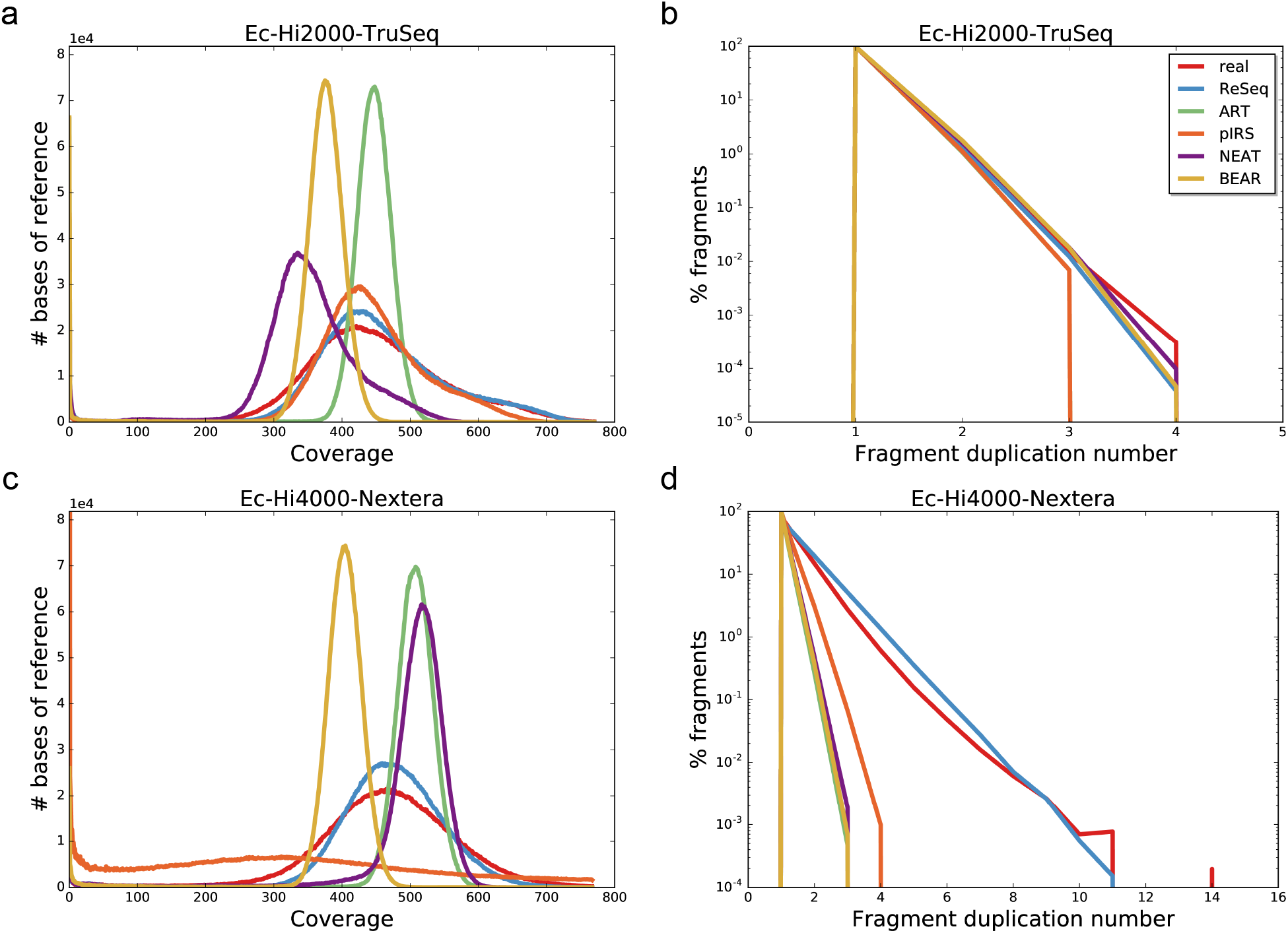
Coverage results. a,c) The base coverage distribution of the mapped reads. b,d) The number of duplications (i.e. read pairs mapped on the same strand to the same start and end position).

As a further test, we can use the technical replicates of Ec-Hi4000-Nextera (Ecoli1_L001) previously described. The correlation of the base coverage between Ecoli1_L001, Ecoli1_L002, Ecoli1_L003, Ecoli2_L001 and Ecoli3_L001 can be seen as a gold standard of how much the base coverage of simulations should be correlated to real data. We calculated the three pairwise Spearman correlations of replicates from the same lane and from the same library, respectively, and compared them to the correlations of three independent simulations based on Ec-Hi4000-Nextera. For the simulations, we calculated the pairwise Spearman correlations with themselves and the three correlations with Ec-Hi4000-Nextera. Table 3 lists the averages of the calculated coverage correlations. For each entry, one of the three corresponding correlations is plotted in Figure S6. The correlations highlight three major points. First, ReSeq strongly increases the correlation between simulation and real data because, despite its high variance, the whole coverage distribution follows the real trend, which is not visible for other simulators. Second, independent simulations from ReSeq approach the correlatedness of real data, but do not populate the low- and high-coverage regions present in real data. Only pIRS performs similarly, but with a correlation of 1, nearly all randomness seems to be removed. Third, despite the major improvements, the correlation between ReSeq and real data is still low and further improvements are possible.

**Table 3:** Average (pairwise) Spearman correlations of base coverage.

|  | Lane | Library | Real | Simulation |
| --- | --- | --- | --- | --- |
| Real | 0.91 | 0.92 | - | - |
| ReSeq | - | - | 0.23 | 0.81 |
| ART | - | - | 0.01 | 0.00 |
| pIRS | - | - | 0.12 | 1.00 |
| NEAT | - | - | 0.10 | 0.26 |
| BEAR | - | - | -0.00 | 0.05 |

Lane: Ec-Hi4000-Nextera (Ecoli1\_L001) and replicates Ecoli1\_L002 and Ecoli1\_L003. Library: Ec-Hi4000-Nextera (Ecoli1\_L001) and replicates Ecoli2\_L001 and Ecoli3\_L001. Real: Ec-Hi4000-Nextera and three simulations. Simulation: Three simulations of Ec-Hi4000-Nextera.

### Comparison of quality values and error rates

Here, we check whether the quality values and error rates show the typical patterns over the read length on all eight datasets (Table 2). Additionally, we test Ec-Hi2500-TruSeq-asm, where the simulation profiles are trained on a non-optimized assembly built from Ec-Hi2500-TruSeq that is highly fragmented and has many duplications (QUAST [39] report in supplement: 61992 contigs, N50 of 402 and covering 98.9% of the *E. coli* reference with every covered reference base being represented, on average, 2.553 times in the assembly). Thus, comparing the results of Ec-Hi2500-TruSeq-asm and Ec-Hi2500-TruSeq gives insight on how well simulators can build profiles from a fragmented assembly (e.g. by filtering). In contrast, using the assembly as simulation template would not allow similar insight, because a template can only be improved by changing it, which affects all simulators equally. Therefore, we continue using the *E. coli* reference as simulation template.

In general, the simulated mean quality values by position resemble those of real data (Fig. S7), except for BEAR, which produces reads of varying length and has for most datasets and positions, average quality scores 5 to 10 Phred units lower than the real data. A likely explanation for the strong deviation is BEAR’s design choice to replace quality values at error positions with qualities generated by the error model instead of having error rates depend on the quality values. For ART, we observe quality values consistently one Phred unit higher than the real data (Fig. S7b-e,g-i). For pIRS and ReSeq, deviations appear with increased position (Fig. S7d,g-i), which is likely caused by training the quality values only on mapped reads. Moreover, in Rs-Hi4000-Nextera, ReSeq experiences a particularly strong deviation in the second read’s quality (Fig. S8d), which is not present in the first read (Fig. S8c). It is also worth mentioning that pIRS uses the same quality distribution for first and second reads, despite the clear differences in real data (Fig. S8). Finally, the mean quality values for Ec-Hi2500-TruSeq-asm and Ec-Hi2500-TruSeq are the same, which is expected, because the qualities are independent of the reference.

The mean error rates by position are also well reproduced in the simulations (Fig. S9), except for BEAR, since it uses DRISEE for training where increased error rates have already been observed [40]. The applied exponential regression seems to amplify the increased error rates for higher positions (Fig. S9d,e,g). At-HiX-TruSeq, Mm-HiX-Unknown and Hs-HiX-TruSeq (Fig. S9g-i) are outside of BEAR’s defined metagenomic use-case, but this does not generally affect performance. On another note, Ec-Hi2000-TruSeq and Ec-Hi2500-TruSeq highlight the weakness of the hard-coded error models used by ART and NEAT (Fig. S9a-c). Such models require well-calibrated quality values (ART) or identical calibrations to their training set (NEAT). Especially in Ec-Hi2000-TruSeq, the quality values are not well-calibrated: while the Phred quality score of 2 predicts a 63% error rate, the observed rate after mapping is only 15%. Therefore, ART has strongly inflated error rates in this dataset, which could be solved by recalibrating the quality values, but then the quality values simulated from ART would be as inaccurate as the error rate is. Finally, ART’s and NEAT’s error rates drastically increase for Ec-Hi2500-TruSeq-asm compared to Ec-Hi2500-TruSeq-asm. However, this seems to be in large part an artifact from the reference-free error estimation, because increasing the coverage reverts the changes completely for ART and mostly for NEAT (Fig. S10).

### Comparison of 51-mer spectrum

A good high-level summary statistic to represent a dataset of genomic reads is the k-mer spectrum, which shows the systematic properties of errors (exponential decrease at low frequencies) and the coverage distribution (shape and position of peaks). Figure 5 (linear scale) and Figure S11 (log scale) display the 51-mer spectra of the datasets and their simulations. The first row (Fig. 5a-c) is based on high-coverage *E. coli* datasets sequenced on the older HiSeq 2000 and 2500 with TruSeq adapters and includes the simulations trained on the fragmented assembly instead of the reference (Fig. 5c). The second row’s bacteria datasets (Fig. 5d-f) are all produced in a single HiSeq 4000 run using Nextera adapters (enzyme fragmentation). They are ordered from left to right by coverage ranging from high (508x) to very high (2901x) and by genome complexity: single sequence (*E. coli*), two sequences where one is not present in the data (*B. cereus*) and multiple sequences including plasmids of varying abundance (*R. sphaeroides*). The plasmids cause multiple smaller peaks in the 51-mer spectrum with lower frequencies than the main peak (Fig. S11f). Additionally, especially in Bc-Hi4000-Nextera (Fig. S11e), we observe variants in the bacteria populations that increase the 51-mer counts between the systematic errors and the signal peak. Although NEAT and ReSeq accept Vcf files, which allows non-diploid variants to be specified, we do not include them in the simulation. In the third row, we have low-coverage (31x-43x), diploid datasets sequenced on HiSeq X Ten machines with TruSeq or related adapters. Here, we do include the variants in the simulations, which results in a second peak in the 51-mer spectrum at half the frequency of the main peak (Fig. 5h,i). Furthermore, the three genomes contain sequences with differences in abundance: in At-HiX-TruSeq (Fig. S11g), we observe a second peak at higher frequencies stemming from the mitochondria; in Mm-HiX-Unknown (Fig. 5h), complex changes in the NIH3T3 cell line [41], including copy-number variations, broaden the signal peak; and, in Hs-HiX-TruSeq (Fig. 5i), the X and Y-chromosomes strengthen the heterozygous peak at half-the coverage of the main peak.

**Figure 5:**
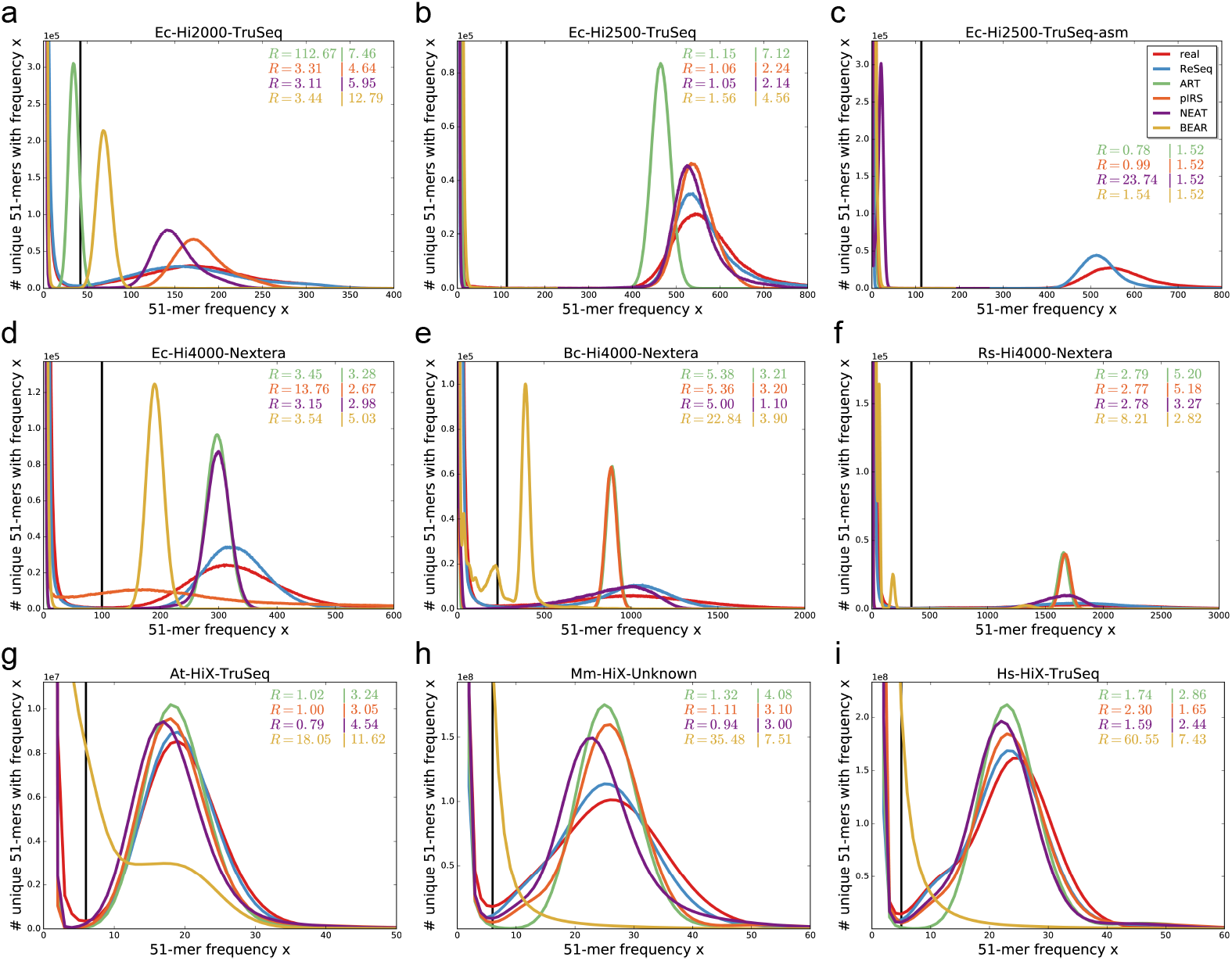
51-mer spectra of real and simulated data. The shape and position of peaks reflects the coverage distribution, while the exponential decrease at low frequencies is defined by systematic errors. The black lines show the minimum between the exponential decrease and signal peak for real data. The first value of *R* gives the sum of relative deviations for frequencies (x) up to the minimum. The second value of *R* gives the sum of absolute deviations for frequencies larger than the minimum. Both values are stated relative to ReSeq. When the signal peak of the simulation does not exist or starts before the minimum in real data the values lose their interpretability. This happens for ART (a,c), pIRS (c,d), NEAT (c) and BEAR (b,c,f-i).

BEAR does not produce a signal peak that matches the real signal’s shape or position in any of the tested datasets. ART is consistently underperforming, for the coverage distribution as well as for the systematic errors. Its error rate inflation for Ec-Hi2000-TruSeq is causing the strong peak shift in the 51-mer spectrum (Fig. 5a). Notably, pIRS struggles with datasets with Nextera adapters: for Ec-Hi4000-Nextera, it results in a rather flat peak and an overabundance of high-frequency 51-mers (Fig. 5d), while it maintains a uniform coverage distribution for the other Nextera datasets (Fig. 5e-f).

ReSeq compares favourably for the coverage distributions of all datasets (Fig. 5) and for the systematic errors of most datasets (Fig. 5a-f, S11a-f). Notably, ReSeq reproduces the coverage peak better on data with TruSeq adapters (Fig. 5a,b,g-i), compared to Nextera adapters (Fig. 5d-f). Furthermore, for all three HiSeq X Ten samples (Fig. 5g-i), ReSeq slightly underestimates the coverage and, more importantly, misses the systematic nature of most errors. The coverage could be manually corrected by specifying a parameter, but the missing systematic errors are caused by the low coverage and are only solvable with more data. Especially in At-HiX-TruSeq (Fig. 5g, S11g), the effects of low coverage are visible.

In Ec-Hi2500-TruSeq-asm (Fig. 5c), the median coverage provided as coverage parameter to ART, pIRS and NEAT is not a good estimator for the real coverage, because the duplicated regions and the many very short hard-to-map contigs in the assembly reduce the median coverage from 1011 to 13. Therefore, ART and pIRS do not have an expected peak, whereas NEAT’s peak is at a very low coverage. ReSeq extracts the coverage itself and is therefore not directly affected. BEAR requires the number of reads and is therefore also not affected, but still does not show an appropriate peak in the k-mer profile. In Figure S12, the median coverage estimated from the mapping to the reference is provided to all simulators except BEAR. For ART, the improved coverage removes nearly all performance losses compared to Ec-Hi2500-TruSeq, which is not surprising, since only InDel rates and the fragmentlength parameters are taken from the assembly. Even though its performance is still low, ART improves relative to pIRS and NEAT, because these two do not cope well with the fragmented assembly and produce rather flat and broad peaks. ReSeq’s coverage model also suffers from using the assembly instead of the reference, but can still maintain a general resemblance to the real data.

### Comparison of assembly continuity

To check whether the improved representation in the k-mer spectrum has consequences for applications, we ran the sga preqc module on all datasets and their simulations (Fig. 6). The preqc module estimates the N50 (length of the shortest contig still necessary to cover 50% of the genome) of short-read assemblies for different k-mer lengths. Due to crashes, Ec-Hi2500-TruSeq-asm is missing pIRS and Rs-Hi4000-Nextera is missing pIRS and ART.

**Figure 6:**
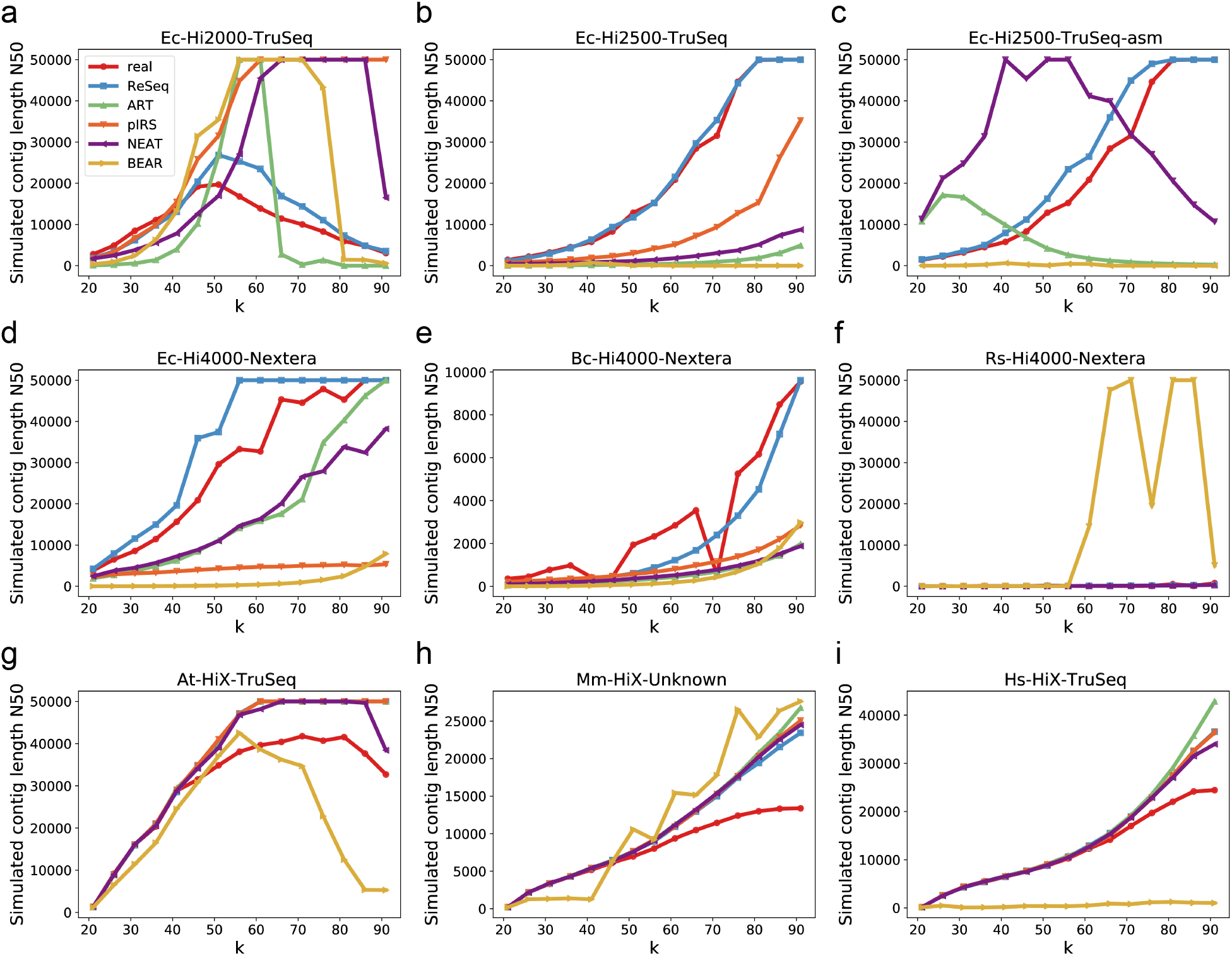
Simulated contig N50 for real and simulated data. c) preqc crashes for pIRS f) preqc crashes for pIRS and ART

For most datasets, ReSeq follows real data better than the other simulators (Fig. 6a-e), except for the low-coverage HiSeq X Ten datasets (Fig. 6g-i), where the missed systematic nature of errors and undetected diploid variants prevent ReSeq from delivering the same performance. For Rs-Hi4000-Nextera, BEAR deviates drastically from real data and other simulators (Fig. 6f). Figure S13 shows the same plot without BEAR, but due to fluctuations in the real N50 values, no strong conclusions can be drawn.

In Ec-Hi2500-TruSeq-asm (Fig. 6c), the coverage-parameter estimate from the assembly strongly affects the N50 values of ART, pIRS and NEAT. Providing the coverage estimate from the reference to all simulators except BEAR (Fig. S14), mostly reverts the drastic changes in N50. pIRS’s difference to real data only slightly increases, compared to using the reference for the complete training (Ec-Hi2500-TruSeq: Fig. 6b), while NEAT does not show the unexpected peak of N50 around *k* = 50 anymore, but does not restore the slight resemblance to the real data it had in Ec-Hi2500-TruSeq, and ART fully recovers the increase in N50 towards high *k* that has at least the same tendency as real data. Noticeably, ReSeq’s N50 values stay close to the real ones no matter how it was trained.

### Comparison of cross-species simulations

An important property of trained profiles is that not only can they be used to simulate the original dataset, but also datasets with alternative templates. To test training and simulation across species, we split six of the datasets into two groups with identical sequencer and fragmentation combinations. The datasets were simulated using the median coverage, template and variants from the simulated datasets; all other parameters and profiles were provided from another dataset within the group. The resulting 51-mer spectra are compared to those of the simulated datasets in Figure 7 (linear scale) and Figure S15 (log scale). In the HiSeq 4000 group with enzyme fragmentation, we used the profiles from Ec-Hi4000-Nextera to simulate Bc-Hi4000-Nextera and Rs-Hi4000-Nextera and the profiles from Rs-Hi4000-Nextera to simulate Ec-Hi4000-Nextera. The results highlight the difficulty of applying profiles across genomes with very different GC content (Fig. 7a-c). In the HiSeq X Ten group with mechanical fragmentation and genomes with similar GC content (Fig. 7d-f), we used the profiles from Mm-HiX-Unknown to simulate At-HiX-TruSeq and Hs-HiX-TruSeq and the profiles from Hs-HiX-TruSeq to simulate Mm-HiX-Unknown.

**Figure 7:**
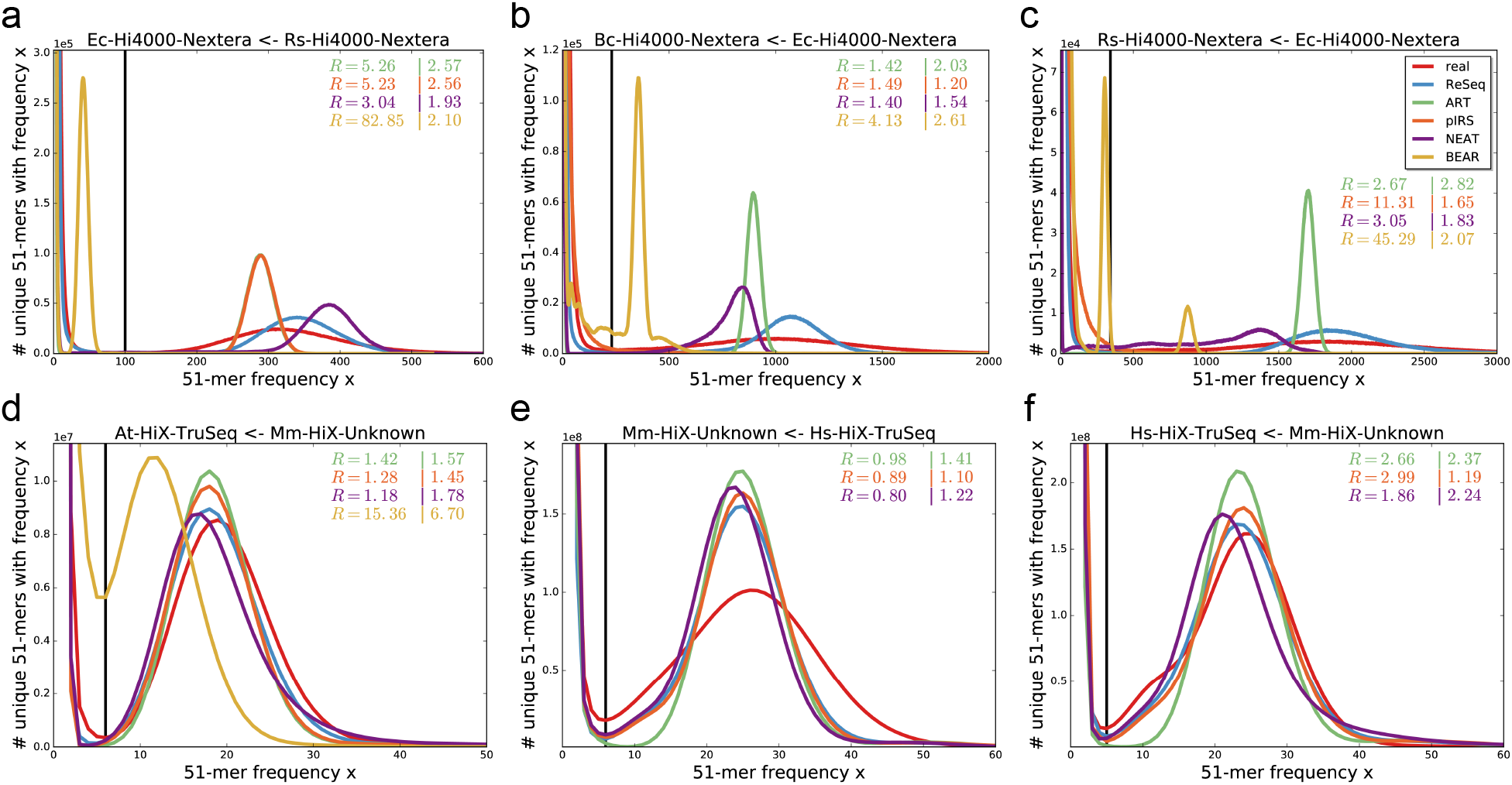
51-mer spectra of real data and cross-species simulations. The simulation profiles are trained on a different training dataset. The pairings are given in the plot titles in the following notation: “simulated dataset ← training dataset”. The shape and position of peaks reflects the coverage distribution, while the exponential decrease at low frequencies is defined by systematic errors. The black lines show the minimum between the exponential decrease and signal peak for real data. The first value of *R* gives the sum of relative deviations for frequencies (x) up to the minimum. The second value of *R* gives the sum of absolute deviations for frequencies larger than the minimum. Both values are stated relative to ReSeq. When the signal peak of the simulation does not exist or starts before the minimum in real data the values lose their interpretability. This happens for BEAR in panel a and c. BEAR is omitted from panel e and f due to its long runtime.

Similar to the case of simulating the original datasets (Fig. 5), BEAR does not create a signal peak that matches the real signal’s shape or position in any of the Nextera datasets (Fig. 7). ART is unaffected by the profile change, but still shows low performance due to the uniformly distributed coverage. pIRS’s performance on cross-species simulations is hard to judge on the Nextera datasets, because pIRS already underperforms when reproducing the original Nextera datasets. For example, pIRS on Ec-Hi4000-Nextera results in a rather flat peak (Fig. 5d) and simulating other datasets from its profile results in no visible peak (Fig. 7b,c). However, for TruSeq adapters and genomes with similar median GC content, pIRS seems to be unaffected by using profiles trained on a different species (Fig. 5g-i, 7d-f)). This is not the case for NEAT, where in the HiSeq 4000 group, minor peaks appear on the low-frequency side of the main peak (Fig. 7c and S15a,c) and in the HiSeq X Ten group, an asymmetric signal peak with a pronounced tail is visible in the simulations based on the Mm-HiX-Unknown profiles (Fig. 7d,f). This is likely caused by other sources of bias being incorporated into the GC bias. ReSeq is missing the abundance information for the reference sequences and therefore does not correctly simulate the plasmids in Rs-Hi4000-Nextera (Fig. 7c), the mitochondrial genome in At-HiX-TruSeq (Fig. S15d), the copy-number variations in Mm-HiX-Unknown (Fig. 7e) and the X and Y chromosome in Hs-HiX-TruSeq (Fig. 7f). This demonstrates how ReSeq nicely separates the individual biases and produces results (Fig. 7) resembling the original datasets (Fig. 5). On Mm-HiX-Unknown, we demonstrate that given the abundance information, ReSeq again produces a coverage peak as good as the one from the simulation trained on the original dataset (Fig. S16 and 5h).

### Comparison of computation requirements

Besides better representation of real data, computational requirements are also important. Figure 8 shows the total CPU time, the elapsed time and the maximum amount of memory used for the training and simulation steps of each simulator for the eight datasets. Reseq lies on the high side of CPU time for training and simulation (Fig. 8a,d). Noticeably, parts of ReSeq’s CPU requirements scale with genome size instead of number of reads, which leads to worse performance in lowly-covered genomes.

**Figure 8:**
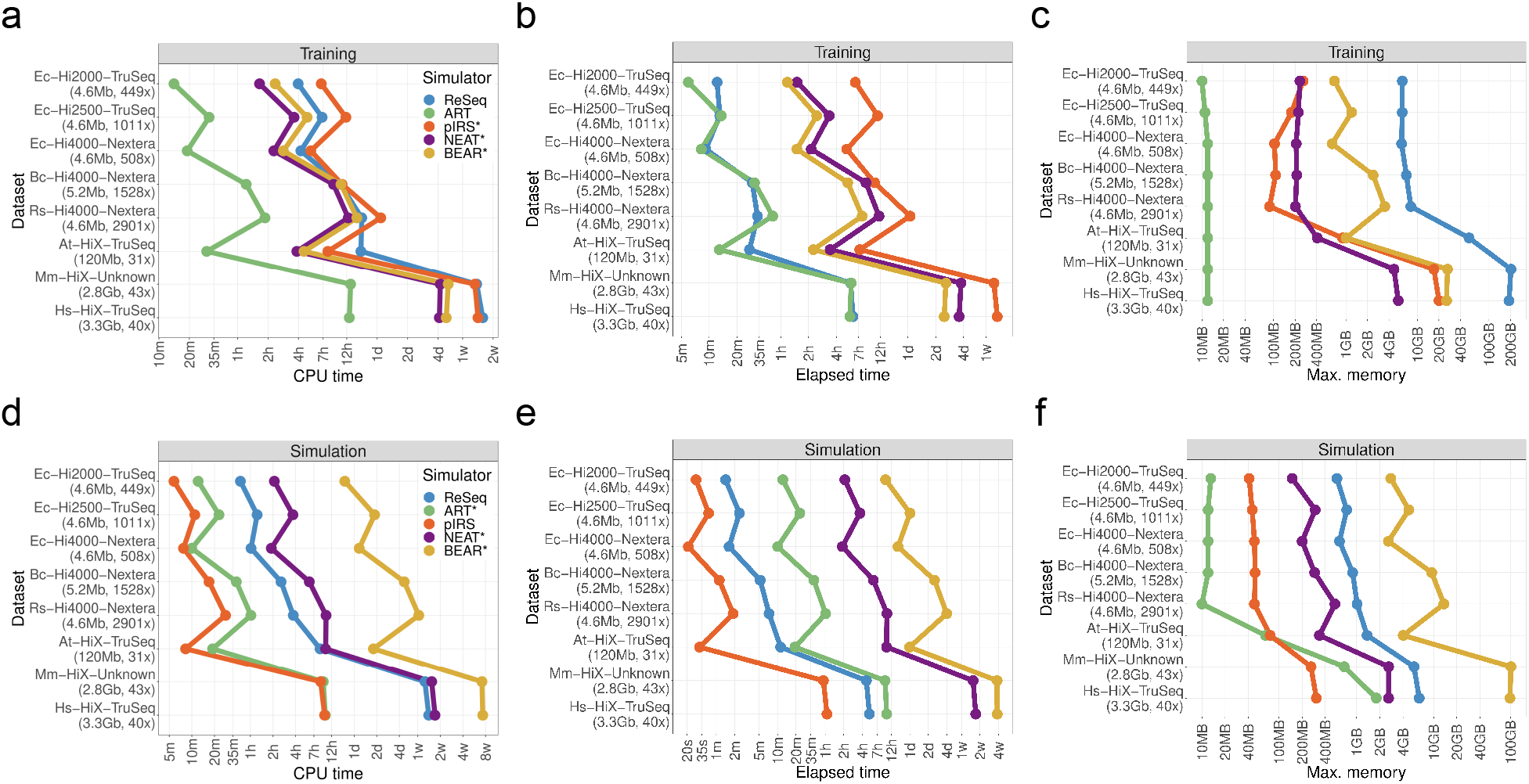
Speed and memory benchmark of simulation process. The training part needs to be performed only once to generate the profiles. The simulation part needs to be done for every synthetic dataset created. The simulators labeled with a * are single threads per task, but often include multiple task being run in parallel. All datasets have the reference size and the coverage specified.

Due to ReSeq’s effective parallelization, its elapsed times are low for this benchmark with 48 virtual CPUs (Fig. 8b,e). In contrast, the single-threaded processes implemented in perl or python have strikingly high elapsed times. This is best visible in Hs-HiX-TruSeq and applies to the training of pIRS (over a week), NEAT (several days) and BEAR (half a week) as well as the simulation of NEAT (close to two weeks) and BEAR (several weeks). For the simulation, NEAT provides a tool to split the template and later merge the reads, which can be used for manual parallelization to reduce the elapsed time.

Memory requirements remain mostly below 10GB for training (Fig. 8c,f). The exceptions are pIRS needing around 20GB to train on human-sized genomes and ReSeq requiring around 200GB for training on these datasets. However, ReSeq’s memory consumption during training scales almost linearly with the amount of CPUs used (Fig. S17). Therefore, the memory requirements can be reduced to 11GB by using a single thread. For simulations, the memory requirements stay also below 10GB, except for BEAR that needs close to 14GB for Rs-Hi4000-Nextera and around 100GB for Mm-HiX-Unknown and Hs-HiX-TruSeq. In typical systems with 2-8GB per core, BEAR’s high-memory requirements may block many CPUs additional to the 2 used ones, which could lead to effective CPU times much higher than the values in Figure 8d.

### Example: genuine comparison of short-read mappers

After comparing ReSeq to other simulators, we demonstrate its use in an example of a simulation study, where the performance of two popular mapping algorithms, bwa and bowtie2, is compared. For the simulation, we trained ReSeq either on bowtie2 or bwa mappings to verify that performance is not biased by the mapper used for training.

As a first quality check, we compare mapping statistics between simulated and real data for three datasets (Fig. 9a-c). In real data, the number of unmapped pairs and single reads is higher compared to the simulation, which is most pronounced in Ec-Hi2500-TruSeq (Fig. 9b), where the mapping of some reads is prevented by deviations between the reference genome and the true genome underlying the data. In contrast, simulated reads are drawn directly from the reference template and thus do not exhibit mapping issues caused by genome deviations. Furthermore, the bwa-based Ec-Hi4000-Nextera simulation (Fig. 9c) compared to the bowtie2-based simulation and real data has less unmapped reads for bowtie2 and less soft-trimming, which highlights that the bwa-based simulation underestimates the adapter content of the reads. As a result, the bowtie2-based simulation is to be trusted more for this dataset. In Ec-Hi2000-TruSeq, the rate of soft clipping drops for the bowtie2-based simulation (Fig. 9a), as it did for the bwa-based simulation of Ec-Hi4000-Nextera (Fig. 9c), but the number of unmapped reads does not drop accordingly and thereby makes adapters an unlikely cause. Finally, the number of insertions and deletions varies in several cases (Fig. 9a-c), but for the general comparison of mappers here, the differences are too low to influence the result.

**Figure 9:**
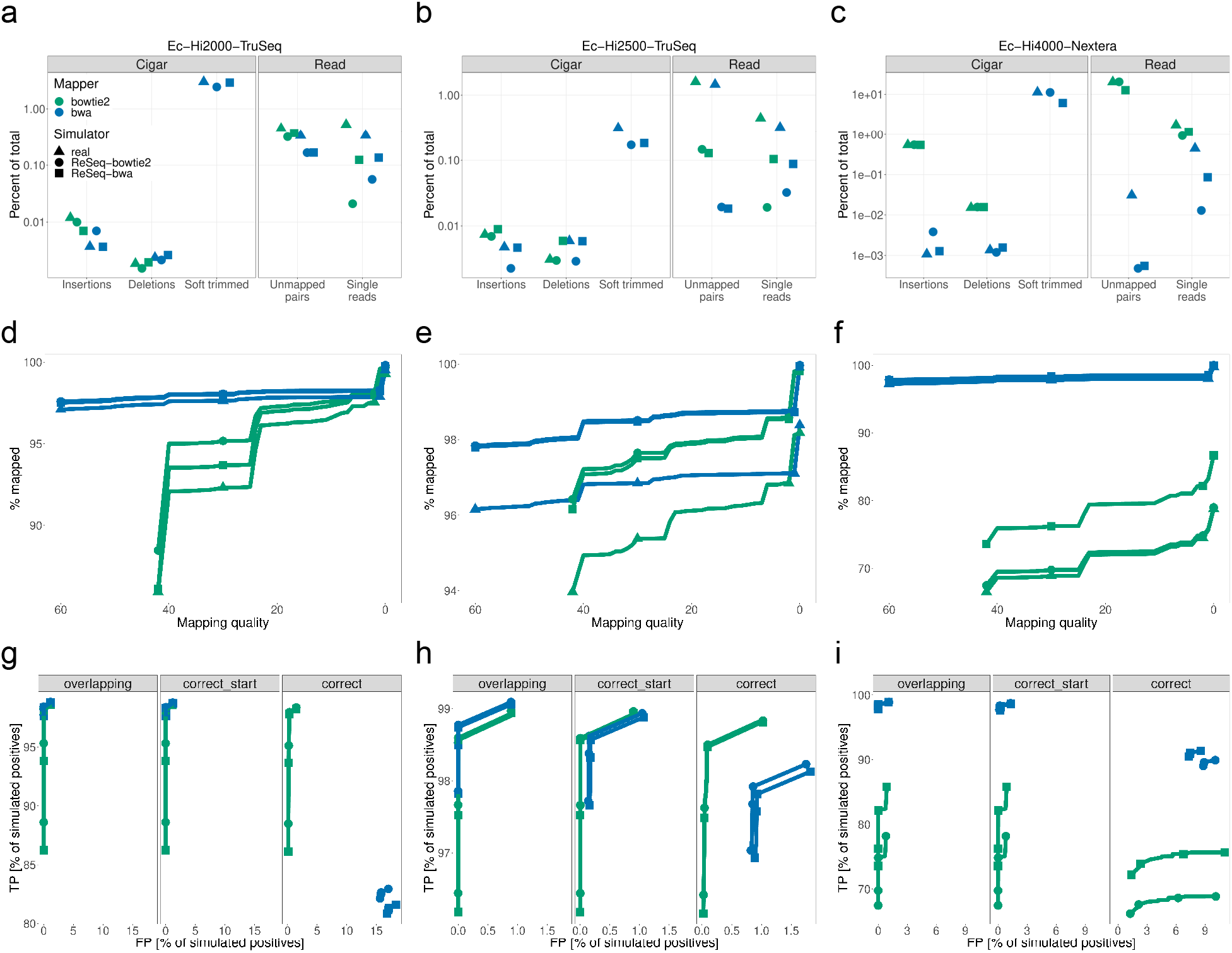
Comparison between bwa and bowtie2 on *E. coli*. Each dataset has simulations trained on mappings from bwa and bowtie2. a-c) Statistics of mapping outcomes. Bowtie2 does not trim and therefore has no soft trimmed bases. d-f) Mapping quality distribution. Mapping qualities are cumulative, i.e. all mapping qualities at the given score or higher. Markers are only shown for mapping qualities 0,2,30,42 for bowtie and 0,l,30,60 for bwa. g-i) Mapping accuracy for all mapping quality thresholds. Markers are only shown for mapping qualities 0,2,30,42 for bowtie and 0,l,30,60 for bwa. Positives are mapped reads, which fulfill the correctness criteria (TP) or do not (FP). Overlapping: True and mapped positions overlap independent of strand. Correct_start: Perfect match of start position and strand. Correct: Perfect match of start and end positions and strand. a,d,g) Ec-Hi2000-TruSeq b,e,h) Ec-Hi2500-TruSeq c,f,i) Ec-Hi4000-Nextera

As a next step, we compare the mapping-quality distributions between simulated and real data (Fig. 9d-f) and observe that the mapping rates increase in mostly identical steps, when mapping quality requirements are lowered. The overall shifts in mapping rates are an alternative representation of the different percentages of unmapped reads discussed in the previous paragraph. A prominent feature not explained by the general shifts can be seen in Ec-Hi2000-TruSeq (Fig. 9d), where the difference in mapping rate between the two simulations is much more pronounced for high-quality mappings compared to low-quality mappings. In light of the observed decrease in soft clipped bases (Fig. 9a), the mapping rate differences can be explained by a lower number of simulated reads with many errors (> 10) for the bowtie2-based simulation compared to the bwa-based (Fig. S18). We observe that bwa handles dense errors by clipping the reads, while bowtie2 reduces the mapping quality. Since an increased soft clipping in real data can also have other causes, such as the deviations from the reference, it is unclear which simulation better represents the real error rates.

Finally, we can use the truth from the simulations to calculate true positives (TP) and false positives (FP) of read mapping (Fig. 9g-i). In Ec-Hi4000-Nextera (Fig. 9i), the TPs strongly differ between simulations due to the missing adapters in the bwa-based simulation. Thus, we use the bowtie2-based simulation in this case. In Ec-Hi2000-TruSeq (Fig. 9g), we also observe strong differences in TPs between simulations, but here they are due to the different amount of reads with many errors (Fig. S18).

Overall, we observe that in the case of strong adapter presence, bwa is the better choice due to its soft trimming capability (Fig. 9i), although bowtie2 allows better control over FPs with adequate mapping quality thresholds in case both read ends are required to map perfectly. Ec-Hi4000-Nextera also highlights how important adapter simulation is to get an adequate mapping comparison (Fig. S19). In the datasets with lower adapter content (Fig. 9g-h), the recommended mapper depends on the downstream application. If perfect mapping is required, bowtie2 is the better choice, since bwa often trims too much. If only overlaps with the true position are needed, for example for counting in intervals, bwa slightly outperforms bowtie2. If having a correct start position is sufficient, the mappers perform approximately the same with a minimal preference for bwa if reads with many errors are frequent and a minimal preference for bowtie2 otherwise.

## Discussion

ReSeq’s advancements in faithfully reproducing real data can improve many benchmark studies. Short-read error-correction methods could previously not been adequately tested on simulations, because all methods scored nearly perfectly [16]. Many error-correction methods are based on splitting erroneous and correct k-mers at the spectrum’s minimum that occurs between the signal peak and the exponential decrease at low k-mer frequencies. ReSeq improves the shape of the exponential decrease by including systematic errors and broadens the signal peak with the extended coverage model. The similarity to real k-mer spectra will drastically increase the difficulty for error-correction methods. Compared to real data, the simulation has the advantage of knowing the true sequence for every read and does not require to derive it by mapping to a reference genome, which can create biases due to repeat regions, ploidy and deviations between data and reference. However, simulations do not have the full complexity of real data and we recommend a combined evaluation that compares the results from simulations and real data, because it has the potential to hint at inconsistencies caused by deriving the ground truth or imperfectly mimicking real data and to roughly estimate remaining uncertainty in the benchmark.

The situation for assembler benchmarks is similar. We have shown above that ReSeq simulates datasets with realistic N50 values and, in contrast to real data, we can be certain that the dataset is perfectly represented by the reference used to evaluate the correctness of the assembly.

In the case of mapping benchmarks, real data with ground truth is incredibly laborious to produce and in comparison to other simulators, ReSeq includes adapters and reads with a large number of errors, which increases the faithfulness of the results. Since variant calling is performed on mapped reads, these improvements will also translate to variant-caller benchmarks. Additionally, the inclusion of systematic errors adds one source of false positive variant calls that is not present for other simulators.

Despite all the advancements of ReSeq, no simulation is perfect and neither are the ground truth estimates from real data. Therefore, it is very important to assess the scope of a benchmark, clarifying under which circumstances the study results are relevant for real-life applications. The scope of simulations can be tested and defined with comparisons between real and simulated data on statistics accessible for both (e.g. mapping rates). Our example study, comparing short-read mappers, is limited in three ways: i) mapped reads do not vary from the reference; ii) the reference is completely assembled and does not have many repeat regions; and, iii) InDel simulations might need improvements for InDel specific benchmarks. In a more complete study, the scope could be extended by using references with different levels of continuity, repetitiveness and divergence from the simulated datasets. If InDels are of interest, a more detailed investigation of the differences between real and simulated data would be needed. For example, examining the InDel distribution over the genome could highlight regions where the reference varies from the true genome of the real data. Larger variations can cause distortions in the mapping and induce false InDels into reads, which are then translated into the simulation profiles. An improved reference would reduce the number of false InDels and thereby increase concordance between simulations.

For optimal simulations with ReSeq, the reference genome and variant calls provided for training should be of high-quality and closely match the used reads. Homozygous variants are preferred directly in the training genome instead of the variant file. Furthermore, over 100-fold coverage is advised to capture systematic errors well. Since adapters are learned from the data, the provided reads should not be trimmed. This requires running the Illumina basecaller with deactivated adapter trimming, which is generally advised, because keeping raw data untrimmed allows to later modify the trimming step that can affect analysis results [42, 43]. Therefore, many but not all public datasets provide untrimmed data.

During the simulation step, the provided template and variants can be synthetic. However, care should be taken that they mimic real genomes in terms of repetitiveness and heterozygosity. Furthermore, the GC content of the template should not deviate strongly from the training reference since GC biases are only accurate in the trainable range.

The only drawback of ReSeq compared to the other simulators are the resource requirements. Due to the almost linear increase in required memory with used cores, this comes down to required CPU time. We optimized the code and replaced many time-consuming calculations with estimations, but further optimizations and estimations could be implemented.

The biggest potential gain in representing Illumina genomic sequencing data could be a predictive model for the systematic errors. This would adjust for differences of systematic-error rates across genomes and could increase sensitivity for lowly-covered genomes. In the absence of a predictive model, it is most important to improve low-coverage datasets by handling the strong dependence of systematic-error rate on the coverage. The dependence is based on the discrete nature of errors and leads in combination with unequal coverage over the genome to inaccuracies, which become exacerbated when higher coverages are simulated from low-coverage statistics.

The coverage model could be extended by taking into account interactions between bases for the flanking bias. Especially for enzyme cutting, the effect of a motif likely varies from the sum of the effects of each base. Also the effect of stretches of GC instead of only the overall GC content could be taken into account, as done for RNA [44].

Finally, additional data types could be supported. For targeted sequencing methods, ReSeq’s coverage model could be extended to capture and reproduce the coverage distribution on and around the targets, including off-target effects and target-dependent enrichments. Defining targets in a general form would allow to reuse the extension to simulate exons and introns for RNAseq. However, this requires overlapping targets, which could be handled with equivalence classes. For methods such as 10x Genomics, barcodes, empty droplets, doublets and index misassignment, could all be added. Combining barcodes and a coverage-model extension for overlapping targets, would allow to simulate single-cell RNAseq datasets at the read level. Metagenomics and allele-specific methylation are already supported, but would profit from a separate simulator comparison.

## Conclusion

ReSeq improves the faithfulness of simulated data for all tested datasets. To achieve this, we solved three major challenges. First, we developed a coverage model that can be trained on complete large genomes. Second, we included systematic errors into the simulation. Third, we efficiently represented the important statistics, such that memory requirements remain constrained and the parameters can still be learned from a single real dataset.

Furthermore, ReSeq provides an easy-to-use training of all required models. No manual choice of parameters is needed, which simplifies usage over a wide range of genomes, Illumina machines and DNA preparations. ReSeq is also more robust to fragmented references during the profile generation compared to pIRS and NEAT.

The simulation of eight diverse datasets showed the importance of choosing good training data that fit the desired simulation in terms of genomic GC distribution, preprocessing (PCR, fragmentation) and sequencing machine. Therefore, hard-coded profiles should be avoided.

ReSeq and all of its code are available [45].

## Methods

### Adapter detection

ReSeq determines adapters automatically from the 10-mers of all unmapped parts of the reads, which in case of unmapped reads are the whole read or otherwise, the read segments that exceed the fragment length. ReSeq collects the abundance as starting 10-mer and the total abundance of 10-mers in the unmapped parts. It does so separately for the first and second reads. If the starting 10-mer appears a second time within the unmapped part, the read is rejected and not taken into account for the 10-mer analysis, therefore removing low-complexity regions.

To find the adapters, the starting 10-mer with the highest abundance is used as a seed and extended separately in both directions with the 10-mer that has the highest total abundance of the four possible choices (i.e. added nucleotides). The extension stops when the most abundant 10-mer has no longer an abundance five times higher than the next highest abundance or the 10-mer is already present in the adapter. ReSeq generally filters 10-mers consisting of a single repeated nucleotide and trims the poly-A tail at the end of the adapters. Figure S20 demonstrates the procedure for the first (of the paired-end) reads of Ec-Hi2000-TruSeq.

The automatic detection might fail in case of low adapter presence and does not handle more than one adapter for the first or the second read. For those cases, adapters can be specified. The adapter sequences are provided in FASTA format and a corresponding matrix defines the valid adapter pairings for first and second read. For the common Nextera XT v2 and single TruSeq adapters [31], the sequences and matrix are provided with ReSeq. After the adapters have been determined or specified, ReSeq uses the code from skewer [46] to find adapters in unmapped and low quality reads. The seqan library [47] is used for storing and reading from disk and general sequence handling.

### Handling of Ns

For the statistics calculation ReSeq ignores positions containing an N in the reference. At the beginning of the simulation, ReSeq replaces regions of 1 to 99 consecutive Ns in the template with random bases. Regions with at least a hundred Ns are replaced with a repeated 4-mer made up from the two bases following and the two bases preceding the region. This prevents replacing the non-assembled parts of the genome with easy-to-assemble random sequences. To use the same bases for the randomly replaced Ns over multiple simulation runs, the replacement can be stored in a FASTA file previous to the simulation.

### Iterative proportional fitting

To save memory and to reduce the size of the required inputs for the statistics calculation, ReSeq stores only the two-dimensional margins for each statistic instead of the whole matrix. Afterwards, the iterative proportional fitting (ipf) procedure [32] is applied, because no formula exists that can combine the full set of two-dimensional margins into a single matrix. Formulas can only satisfy at most a number of two-dimensional margins equal to the dimension of the matrix [48]. The ipf procedure adjusts the values in the matrix such that they fulfill one given margin at a time, looping over all given margins. For each positions *ij* in the currently handled margin *s* all contributing matrix elements *m_ijkln_* are adjusted in the following way:

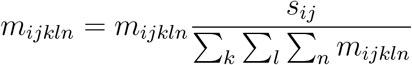

Repeating those adjustments many times for all margins has been shown to converge towards the solution [32].

In its original form, the iterative proportional fitting stores the full matrix, which potentially requires TBs of memory. However, the fitting can be nicely combined with a data reduction technique described by Fienberg [33]. There, a matrix element is stored as the sum of average differences from the lower dimensional average. For a two-dimensional matrix, the full description of element *m_ij_* is:

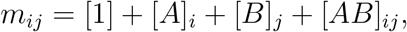

where [1] is the average value of the whole matrix, [*A*]_*i*_ is the average value of row *i* minus the overall average [1], [*B*]_*j*_ is the average value of column *j* minus the overall average [1], and [*AB*]_*ij*_ is the difference of the element *ij* from the average predicted by [1], [*A*]_*i*_ and [*B*]_*j*_. The full representation does not reduce memory, since the matrix [*AB*] is of the same size as the original matrix *m*. However, for our four and five-dimensional matrices, truncating the representation at two dimensions drastically reduces the storage requirements. This truncation corresponds to the two-dimensional margins used for the iterative proportional fitting. ReSeq includes the overall average [1] and the one-dimensional vectors [*A*]_*i*_, [*B*]_*j*_,…, [*E*]_*n*_ into the two-dimensional matrices. Furthermore, the exponential function is applied to guarantee positive numbers. Thereby, the representation of a five dimensional matrix is reduced to the following combination of ten two-dimensional matrices:

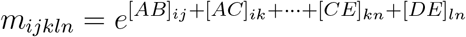

### Stored probabilities and conditions

Figure 1 lists all dimensions of the six statistics matrices. The matrices are used to query the probabilities along the first dimension, conditional on all other dimensions. The probabilities for the five possible systematic error tendencies (random error and the four nucleotides) depend on the position in a systematic-error region (see *Distinguishing systematic and random errors* below), the GC content before the given base, the systematic error rate at the first base in an error region, the reference base, the previous reference base and the most frequent nucleotide within the last five bases (dominant reference base). If multiple nucleotides have the same frequency within the last five bases, ReSeq choses the one that is the closest to the current base as dominant reference base. The GC content is calculated in a window from half a read length before the base up to the base. The error-region position is cut into bins of ten bases with the first bin going from position one to ten. The position of zero means either the position where the error region starts, or a position outside of error regions (i.e. no dependence on previous systematic errors). The probabilities for the systematic error rate at each template position depend on the same variables, except that the previous and dominant reference base are replaced by the dominant error.

The sequence quality is based on the GC content of the read, its mean systematic error rate, fragment length, tile and whether it is the first or second read(template segment). Fragment lengths are binned into bins of ten for this purpose.

The four insertion probabilities (one for each nucleotide) and the deletion probability depend on the position in the current insertion or deletion, the position in the read, the GC content of the error-free read and the last regular base call. ReSeq also models whether the read position directly before the potential insertion or deletion is itself an insertion or deletion (InDel before), since this means that the InDel will be an extension of the current InDel.

Quality values are assigned based on the drawn sequence quality, previous base quality, read position, systematic error rate, template segment, tile and reference base.

The regular base calls are determined from the base quality, read position, number of errors already in the read, systematic error rate, template segment, tile, reference base and dominant error.

### Distinguishing systematic and random errors

To formally test whether an error is systematic or random, ReSeq determines the most abundant substitution error and its count for every position (reference base, strand). Then the probability is calculated that at least this many errors are observed under the null hypothesis (a P-value). We assume random errors (the null hypothesis) to have equal probabilities for every base in non-overlapping windows of the genome and to convert bases with equal probability to the three possible erroneous nucleotides. Under this simplification, the number of conversions into a specific nucleotide at a given position follows a Poisson distribution with a mean *μ* equal to the coverage *N_c_* at the given position times the median error rate *r_e_* divided by three: 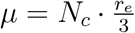. The median error rate is calculated in bins of 10^4^ reference bases (i.e. 2 · 10^4^ individual positions due to the separation of the two strands).

After the P-value calculation, we apply the Benjamini-Hochberg procedure [49] and call errors systematic if their adjusted P-value is less than 0.05. If both strands are reported as systematic, there is potentially a variant at this reference base. Therefore, reference bases are ignored for all statistics, if at this position the most abundant error of the forward strand matches the complemented most abundant error of the reverse strand. When ReSeq detects a non-variant systematic error, the position on this strand gets assigned the most abundant substitution error as systematic error tendency and its frequency as error rate. Furthermore, the position is set as the start of an systematic-error region with a length of one read length. The error region is defined by the error rate of the systematic error starting it. If another systematic error is detected within the error region it is considered to be caused from the same trigger sequence as the first systematic error, except if it has a higher error rate in which case the earlier error region stops and a new error region begins.

### Removing low-quality sites

To filter low-quality sites, the start position of reads on the forward strand and the end position of reads on the reverse strand are considered. Start positions of low-quality reads (mapping quality < 10) are connected if less than 10 other reads start between them. Unconnected positions are likely caused by the read and not the site and thus are treated as high-quality. Connected start positions instead highlight low-quality regions that in addition to all positions between them, are ignored for the coverage model. Similarly, we ignore sites ending on or between connected end positions of low-quality reads.

### Bias fit selection

The GC and flanking bias estimation uses only the longest reference sequences covering 80% of the genome (L80). ReSeq performs fits for some of the thirty most abundant fragment lengths in each of those sequences. How many of the thirty most abundant fragment lengths are fitted depends on the number of reference sequences in the L80. For a single reference sequence, all thirty are fitted. For twenty and more sequences, only five fits per sequence are equally distributed over the thirty most abundant fragment lengths. For 2-20 sequences, the number of fits per reference sequence is gradually reduced. For example, the single sequence of *E. coli* results in thirty fits, while the four (of seven) sequences of *A. thaliana* in the L80, result in 4 · 15 = 60 fits.

### Flanking and GC bias formulas

The mean of the negative binomial, as described in the *Coverage model* section, has the following form:

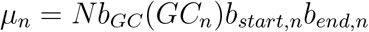

where *n* is the site in the genome, *N* is the normalization factor and the GC or flanking biases are represented by the *b* parameters. For the flanking bias, we tested two choices to combine the individual positions: sum and product. If we choose the product of the positions *p*, the flanking bias at the start results in:

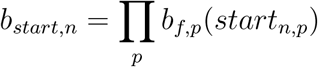

In contrast, the sum over the positions can become negative, so we apply twice the inverse logit. Since the center of the logit is approximately linear, this should keep the summation behaviour:

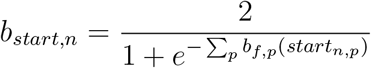

For the GC bias, we use an exponential to prevent negative values and thus keep the canonical link function:

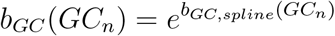

Applying the inverse logit, similar to the flanking bias, improves the speed or convergence rate for some datasets, but generally performs worse (Fig. S21).

### GC knot adjustment

In the second step of the GC and flanking bias fit for the *Coverage model*, the GC bias is fitted with a spline, which requires defining the six knot positions. The highest and lowest knot are fixed at the highest and lowest GC bin. GC bins without at least one counted fragment in the real data cannot host a knot. Beyond the outer knots, the GC bias is assumed to remain constant. For the inner four knots, ReSeq tries to find the ideal distribution. First, the four knots are distributed to have an equal spacing according to the number of sites. Then ReSeq maximizes the Poisson likelihood with the flanking biases fixed to the values determined in the first step of the GC and flanking bias fit (Poisson without GC spline). It additionally performs this maximization for all knot distributions that can be reached from the starting distribution by moving only a single of the inner knots, while maintaining knot order. From the performed fits, ReSeq greedily chooses the knot distribution with the highest likelihood as the new starting distribution and repeats this procedure until convergence.

### GC biases renormalization

After the third step of the GC and flanking bias fit, the 101 GC biases are renormalized to remove the normalization parameter and make them comparable across all combinations of reference sequence and fragment length. For this, ReSeq only takes the best-represented GC biases into account to calculate the renormalization factor, which prevents the highly-variable bins from disturbing the normalization. The best-represented GC biases are those with the most sites that cover together at least 80% of the total number of sites. After the normalization, the mean of those best-represented biases equals 1.

### Parameter combination with medians

The flanking biases and the dispersion parameters from the individual fits for each combination of reference sequence and fragment length are combined with a standard median. In contrast, the GC biases need a weighted median, because not all fits contain a high amount of information about every GC bin.

The unnormalized weight of a given GC bias in a fit depends on the amount of sites *l_sites,GC_* with the corresponding GC content and equals to 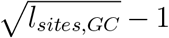. To account for the connection between the biases due to the spline, a fraction of the weight is also added to the other bins depending on the distance:

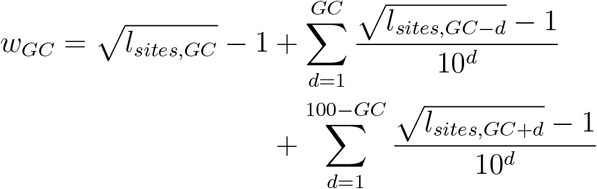

Those weights are then normalized such that the sum over all GC bins equals 1 in each fit, which makes sure that all fits are equally important.

### Corrected reference sequence and fragment length counts

To minimize biases, ReSeq only counts read pairs that pass all filters (mapping quality, forward-reverse order, maximum fragment length). Furthermore, the read pairs must not fall on a site excluded due to low-quality mappings. To avoid a distortion of the counts from the last filter, ReSeq scales the reference-sequence counts from the high-quality part (that remains after filtering) to the full sequence.

To properly handle short fragment lengths, ReSeq searches for adapters, whose presence frequently leads to no or low-quality mappings and subsequent exclusion from the analysis. Hence, if the scan detects adapters, ReSeq will recover otherwise excluded unmapped and low-quality reads, incorporating the pairs into the fragment-length distribution.

### Bias sum estimation

Summing up the bias over all sites is an essential part of ReSeq. It is necessary to calculate the fragment-length biases during the statistics calculation and to normalize the biases during the simulation. To speed up the bias sum, ReSeq calculates in both use cases the sum only every 20th fragment length (the samples) and estimates the other fragment lengths. The sampling stats at the first fragment length with non-zero counts and stops as soon as it encounters a zero count, except if this would exclude sampling positions with at least 10 counts. The estimation of the non-sampled fragment length is different in the statistics calculation and the simulation due to different requirements.

In the statistics calculation, ReSeq first obtains the fragment-length biases for the sampled lengths (as described in the Results section). After this fit, ReSeq extends the estimates to all biases by using the ratio of counts over bias from the samples. This ratio is almost constant except for the shorter fragment length. Therefore, it is much easier to describe by a natural spline and requires less sampling than the biases itself. ReSeq estimates the parameters of the natural spline by a non-linear least square fit to the samples using the L-BFGS algorithm [36, 37] for minimization. ReSeq starts by assigning a knot to every sample and then gradually removes the knots for which the sum over the squared residuals 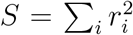 increases the least, until only 6 knots are left. Occasionally, ReSeq performs a greedy search for the optimal knot positions during the reduction, which is similar to the one described above in *GC knot adjustment*. Finally, ReSeq uses the optimized spline to calculate the bias estimates for all fragment lengths.

In the simulation, ReSeq only requires the total bias sum over the whole genome and all fragment lengths. Thus, the precision of individual values is secondary and another approach is chosen, where ReSeq applies a spline to interpolate the ratio of the bias sum over the fragment-length bias using every sample as a knot. From the ratios, the bias sums are retrieved and subsequently summed over all fragment lengths. Additionally, the simulation requires the maximum bias for each fragment length, which ReSeq estimates by using the maximum ratio over all samples multiplied with the respective fragment-length bias.

### Reproducibility of simulator comparison

Except for the parts that are specifically mentioned in this paragraph, everything for this paper was run through snakemake [50] with the scripts available on github [45]. Figures 3, S1, S3, S4, S20 and S21 represent intermediate values from ReSeq. The output of the intermediate values requires additional calculations and disk writing in performance-critical functions. Therefore, it is triggered by hard-coded flags and not by parameters to allow the compiler to remove unneccessary code and prevent speed-loss during normal use, when the additional outputs are not needed. Due to the hard-coded flags, the creation of the intermediate values is not included in the snakemake pipeline. Instead, we added the csv files containing the intermediate values to the git repository. Moreover, the computing requirements were manually extracted from the timing files and inserted into a csv file included in the git repository. The IGV [34, 35] screenshots included in Figure 2 are also in the repository, so that all plotting scripts can be called. We compared the newest versions of the simulators (Table S1) and all other software and their versions are stated in Table S2. The datasets with sample IDs Ecoli1_L001, Bcereus1_L001, Rsphaeroides1_L001, Ecoli1_L002, Ecoli1_L003, Ecoli2_L001 and Ecoli3_L001 were downloaded as raw data from Illumina BaseSpace. We ran the conversion from bcl to fastq format locally with disabled adapter trimming.

### Simulations

The references for each dataset were corrected with pilon [51] before subsequent trainings, simulations and evaluations. If not stated otherwise, all reads were mapped with bowtie2 [52]. For diploid organisms, freebayes [53] called the variants and we filtered out reported variants with qualities below 10. For simulators requiring coverage as a parameter, we calculated the median from the coverage summary of samtools stats [54]. Since BEAR only accepts the number of reads as a parameter, we set it to match the total number of reads in the real dataset. Additionally, BEAR requires the sequence abundances, where we used its own parametric abundance profile generation with high species complexity. The insertion and deletion rates provided to ART were obtained from parsing the CIGAR strings in the bam files. With *M*, *I*, *D* being the number of matches, insertions and deletions in the CIGAR strings, respectively, the insertion rate is 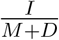 and the deletion rate is 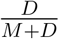. The fragment lengths obtained from the mapped reads were fitted with a Gaussian to retrieve the fragment length parameters. In the case of Bc-Hi4000-Nextera, Ec-Hi4000-Nextera and Rs-Hi4000-Nextera, the fragment length was not Gaussian distributed, but followed an exponential decay. Therefore, we set the mean to the lowest value accepted by all simulators, which is the read length plus one, and manually fitted the variance for the second half of the Gaussian. The length of sliding windows were set for all datasets to the specified fragment length.

### Evaluation of simulator performance

To evaluate performance, we counted 51-mers with Jellyfish [55], except for the human and mouse datasets, where memory consumption skyrocketed and kmc [56] using disk storage was applied instead. The preqc module of sga [57] calculated in a reference-free way the mean quality values and error rates by position as well as the simulated contig length N50. We modified the preqc plotting script to adjust it to the general style. Further plots were obtained with ReSeq using bowtie2 [52] mappings. In case of simulated datasets without adapters, the evaluation with ReSeq requires to specify TruSeq adapters as a decoy. To calculate the coverage correlation, we counted the base coverage with samtools depth [54]. For the short-read assembly, we called sga [58] with default parameters, except for the ropebwt algorithm for indexing. The assembly statistics were calculated by QUAST [39] with the minimum contig length set to zero.

### Speed and memory benchmarks

We benchmarked the speed and memory usage with GNU time. The single-threaded processes were run on a machine with four AMD Opteron(tm) Processor 6376 with a total of 64 cores and 512GB of memory operating Ubuntu 16.04.9. Due to their single threaded nature, other programs were running in parallel to get a realistic use case. The multi-threaded processes ran on single cluster nodes with 384 GB memory and two Intel Xeon Gold 6126 resulting in 48 vCPUs. All 48 possible threads were provided to the processes, even though ReSeq is the only tool to use all of them. To report CPU time, we summed user and system time. We summed CPU times of all processes in a category, as well as elapsed times for multi-threaded processes. For single-threaded processes, we summed the elapsed times only if the processes were dependent and needed to be run in series. Otherwise we took the maximum. For maximum memory, we report the maximum resident set size. The memory maxima were summed if single-threaded processes are run in parallel and the maximum was taken if they are run in series. For multi-threaded processes, we always took the maximum memory.

## Supporting information

Supplementary figures, tables and formulas

QUAST assembly report

ReSeq Manual

## 1 Availability of data and materials

Only public data were used during this study. The datasets with accession IDs SRR490124, SRR3191692, ERR2017816, ERR3085830 and ERR1955542 can be downloaded from the European Nucleotide Archive: https://www.ebi.ac.uk/ena

The datasets with sample IDs Ecoli1_L001, Bcereusl_L001, Rsphaeroidesl_L001, Ecoli1_L002, Ecoli1_L003, Ecoli2_L001 and Ecoli3_L001 are part of the ‘HiSeq 4000: Nextera XT (B.cereus, E.coli, R.sphaeroides)’ project in Illumina’s BaseSpace Sequence Hub: https://basespace.illumina.com

ReSeq and all of its code are available under the MIT License at: https://github.com/schmeing/ReSeq

The snakemake pipeline and custom scripts used for this publication are available under: https://github.com/schmeing/ReSeq-paper

## Acknowledgements

The authors thank members of the Robinson Lab at the University of Zurich for valuable feedback and the Functional Genomics Center Zurich for providing helpful datasets.

## Author’s contributions

Stephan Schmeing (SMG) conceived the study. MDR supervised the study. SMG developed the simulator and its methods and performed the benchmarking. SMG and MDR wrote the manuscript. All authors read and approved the final manuscript.

