## Supplementary figures, tables and formulas for "ReSeq simulates realistic Illumina high-throughput sequencing data"

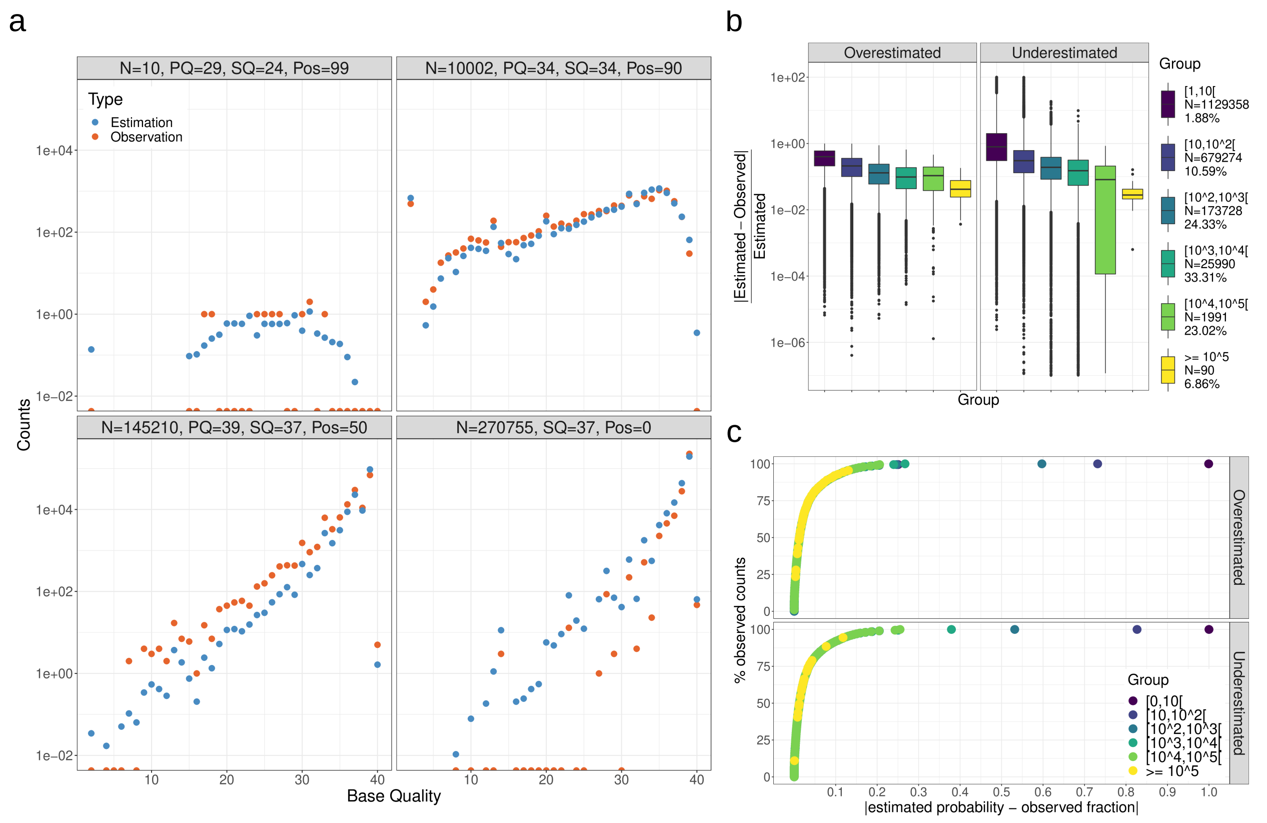


**Figure S1 Accuracy of two-dimensional estimation of quality matrix for Ec-Hi2000-TruSeq.** All values are only based on the matrix part for the first read, nucleotide A and error rate 0. a) Sample slices of the matrix comparing the estimation to the observed counts for different values of previous quality, sequence quality and position. $N$ specifies the total number of observed counts in this slice. The two lower slices with high total number of counts show consistent, moderate over- or underestimation, while the top left slice with only 10 total counts shows how the estimation deals well with low numbers of observed counts. b) Relative difference between observation and estimation grouped by number of observed counts. The plot lists for each group the number of compared values $N$ and the percentage of counts from the matrix. Deviations from low counts are also caused by the integer nature of counts and not necessarily an estimation error. More than 75% of the values with at least 100 observed counts deviate by less than 30% from the observation. c) The maximum absolute deviation for the given percent of all observed counts. The deviation is normalized by the total number of counts in each slice of previous quality, sequence quality and position. The deviations falling into each percentage bin are grouped based on their observed counts. For more than 80% of the data the deviation is below 0.05.

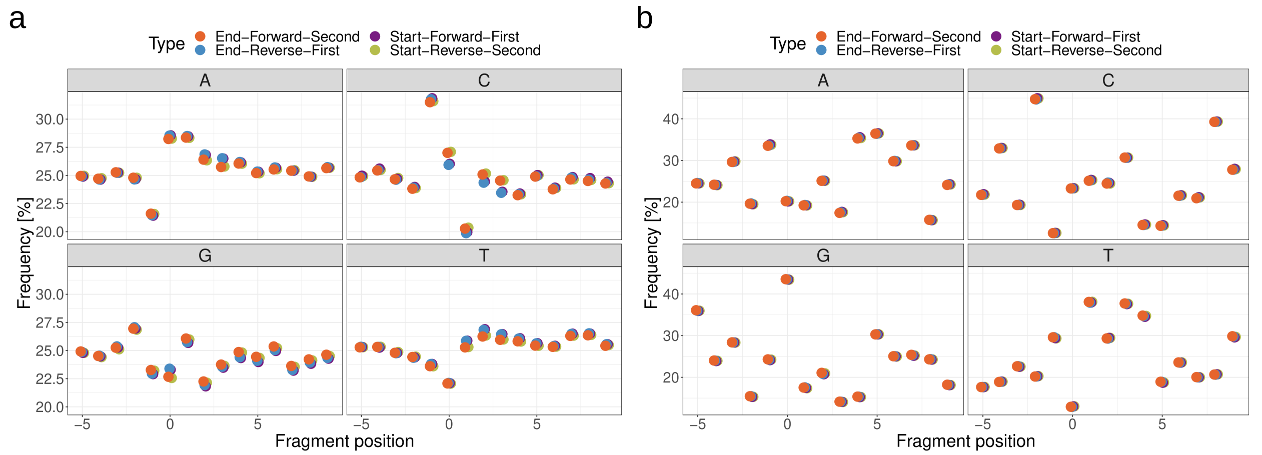


**Figure S2 Observed frequency of nucleotides in the fragments.** The position is relative to the fragment start/end. Positive positions and zero are in the fragment. Negative positions are outside of the fragment. The flanking bias is less pronounced in the mechanically-fragmented Ec-Hi2000-TruSeq (a) compared to Ec-Hi4000-Nextera (b), where enzymes have been used. The colour specifies, whether the flanking sequences of the start or end of the fragment are shown, if the fragment is on the forward or reverse strand and if the first or second read maps on this side of the fragment. A small deviation between first and second reads is visible (a), which is not included in Reseq's coverage model.

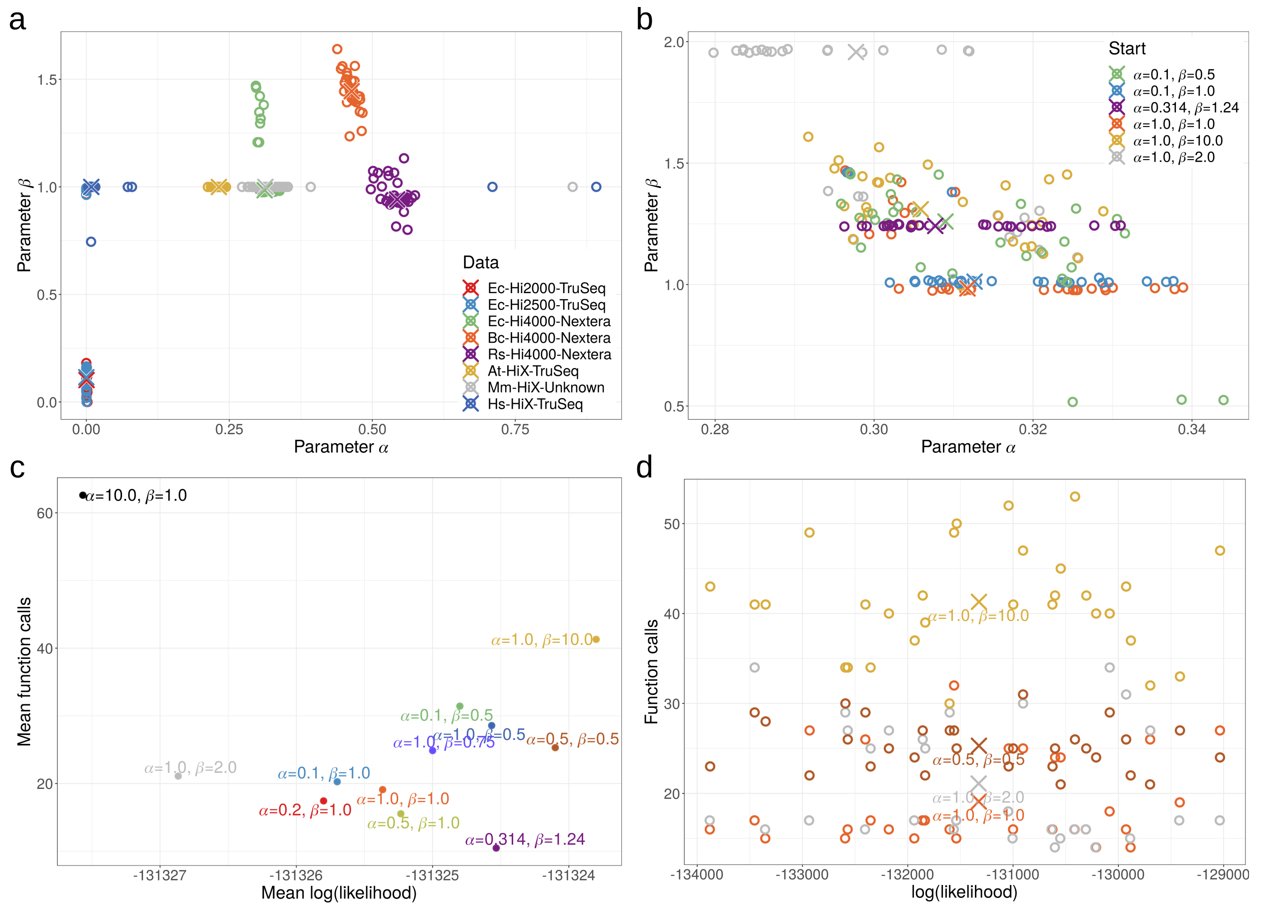


**Figure S3 Dispersion parameter overview.** a) Comparison of the resulting parameters over 8 tested datasets. Each circle is a fit. Crosses represent the median (final result) for each dataset. Ec-Hi2000-TruSeq and Ec-Hi2500-TruSeq seem to suppress parameter $\alpha$, while all other datasets require it. A likely explanation are patterned flow cells, which are build into newer Illumina sequencers, but not into the HiSeq2000/2500 used for those two datasets. Strong increases of optical duplicates for patterned flow cells have been reported before [1]. That parameter $\beta$ is fixed to its start value of 1 is a precision artifact (panel b) mostly observed for low-coverage datasets. b) Converged parameters for Ec-Hi4000-Nextera depending on the start parameters. Circles are single fits and crosses medians for each set of start parameters. If parameter $\beta$ starts too close to the optimal value, it remains in proximity of its starting value. Changing the default start parameters is not a good solution (panel c). c) Final log(likelihood) vs. total number of calls to the likelihood calculation for Ec-Hi4000-Nextera. Each dot represents the mean over all fits. Parameter $\alpha$ has a stronger influence on the likelihood than parameter $\beta$. Start values further away from the optimal value are not guaranteed to increase the likelihood and often need more function calls. d) Final log(likelihood) vs. total number of calls to the likelihood calculation for Ec-Hi4000-Nextera. Each circle is a single fit and the crosses are the mean values shown in (c). The mean shift of log(likelihood) caused by different start parameters is minor compared to the spread between individual fits.

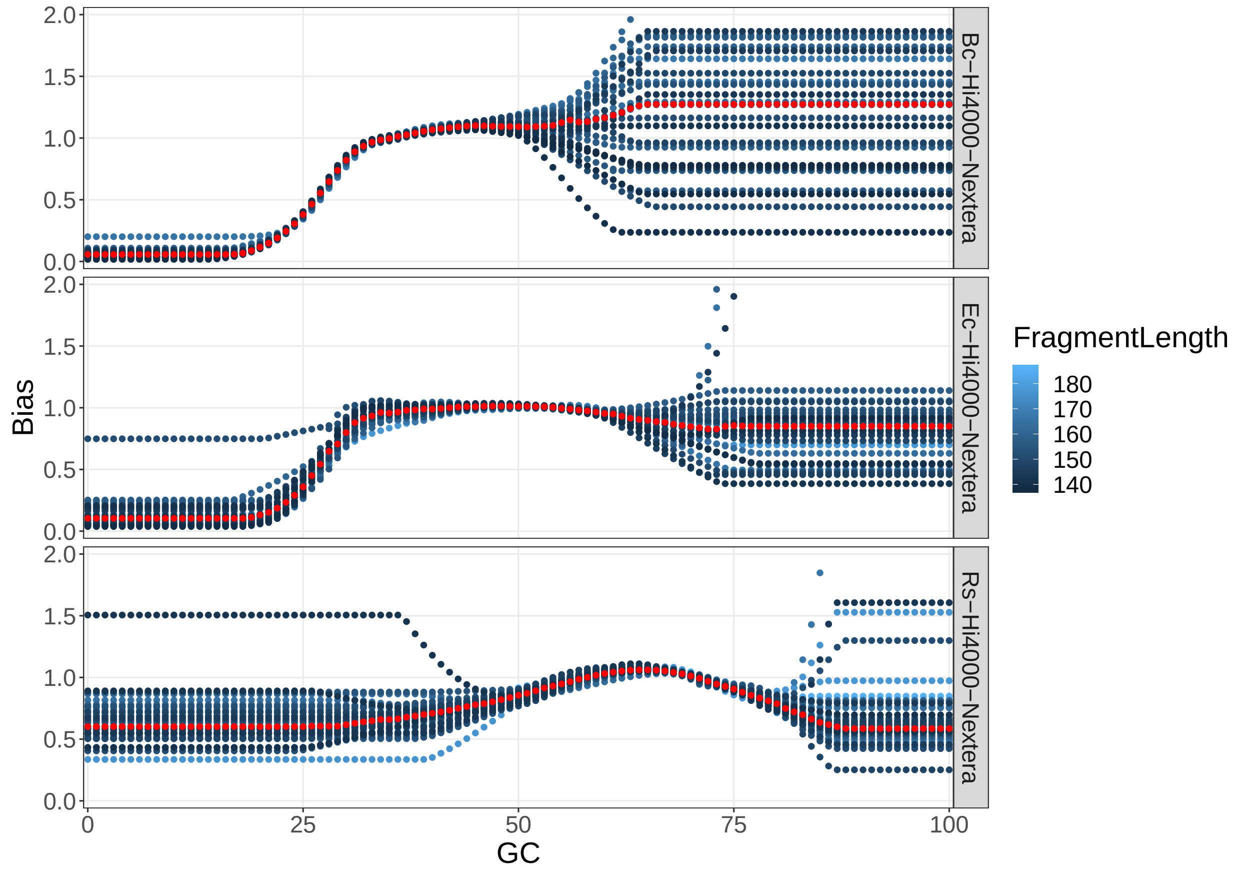


**Figure S4 Comparison of GC bias for three dataset created with Nextera adapters.**

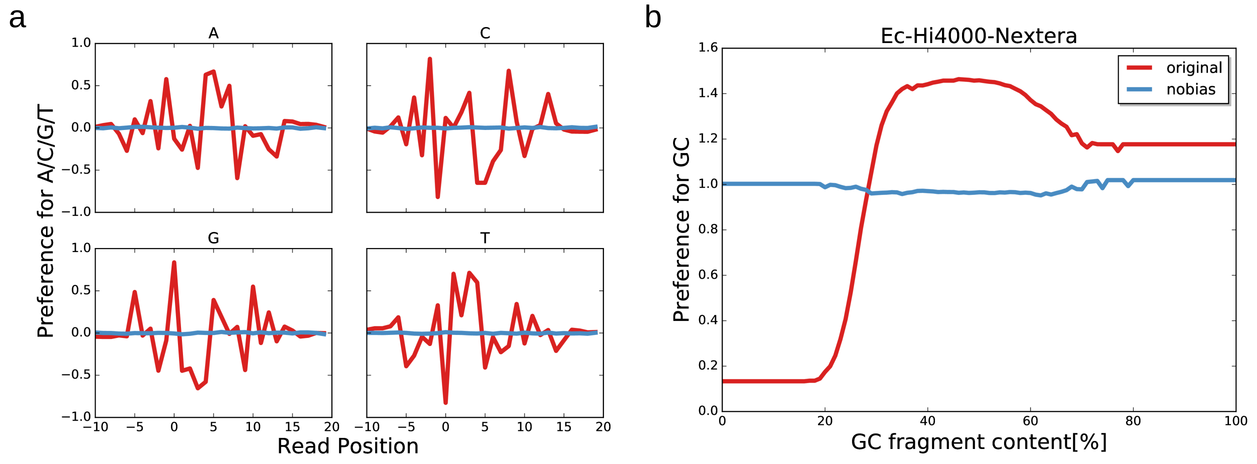


**Figure S5 Estimated bias for simulated, uniformly distributed data based on Ec-Hi4000-Nextera.** The bias estimated for the original dataset is given as a reference for the magnitude of the biases. a) Flanking bias b) GC bias

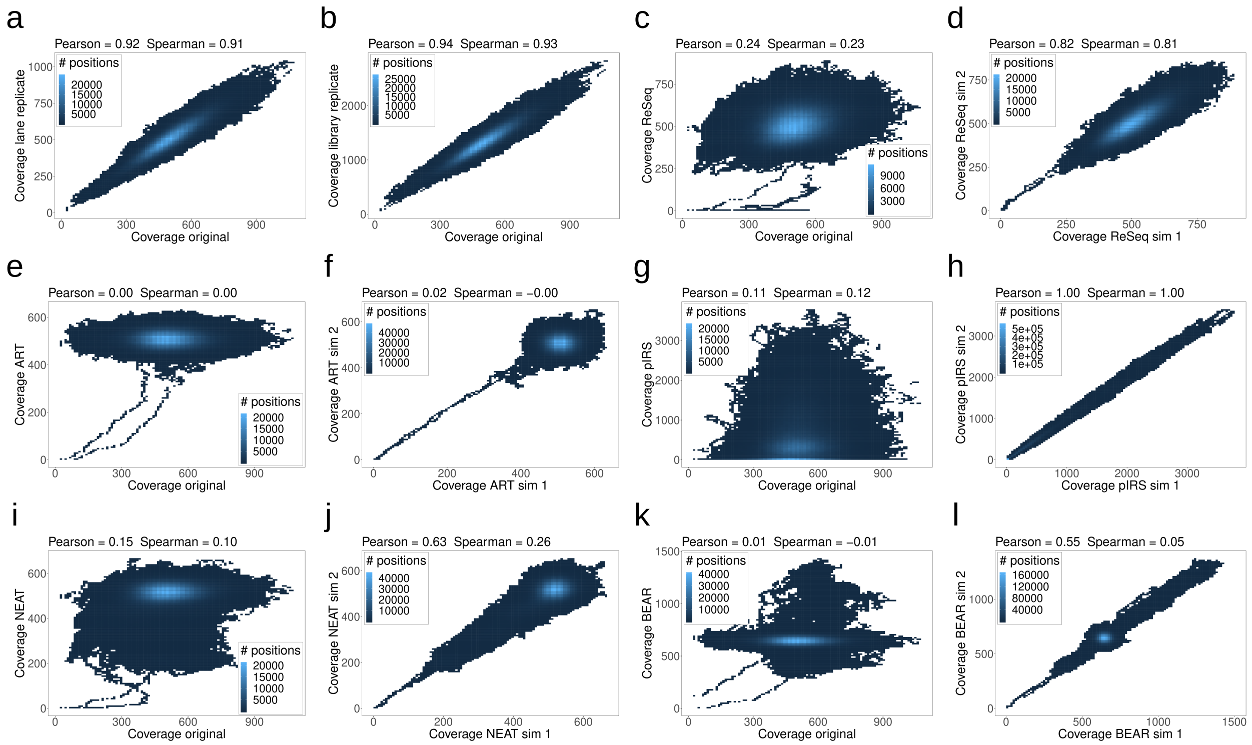


**Figure S6 Correlations of real and simulated coverages across datasets.** a) Ec-Hi4000-Nextera (Ecoli1_L001) vs Ecoli1_L002 b) Ec-Hi4000-Nextera (Ecoli1_L001) vs. Ecoli2_L001 c) Ec-Hi4000-Nextera vs. ReSeq d) ReSeq e) Ec-Hi4000-Nextera vs. ART f) ART g) Ec-Hi4000-Nextera vs. pIRS h) pIRS i) Ec-Hi4000-Nextera vs. NEAT j) NEAT k) Ec-Hi4000-Nextera vs. BEAR l) BEAR

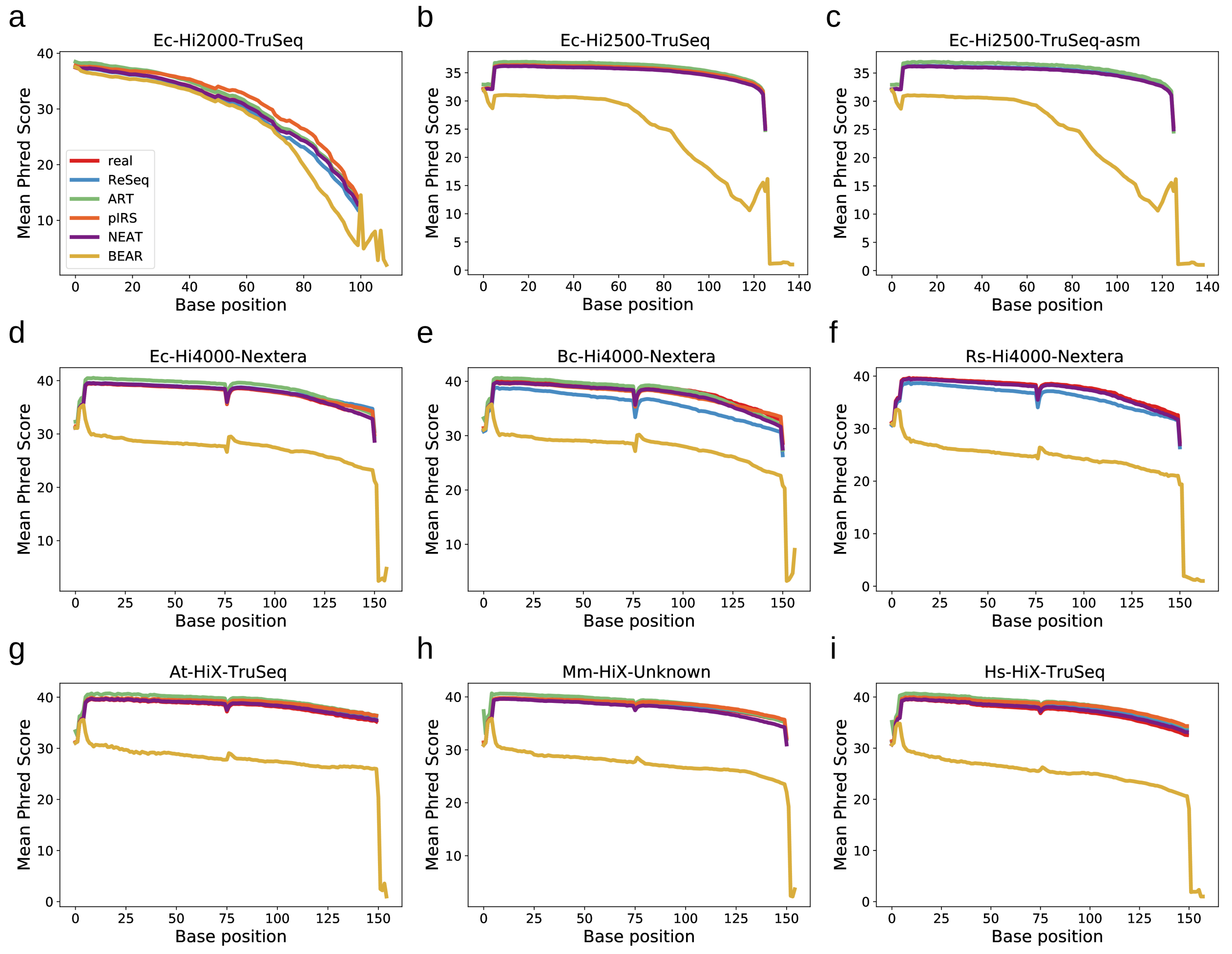


**Figure S7 Mean quality values by position for real and simulated data.** Note that: c) preqc crashed for pIRS f) preqc crashed for pIRS and ART

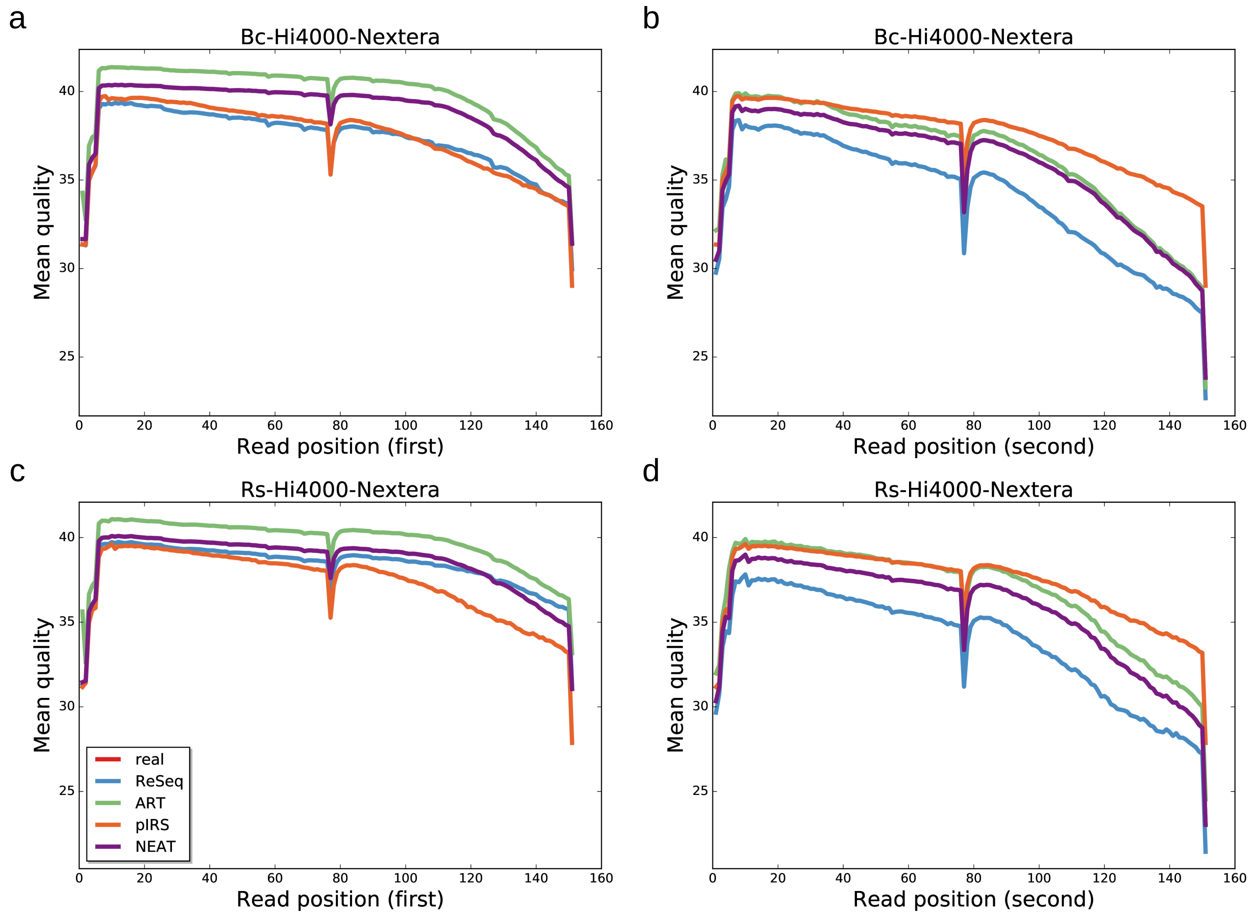


**Figure S8 Mean quality values by position for real and simulated data, separately for first and second read.** NEAT follows the real data perfectly and covers the real data line. This is slightly different from Figure S7, where preqc filters some reads. BEAR is excluded from this plot to achieve better resolution for the other simulators.

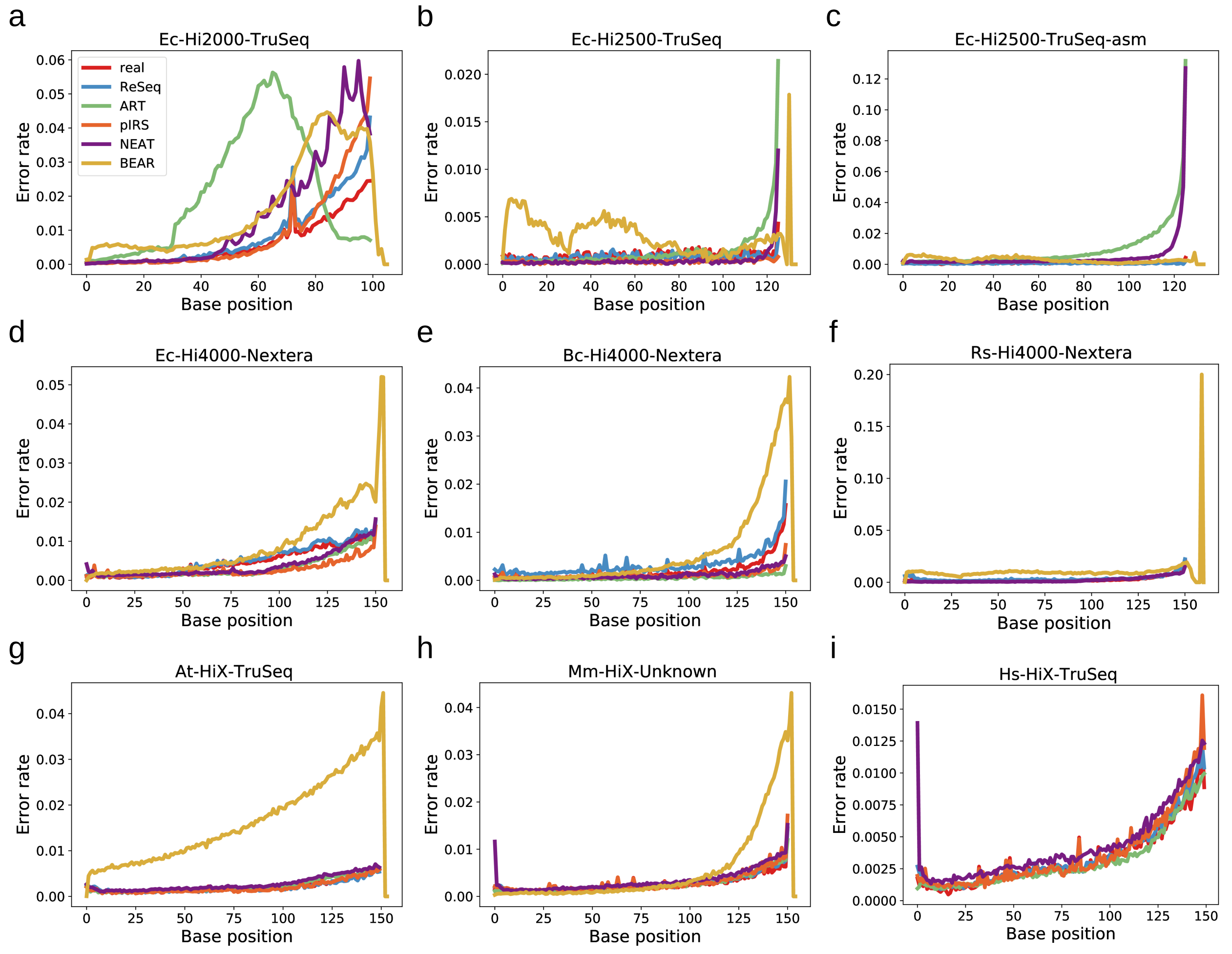


**Figure S9 Mean error rates by position for real and simulated data.** Note that: c) preqc crashed for pIRS f) preqc crashed for pIRS and ART i) BEAR was excluded due to sporadic reports of error rates up to 1 for positions up to 700.

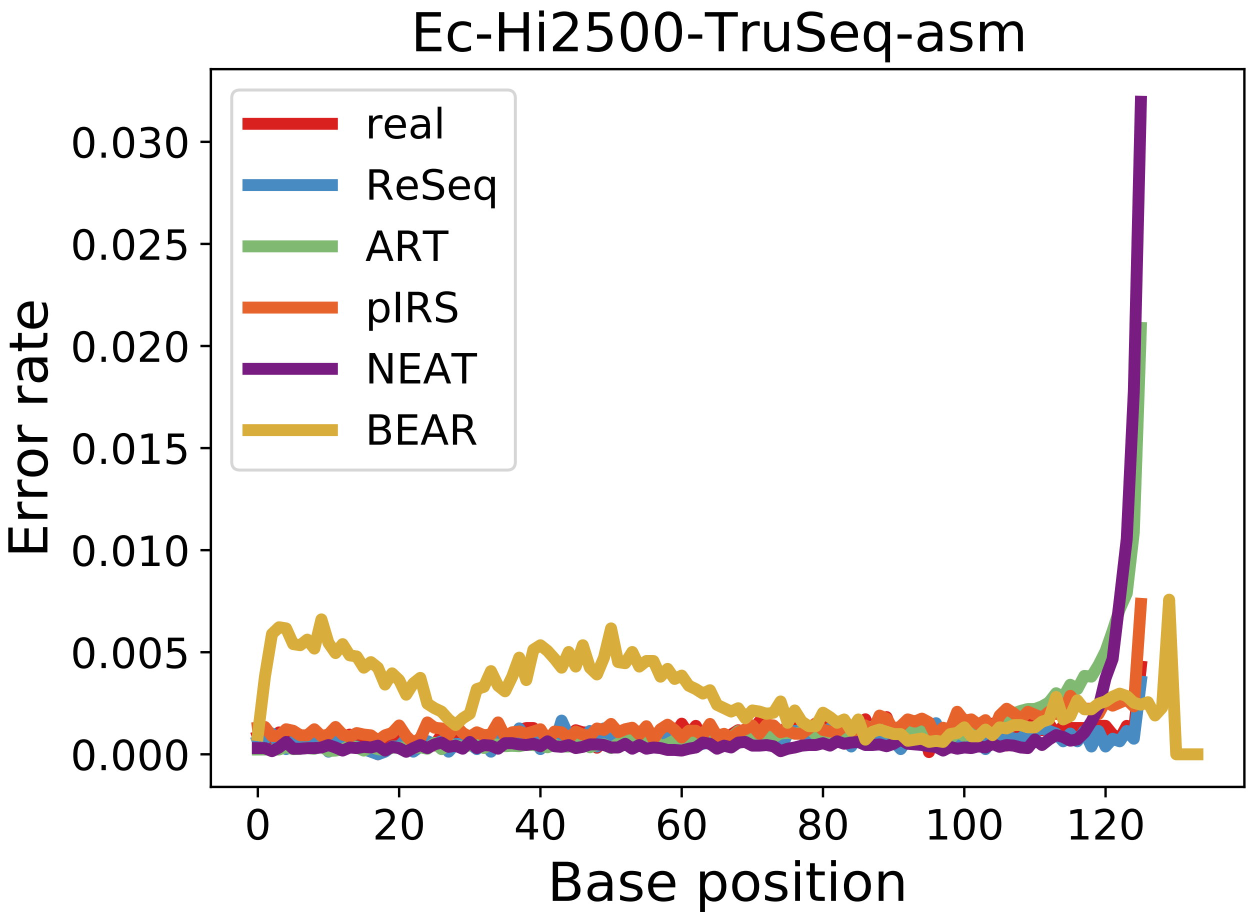


**Figure S10** Mean error rates by position for real and simulated data for Ec-Hi2500-TruSeq-asm with the median coverage from Ec-Hi2500-TruSeq provided to all simulators except BEAR.

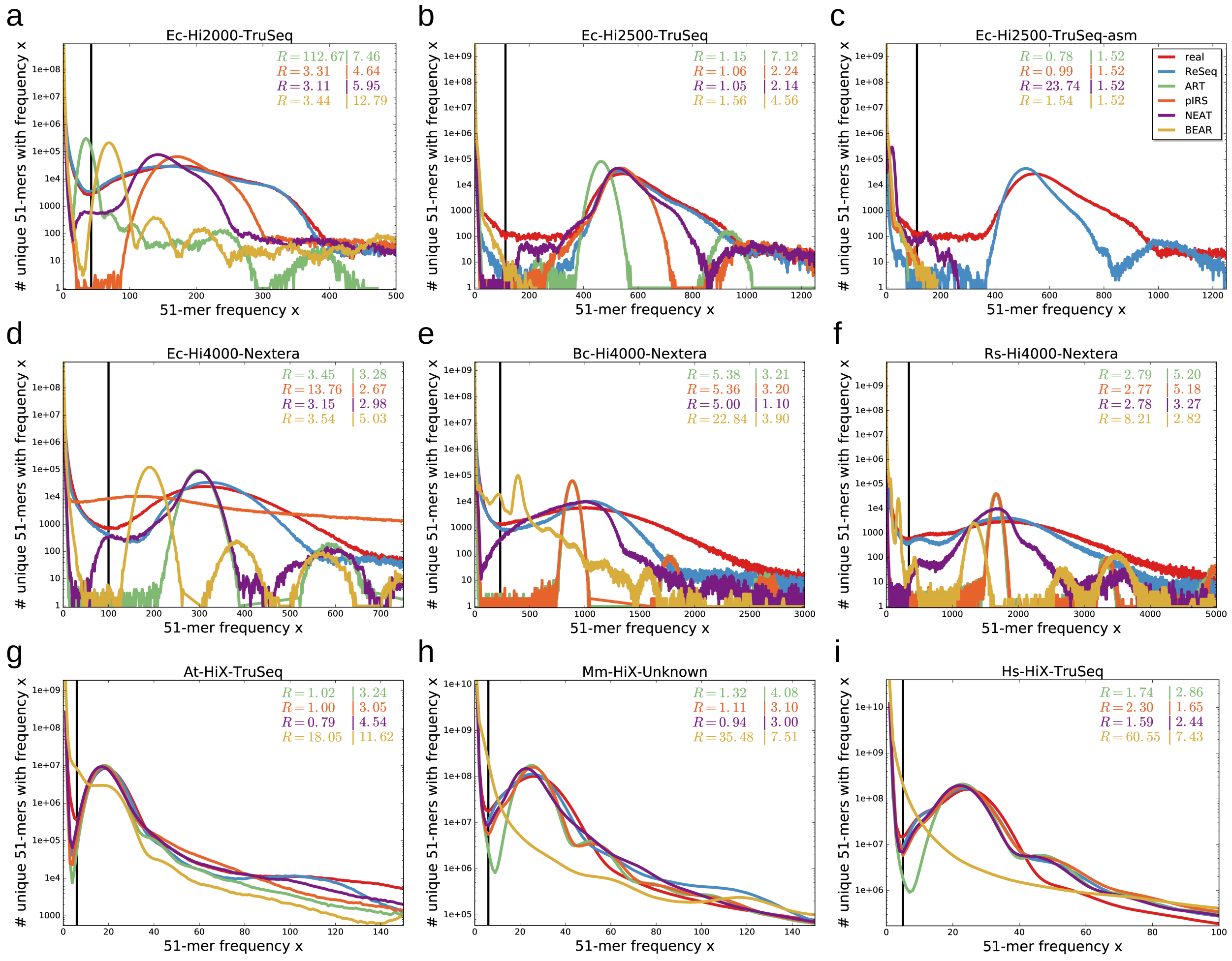


**Figure S11 51-mer spectra of real and simulated data.** The shape and position of peaks reflects the coverage distribution, while the exponential decrease at low frequencies is defined by systematic errors. The black lines show the minimum between the exponential decrease and signal peak for real data. The first value of $R$ gives the sum of relative deviations for frequencies (x) up to the minimum. The second value of $R$ gives the sum of absolute deviations for frequencies larger than the minimum. Both values are stated relative to ReSeq. When the signal peak of the simulation does not exist or starts before the minimum in real data, the values lose their interpretability, which happens especially for ART (a,c), pIRS (c,d), NEAT (c) and BEAR (b,c,f-i).

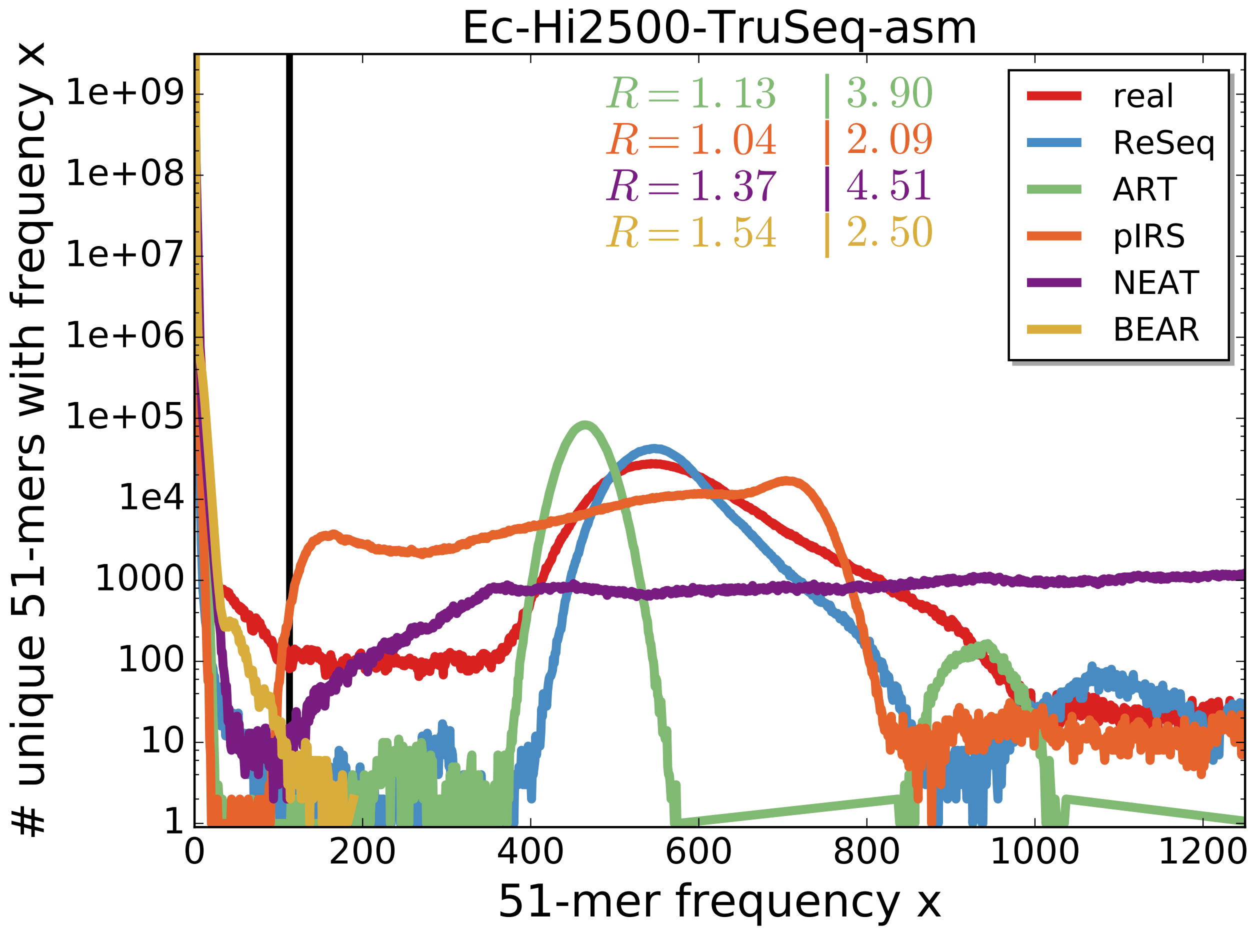


**Figure S12 51-mer spectra of real and simulated data for Ec-Hi2500-TruSeq-asm with the median coverage from Ec-Hi2500-TruSeq provided to all simulators except BEAR.** The shape and position of peaks reflects the coverage distribution, while the exponential decrease at low frequencies is defined by systematic errors. The black lines show the minimum between the exponential decrease and signal peak for real data. The first value of $R$ gives the sum of relative deviations for frequencies (x) up to the minimum. The second value of $R$ gives the sum of absolute deviations for frequencies larger than the minimum. Both values are stated relative to ReSeq. When the signal peak of the simulation does not exist or starts before the minimum in real data, the values lose their interpretability, which happens for BEAR and to a smaller extent for pIRS.

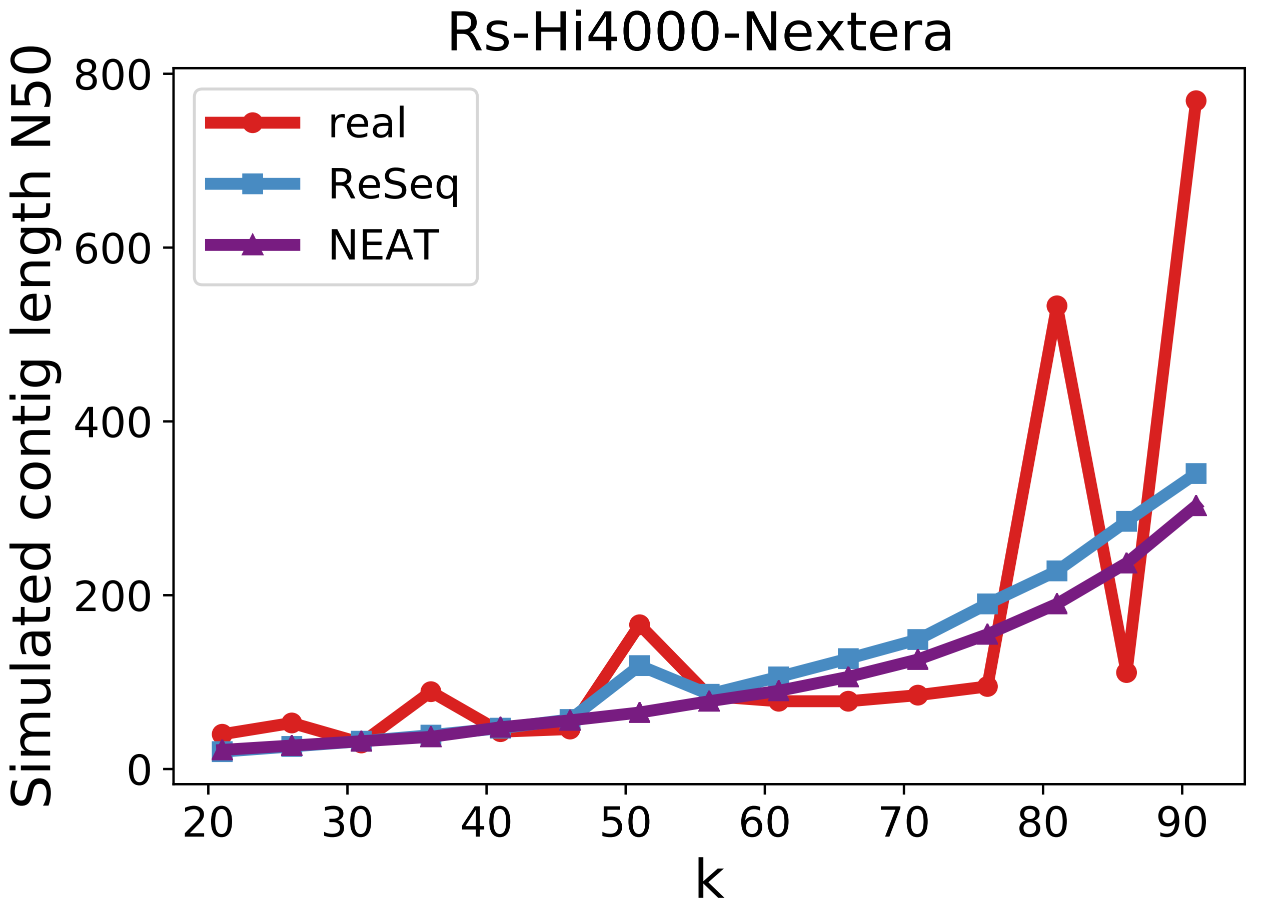


**Figure S13 Simulated contig N50 for real and simulated data for RS-Hi4000-Nextera with BEAR removed.** Bear was removed due to its strong deviation from the real N50 values. pIRS and ART are also missing, because the preqc module crashed while processing them.

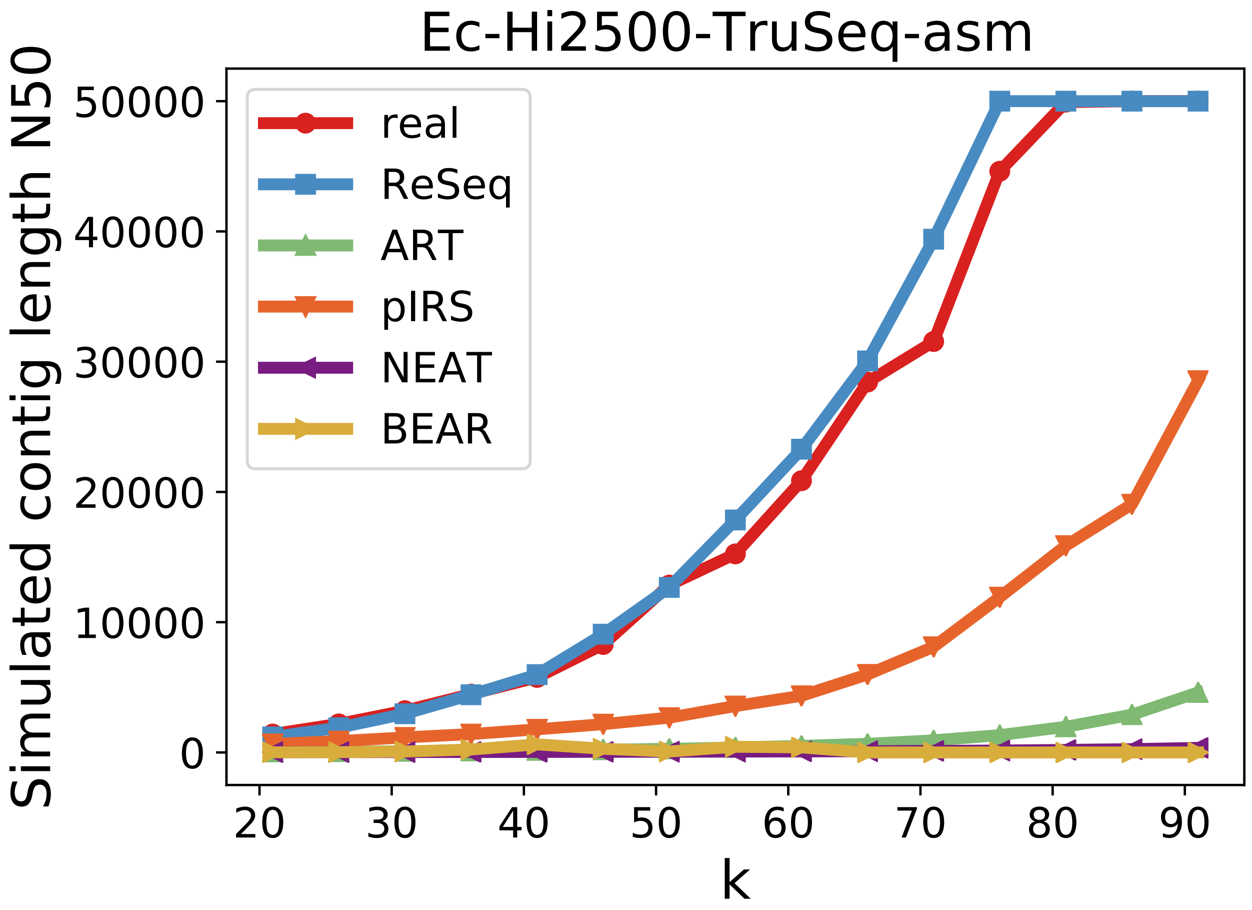


**Figure S14** Simulated contig N50 for real and simulated data for Ec-Hi2500-TruSeq-asm with the median coverage from Ec-Hi2500-TruSeq provided to all simulators except BEAR.

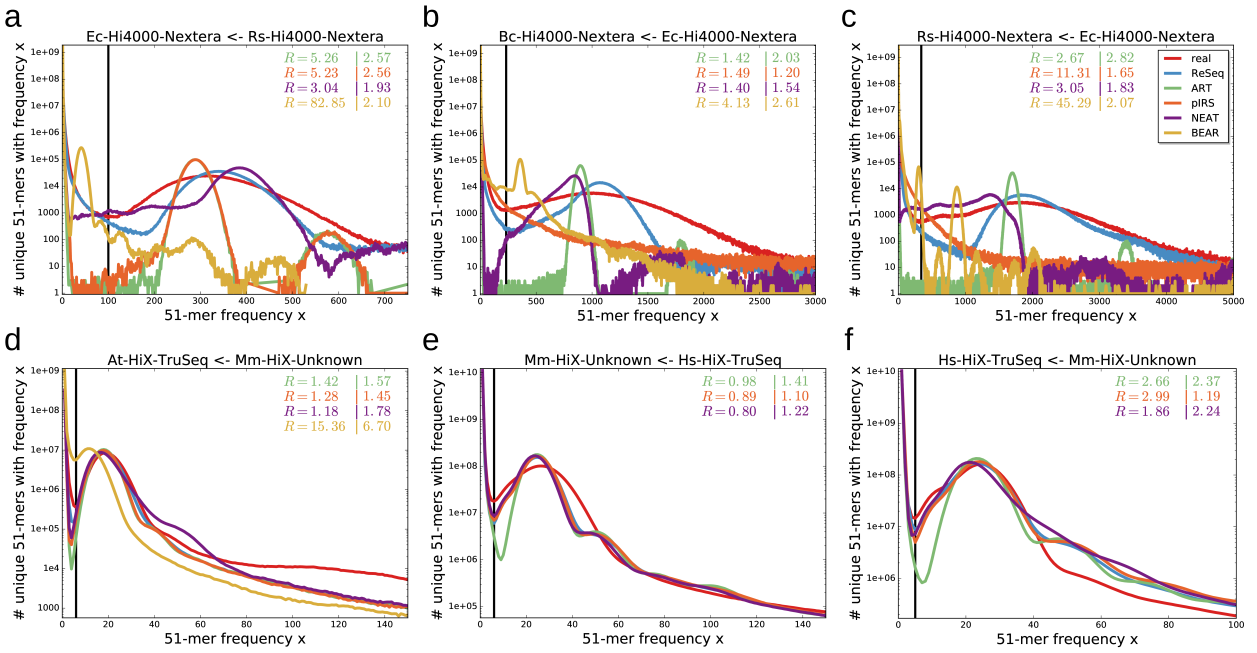


**Figure S15 51-mer spectra of real and cross-simulated data.** The simulation profiles are trained on a different training dataset. The pairings are given in the plot titles in the following notation: "simulated dataset $\leftarrow$ training dataset". The shape and position of peaks reflects the coverage distribution, while the exponential decrease at low frequencies is defined by systematic errors. The black lines show the minimum between the exponential decrease and signal peak for real data. The first value of $R$ gives the sum of relative deviations for frequencies (x) up to the minimum. The second value of $R$ gives the sum of absolute deviations for frequencies larger than the minimum. Both values are stated relative to ReSeq. When the signal peak of the simulation does not exist or starts before the minimum in real data, the values lose their interpretability, which happens for BEAR in panel a and c.

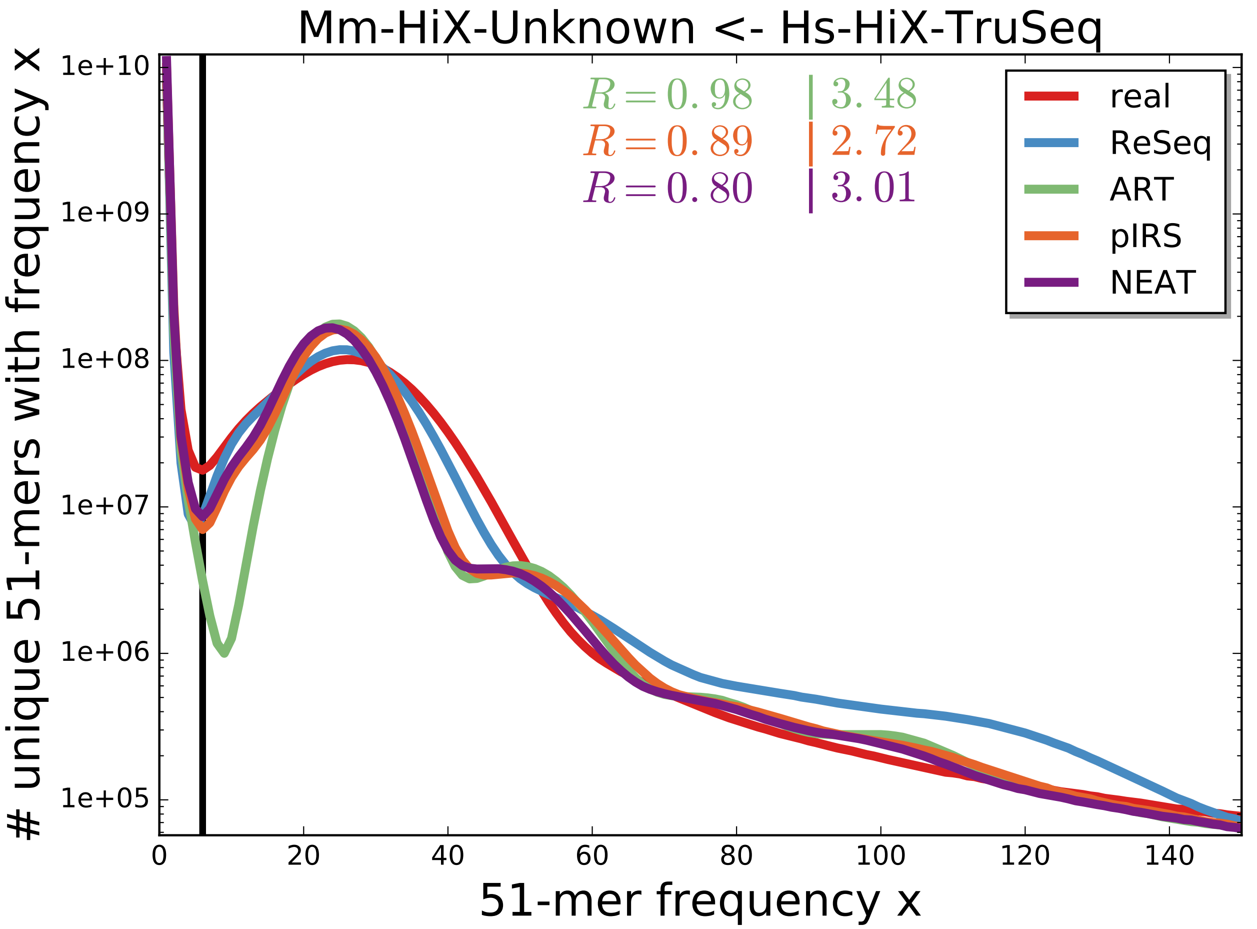


**Figure S16 51-mer spectrum for real and simulated data of Mm-HiX-Unknown.** The simulations are trained on Hs-HiX-TruSeq. ReSeq was provided with the reference bias trained on Mm-HiX-Unknown to reproduce the known fluctuations in chromosome abundance in this cell-line. The shape and position of peaks reflects the coverage distribution, while the exponential decrease at low frequencies is defined by systematic errors. The black lines show the minimum between the exponential decrease and signal peak for real data. The first value of $R$ gives the sum of relative deviations for frequencies (x) up to the minimum. The second value of $R$ gives the sum of absolute deviations for frequencies larger than the minimum. Both values are stated relative to ReSeq. When the signal peak of the simulation does not exist or starts before the minimum in real data the values lose their interpretability.

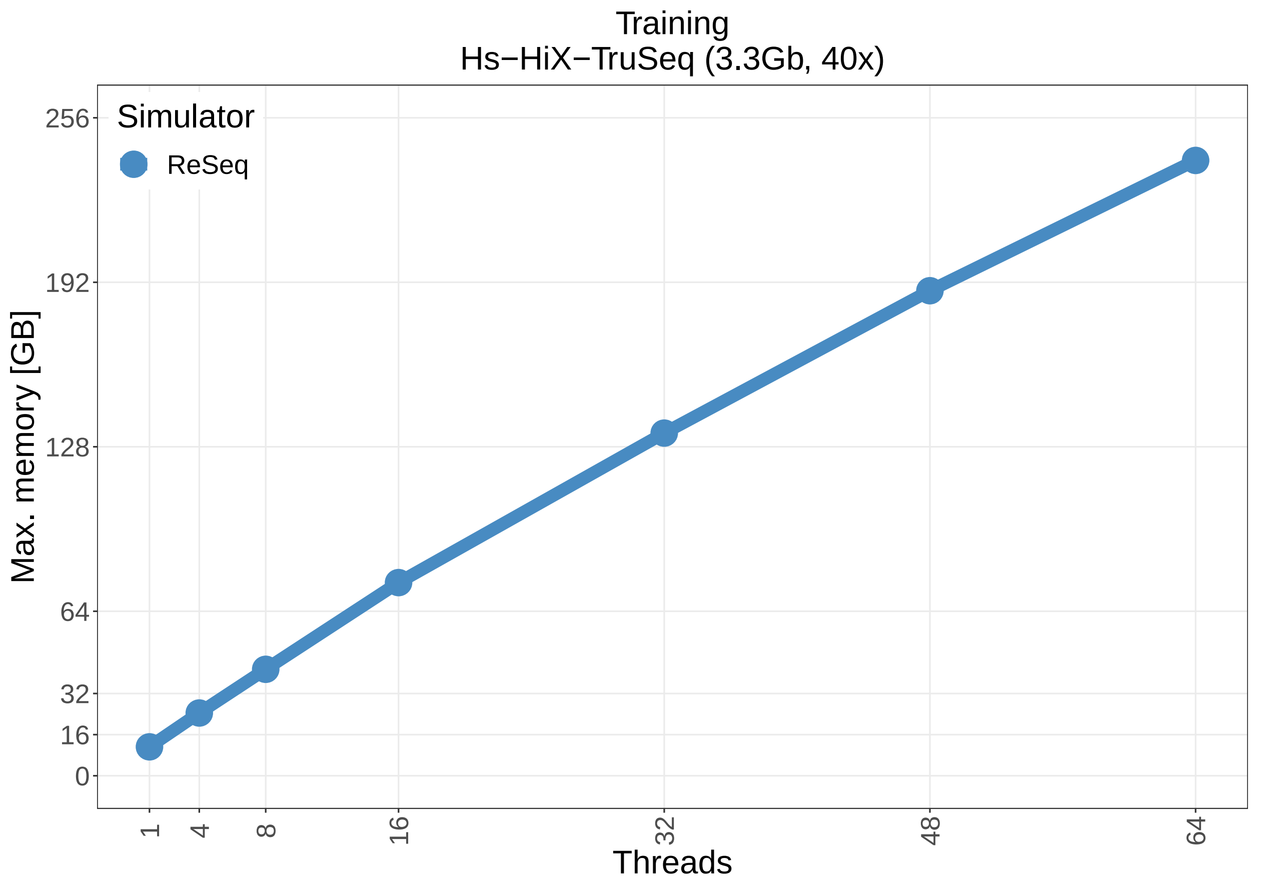


**Figure S17 Memory requirements for ReSeq training on Hs-HiX-TruSeq for different numbers of computing threads.**

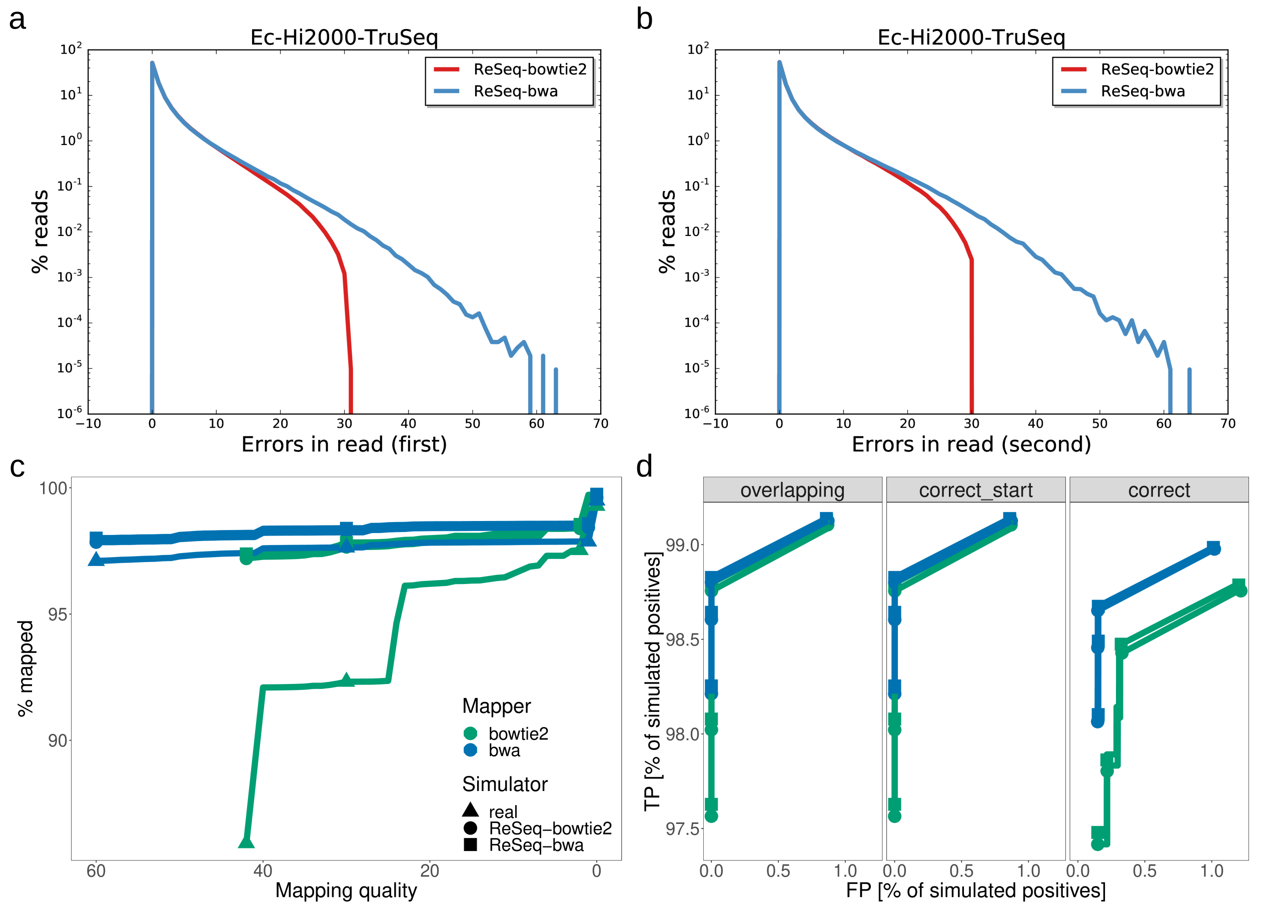


**Figure S18 Difference in errors per read between ReSeq trained on bowtie2 and bwa mappings.** a,b) Number of errors observed by ReSeq in the Ec-Hi2000-TruSeq dataset using bowtie2 and bwa mappings of first (a) and second reads (b). c) Mapping rate over mapping quality for real and simulated data, with all reads removed that have more than 10 simulated errors. Real data are unfiltered. Mapping qualities are cumulative, i.e. all mapping qualities at the given score or higher. Markers are only shown for mapping qualities 0,2,30,42 for bowtie and 0,1,30,60 for bwa. Due to the highly-erroneous read removal, the mapping rates for bowtie2 are nearly identical independent of what mapper was used for the training data. d) Mapping accuracy for all mapping quality thresholds with all reads removed that have more than 10 simulated errors. Markers are only shown for mapping qualities 0,2,30,42 for bowtie and 0,1,30,60 for bwa. Positives are mapped reads, which fulfill the correctness criteria (TP) or do not (FP). Overlapping: True and mapped positions overlap independent of strand. Correct_start: Perfect match of start position and strand. Correct: Perfect match of start and end positions and strand. Due to the highly-erroneous read removal, bwa scores identically, when requiring perfect matches, independent of what mapper was used for the training data.

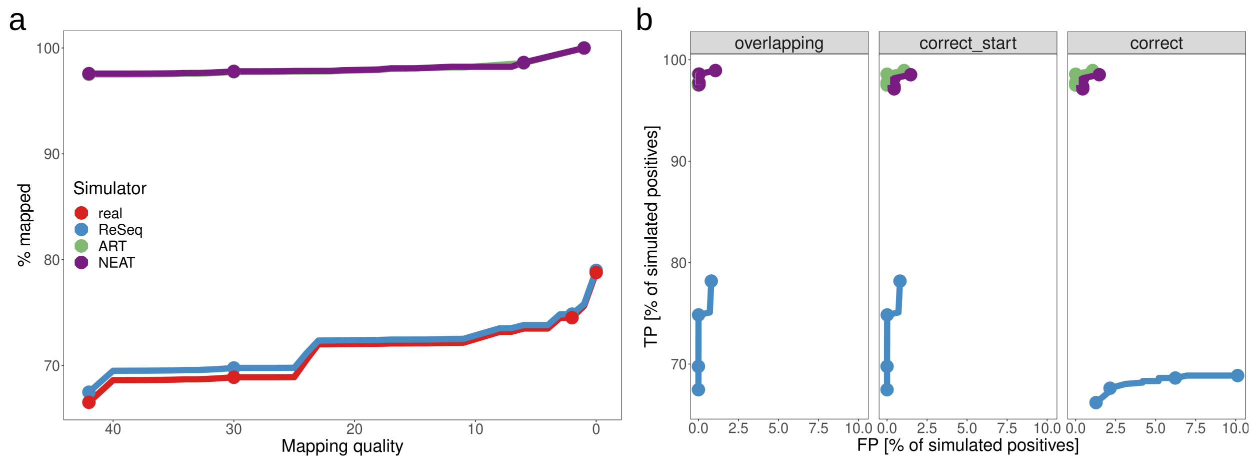


**Figure S19 Comparison of bowtie2 mapping on Ec-Hi4000-Nextera for different simulations.** pIRS and BEAR have been omitted for this analysis due to their inaccurate representation of the real coverage distribution (S6g) or error rate (S9d) for this dataset. The discrepancies caused by not simulating adapters (ART, NEAT) are clearly visible. a) Mapping quality distribution. Mapping qualities are cumulative, i.e. all mapping qualities at the given score or higher. Markers are only shown for mapping qualities 0,2,30,42. Instead of 0 and 2, ART and NEAT show markers for mapping quality 1 and 6, which are the two lowest assigned qualities in their case. b) Mapping accuracy for all mapping quality thresholds. Markers are only shown for mapping qualities 0,2,30,42. Instead of 0 and 2, ART and NEAT show markers for mapping quality 1 and 6, which are the two lowest assigned qualities in their case. Positives are mapped reads, which fulfill the correctness criteria (TP) or do not (FP). Overlapping: True and mapped positions overlap independent of strand. Correct_start: Perfect match of start position and strand. Correct: Perfect match of start and end positions and strand.

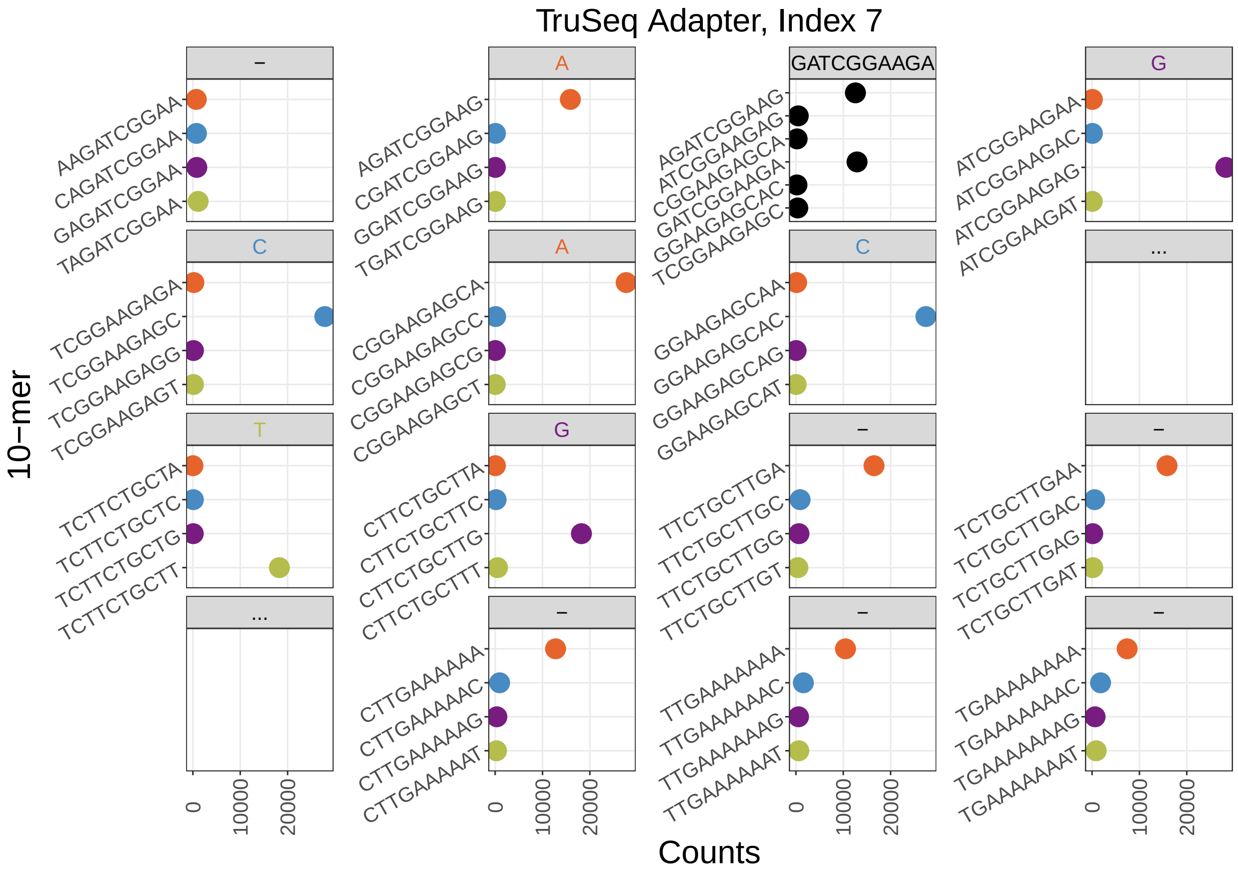


**Figure S20 10-mers and their counts for the automatic adapter detection in first reads of Ec-Hi2000-TruSeq.** The panel with black dots and a full 10-mer as label is the starting 10-mer. Only the six 10-mers with the highest counts are shown for this panel. The other panels show the four possible extending 10-mers for both sides with the true adapter sequence in the label. The direction is from left to right and then from top to bottom. Empty panels with dots in the label represent positions skipped in the plotting. The first and the last panel are not included in the detected adapter sequence, because they fulfill the stopping requirement of the second highest count being bigger than one fifth of the highest count. The poly-A tail, visible in the six panels after the end of the true adapter sequence, is cut from the detected adapter sequence. The only panel that has more than one dominant 10-mer is the starting panel, but the two dominant 10-mers in it are just shifts by one position. Therefore, independent of the selected 10-mer the final detected adapter sequence is the same.

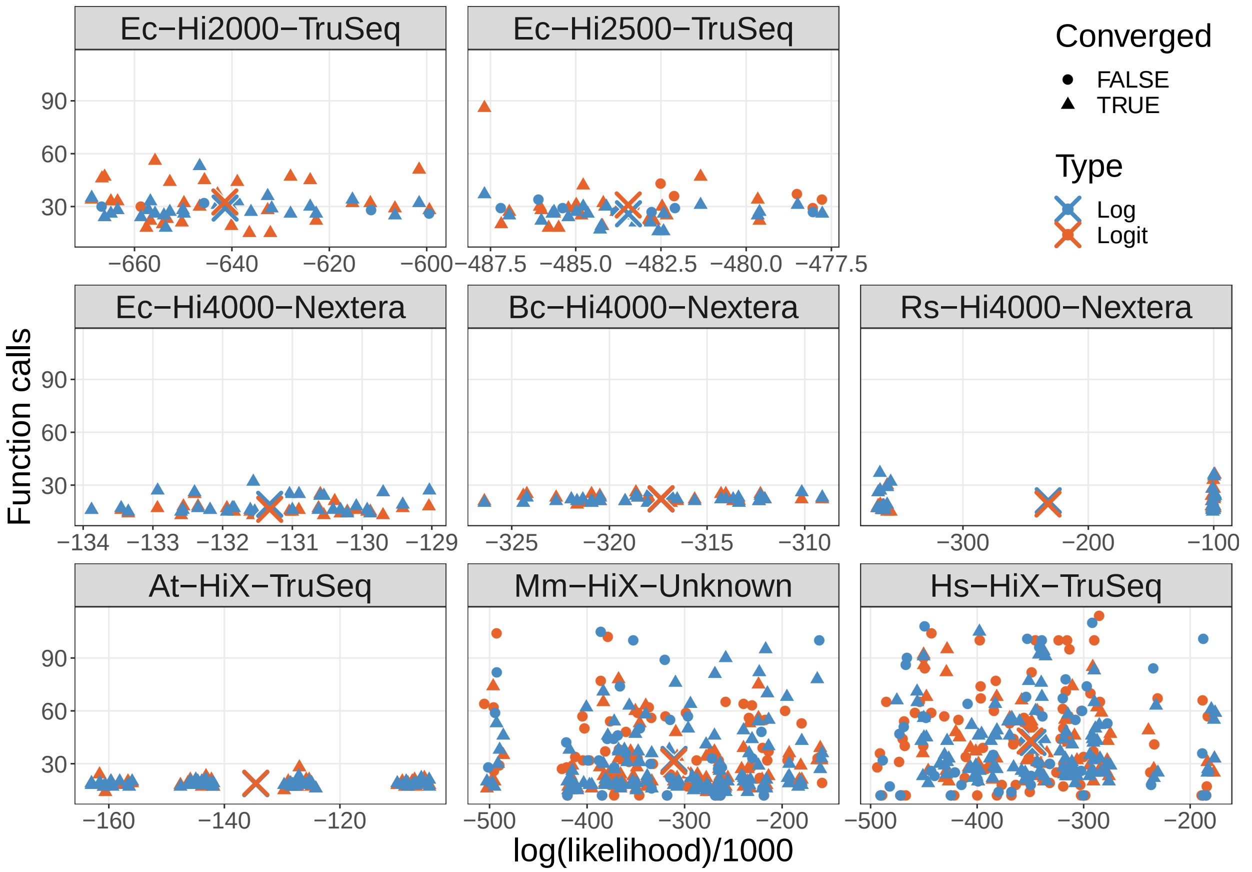


**Figure S21 Comparison of log and logit link function for the GC bias.** Dots and triangles are individual fits. Crosses are the mean value for that dataset.

### Supplementary Tables

**Table S1** Used simulator versions.

| Simulator | Version |
| --- | --- |
| ReSeq | 1.0 |
| | ac75312d263efde27d3655150fae474a5fdbf7d6 |
| pIRS | 2.0.0 |
| | bee9b594f4d0e10580aae77ec411cecec4a77219 |
| ART | 2.5.8 |
| | MountRainier-2016-06-05 |
| NEAT | v2.0 |
| | cdb869a2451221ab57bffeffe50329cdd1467c2f |
| BEAR | 02274019ce7c2ac70c2f642368bc0682fb97446a |

**Table S2** Used program versions.

| Program | Version |
| --- | --- |
| bcl2fastq | v2.20.0.422 |
| bcftools [2] | 1.9 |
| bedtools [3] | v2.25.0 |
| bowtie2 [4] | 2.2.5 |
| bwa mem [5] | 0.7.13-r1126 |
| freebayes [6] | v1.3.1-dirty |
| GNU time | 1.7 |
| igv [7, 8] | 2.6.3 |
| jellyfish [9] | 2.0.0 |
| kmc [10] | 3.1.1 (2019-05-19) |
| pilon [11] | 1.21 |
| preqc [12] | 0.10.14 |
| quast [13] | 4.4 |
| samtools [14] | 1.9 |
| sga [15] | 0.10.14 |
| snakemake [16] | 3.5.5 |
| soap [17] | 2.21 |

### Supplementary Formulas

Here the exact likelihood and gradient formulas that are used in the program for the coverage model fit are derived, starting from the formulas and information in the main document. While the formulas from the main document are repeated, only new symbols will be explained.

#### Step 1 Poisson

Internally, the normalization $N$ is always part of the GC bias, because it is absorbed by it for the log-likelihood calculation during this step.

$$\begin{aligned}\mu_n &= N b_{GC}(GC_n) b_{start,n} b_{end,n}\\ &= b^N_{GC} b_{start,n} b_{end,n}\\ &= b^N_{GC}\hat{\mu}_n\end{aligned}$$

$$L_P = \prod_n \frac{\mu_n^{k_n}}{k_n!} e^{-\mu_n}$$

$$\begin{aligned}\log(L_P) &= \sum_n \left[k_n \log(\mu_n) - \log(k_n!) - \mu_n\right]\\ &= \left[-\sum_n \log(k_n!)\right] + \sum_{GC}\sum_{n(GC)} \left[k_n \log(b^N_{GC}\hat{\mu}_n) - b^N_{GC}\hat{\mu}_n\right]\\ &= \left[-\sum_n \log(k_n!)\right] + \sum_{GC}\sum_{n(GC)} \left[k_n \log(b^N_{GC}) + k_n\log(\hat{\mu}_n) - b^N_{GC}\hat{\mu}_n\right]\\ &= \left[-\sum_n \log(k_n!)\right] + \sum_{GC}\left\{\log(b^N_{GC})\left[\sum_{n(GC)} k_n\right] + \left[\sum_{n(GC)} k_n\log(\hat{\mu}_n)\right] - b^N_{GC}\left[\sum_{n(GC)}\hat{\mu}_n\right]\right\}\end{aligned}$$

$$\frac{\partial \log(L_P)}{\partial b^N_{GC}} = \frac{1}{b^N_{GC}}\left[\sum_{n(GC)} k_n\right] - \left[\sum_{n(GC)}\hat{\mu}_n\right]$$

$$\begin{aligned}0 = \frac{\partial \log(L_P)}{\partial b^N_{GC}} \quad &\Leftrightarrow \quad \sum_{n(GC)}\hat{\mu}_n = \frac{1}{b^N_{GC}}\sum_{n(GC)} k_n\\ &\Leftrightarrow \quad b^N_{GC} = \frac{\sum_{n(GC)} k_n}{\sum_{n(GC)}\hat{\mu}_n} = \frac{\textcircled{1}}{\textcircled{2}}\end{aligned}$$

In the software, the variable names are ① gc_count_[gc] and ② gc_bias_sum[gc].second. When we use the calculated $b^N_{GC}$ in the likelihood, we always have perfect normalization, thus the normalization is absorbed by the GC bias.

$$\begin{aligned}\log(L_P) &= \left[-\sum_n \log(k_n!)\right] + \sum_{GC}\left\{\left[\sum_{n(GC)} k_n\log(\hat{\mu}_n)\right] + \left[\sum_{n(GC)} k_n\right]\log(b^N_{GC}) - \left[\sum_{n(GC)} k_n\right]\right\}\\ &= \textcircled{3} + \sum_{GC}\left\{\textcircled{4} + \textcircled{1}\log(b^N_{GC}) - \textcircled{1}\right\}\end{aligned}$$

③ loglike_poisson_base_, ④ gc_bias_sum[gc].first. This leaves the $b_{f,p}$ to be fitted, thus we need their gradients.

$$\frac{\partial b_{GC}^N}{\partial b_{f,p}} = -\frac{b_{GC}^N}{\sum_{n(GC)} \hat{\mu}_n} \sum_{n(GC)} \frac{\partial \hat{\mu}_n}{\partial b_{f,p}}$$

$$\begin{aligned}
\frac{\partial \log(L_P)}{\partial b_{f,p}} &= \sum_{GC} \left\{ \left[ \sum_{n(GC)} \frac{k_n}{\hat{\mu}_n} \frac{\partial \hat{\mu}_n}{\partial b_{f,p}} \right] + \frac{\sum_{n(GC)} k_n}{b_{GC}^N} \frac{\partial b_{GC}^N}{\partial b_{f,p}} \right\} \\
&= \sum_{GC} \left\{ \left[ \sum_{n(GC)} \frac{k_n}{\hat{\mu}_n} \frac{\partial \hat{\mu}_n}{\partial b_{f,p}} \right] - \left[ \sum_{n(GC)} \hat{\mu}_n \right] \frac{b_{GC}^N}{\sum_{n(GC)} \hat{\mu}_n} \sum_{n(GC)} \frac{\partial \hat{\mu}_n}{\partial b_{f,p}} \right\} \\
&= \left[ \sum_{n} \frac{k_n}{\hat{\mu}_n} \frac{\partial \hat{\mu}_n}{\partial b_{f,p}} \right] - \left\{ \sum_{GC} b_{GC}^N \sum_{n(GC)} \left[ \frac{\partial \hat{\mu}_n}{\partial b_{f,p}} \right] \right\} \\
&= ⑤ - \left\{ \sum_{GC} b_{GC}^N \sum_{n(GC)} ⑥ \right\}
\end{aligned}$$

⑤ grad_sur_[sur], ⑥ grad_gc_bias_sum[gc][sur]

For the sum version of the flanking bias, this leads to the following:

$$b_{start,n} = \frac{2}{1 + e^{-\sum_p b_{f(p,start),p}}}$$

$$\begin{aligned}
\frac{\partial b_{start,n}}{\partial b_{f,p}} &= -\frac{b_{start,n}}{1 + e^{-\sum_{\tilde{p}} b_{f(\tilde{p},start),\tilde{p}}}} e^{-\sum_{\tilde{p}} b_{f(\tilde{p},start),\tilde{p}}} \left( -\delta_{f(\tilde{p},start),f} \right) \\
&= b_{start,n} \delta_{f(\tilde{p},start),f} \frac{e^{-\sum_{\tilde{p}} b_{f(\tilde{p},start),\tilde{p}}} + 1 - 1}{1 + e^{-\sum_{\tilde{p}} b_{f(\tilde{p},start),\tilde{p}}}} \\
&= b_{start,n} \delta_{f(\tilde{p},start),f} \left( 1 - \frac{1}{1 + e^{-\sum_{\tilde{p}} b_{f(\tilde{p},start),\tilde{p}}}} \right) \\
&= b_{start,n} \delta_{f(\tilde{p},start),f} \left( 1 - \frac{b_{start,n}}{2} \right)
\end{aligned}$$

$$\begin{aligned}
\frac{\partial \hat{\mu}_n}{\partial b_{f,p}} &= \frac{\partial b_{start,n}}{\partial b_{f,p}} b_{end,n} + b_{start,n} \frac{\partial b_{end,n}}{\partial b_{f,p}} \\
&= b_{start,n} \delta_{f(\tilde{p},start),f} \left( 1 - \frac{b_{start,n}}{2} \right) b_{end,n} + b_{start,n} b_{end,n} \delta_{f(\tilde{p},end),f} \left( 1 - \frac{b_{end,n}}{2} \right) \\
&= \hat{\mu}_n \left[ \delta_{f(\tilde{p},start),f} \left( 1 - \frac{b_{start,n}}{2} \right) + \delta_{f(\tilde{p},end),f} \left( 1 - \frac{b_{end,n}}{2} \right) \right]
\end{aligned}$$

$$\begin{aligned}
\frac{\partial \log(L_{PS})}{\partial b_{f,p}} = & \left[ \sum_n k_n \left[ \delta_{f(\tilde{p},start),f} \left( 1 - \frac{b_{start,n}}{2} \right) + \delta_{f(\tilde{p},end),f} \left( 1 - \frac{b_{end,n}}{2} \right) \right] \right] \\
& - \left\{ \sum_{GC} b_{GC}^N \left[ \sum_{n(GC)} \hat{\mu}_n \left[ \delta_{f(\tilde{p},start),f} \left( 1 - \frac{b_{start,n}}{2} \right) + \delta_{f(\tilde{p},end),f} \left( 1 - \frac{b_{end,n}}{2} \right) \right] \right] \right\}
\end{aligned}$$

Inspired by biases of the four nucleotides at a position not being independent, we do not directly use $b_{f,p}$ as fit parameters, but $\tilde{b}_{f,p}$. However, we still have 4 parameters per position, because our attempts to reduce this to 3 did not improve the fitting.

$$b_{f,p} = \tilde{b}_{f,p} - \frac{\sum_{\tilde{s}} \tilde{b}_{\tilde{s},p}}{4} + \delta_{p0}\tilde{b}_{shift}$$

$\tilde{b}_{shift}$ is an additional parameter that can shift the spread of different $\sum_p b_{f(p,start),p}$ to an ideal range for the inverse logit transformation.

$$\begin{aligned}
\frac{\partial \log(L_{PS})}{\partial \tilde{b}_{f,p}} &= \sum_{\tilde{s}} \frac{\partial b_{\tilde{s},p}}{\partial \tilde{b}_{f,p}} \frac{\partial \log(L_P)}{\partial b_{\tilde{s},p}} \\
&= \frac{\partial \log(L_P)}{\partial b_{f,p}} - \frac{\sum_{\tilde{s}} \frac{\partial \log(L_P)}{\partial b_{\tilde{s},p}}}{4}
\end{aligned}$$

$$\begin{aligned}
\frac{\partial \log(L_{PS})}{\partial \tilde{b}_{shift}} &= \sum_{\tilde{s}} \frac{\partial b_{\tilde{s},0}}{\partial \tilde{b}_{shift}} \frac{\partial \log(L_P)}{\partial b_{\tilde{s},0}} \\
&= \sum_{\tilde{s}} \frac{\partial \log(L_P)}{\partial b_{\tilde{s},0}}
\end{aligned}$$

For the product version of the flanking bias, the likelihood and gradients are the following.

$$b_{start,n} = \prod_p b_{f(p,start),p}$$

$$\frac{\partial b_{start,n}}{\partial b_{f,p}} = \delta_{f(p,start),f} \frac{b_{start,n}}{b_{f,p}}$$

$$\begin{aligned}
\frac{\partial \hat{\mu}_n}{\partial b_{f,p}} &= \frac{\partial b_{start,n}}{\partial b_{f,p}} b_{end,n} + b_{start,n} \frac{\partial b_{end,n}}{\partial b_{f,p}} \\
&= \delta_{f(p,start),f} \frac{b_{start,n}}{b_{f,p}} b_{end,n} + b_{start,n} \delta_{f(p,end),f} \frac{b_{end,n}}{b_{f,p}} \\
&= \frac{\hat{\mu}_n}{b_{f,p}} \left(\delta_{f(p,start),f} + \delta_{f(p,end),f}\right)
\end{aligned}$$

$$\begin{aligned}
\frac{\partial \log(L_{PP})}{\partial b_{f,p}} = & \left[\sum_n \frac{k_n}{b_{f,p}} \left(\delta_{f(\tilde{p},start),f} + \delta_{f(\tilde{p},end),f}\right)\right] \\
& - \left\{\sum_{GC} b_{GC}^N \left[\sum_{n(GC)} \frac{\hat{\mu}_n}{b_{f,p}} \left(\delta_{f(\tilde{p},start),f} + \delta_{f(\tilde{p},end),f}\right)\right]\right\}
\end{aligned}$$

Similar to the sum version, the real fit parameters are $\tilde{b}_{f,p}$.

$$b_{f,p} = \frac{4\tilde{b}_{f,p}}{\sum_{\tilde{s}} \tilde{b}_{\tilde{s},p}}$$

$$\begin{aligned}
\frac{\partial b_{s,p}}{\partial \tilde{b}_{f,p}} &= 4\frac{\delta_{s,f}\left[\sum_{\tilde{s}}\tilde{b}_{\tilde{s},p}\right]-\tilde{b}_{s,p}}{\left[\sum_{\tilde{s}}\tilde{b}_{\tilde{s},p}\right]^2} \\
&= \frac{4\delta_{s,f}-\frac{4\tilde{b}_{s,p}}{\sum_{\tilde{s}}\tilde{b}_{\tilde{s},p}}}{\sum_{\tilde{s}}\tilde{b}_{\tilde{s},p}} \\
&= \frac{4\delta_{s,f}-b_{s,p}}{\sum_{\tilde{s}}\tilde{b}_{\tilde{s},p}}
\end{aligned}$$

$$\begin{aligned}
\frac{\partial \log(L_{PP})}{\partial \tilde{b}_{f,p}} &= \sum_s \frac{\partial b_{s,p}}{\partial \tilde{b}_{f,p}}\frac{\partial \log(L_P)}{\partial b_{s,p}} \\
&= \frac{4\frac{\partial \log(L_P)}{\partial b_{f,p}}-\left[\sum_s b_{s,p}\frac{\partial \log(L_P)}{\partial b_{s,p}}\right]}{\sum_{\tilde{s}}\tilde{b}_{\tilde{s},p}}
\end{aligned}$$

Step 2 GC bias spline

$$b_{GC,spline}(GC) = c_1(GC) + c_2(GC)x(GC) + c_3(GC)x^2(GC) + c_4(GC)x^3(GC)$$

$$c_l(GC) = \sum_{j=1}^{6} t_{l,j}(GC)s_j$$

$s_j$ are the six spline parameters, $t_{l,j}(GC)$ are coefficients calculated from the knot positions and $x(GC)$ is the distance to the last knot.

$$\frac{\partial b_{GC,spline}(GC)}{\partial s_{\tilde{j}}} = t_{1,\tilde{j}}(GC) + t_{2,\tilde{j}}(GC)x(GC) + t_{3,\tilde{j}}(GC)x^2(GC) + t_{4,\tilde{j}}(GC)x^3(GC)$$

The likelihood is the Poisson likelihood without the substitution of $b^N_{GC}$

$$\begin{aligned}
\log(L_B) &= \left[-\sum_n \log(k_n!)\right] + \sum_{GC}\left\{\left[\sum_{n(GC)} k_n \log(\hat{\mu}_n)\right] + \left[\sum_{n(GC)} k_n\right]\log(b^N_{GC}) - b^N_{GC}\left[\sum_{n(GC)}\hat{\mu}_n\right]\right\} \\
&= \textcircled{3} + \sum_{GC}\left\{\textcircled{4} + \textcircled{1}\log(b^N_{GC}) - b^N_{GC}\textcircled{2}\right\}
\end{aligned}$$

$$\begin{aligned}
\frac{\partial \log(L_B)}{\partial s_{\tilde{j}}} &= \frac{\partial \log(L_B)}{\partial b^N_{GC}}\frac{\partial b^N_{GC}}{\partial b_{GC,spline}}\frac{\partial b_{GC,spline}}{\partial s_{\tilde{j}}} \\
&= \sum_{GC}\left\{\left(\frac{1}{b^N_{GC}}\left[\sum_{n(GC)} k_n\right] - \left[\sum_{n(GC)}\hat{\mu}_n\right]\right)\frac{\partial b^N_{GC}}{\partial b_{GC,spline}}\frac{\partial b_{GC,spline}}{\partial s_{\tilde{j}}}\right\} \\
&= \sum_{GC}\left\{\left(\left[\sum_{n(GC)} k_n\right] - b^N_{GC}\left[\sum_{n(GC)}\hat{\mu}_n\right]\right)\left(\frac{1}{b^N_{GC}}\frac{\partial b^N_{GC}}{\partial b_{GC,spline}}\right)\frac{\partial b_{GC,spline}}{\partial s_{\tilde{j}}}\right\}
\end{aligned}$$

For the exponential version:

$$b^N_{GC} = Ne^{b_{GC,spline}}$$

$$\frac{\partial b_{GC}^N}{\partial b_{GC,spline}} = b_{GC}^N$$

For the inverse logit version:

$$b_{GC}^N = \frac{2N}{1 + e^{-b_{GC,spline}}}$$

$$\begin{aligned}
\frac{\partial b_{GC}^N}{\partial b_{GC,spline}} &= -2N \frac{-e^{-b_{GC,spline}}}{(1 + e^{-b_{GC,spline}})^2} \\
&= b_{GC}^N \frac{e^{-b_{GC,spline}}}{1 + e^{-b_{GC,spline}}} \\
&= b_{GC}^N \frac{1}{1 + e^{b_{GC,spline}}}
\end{aligned}$$

Step 3 Negative binomial

$$L_{NB} = \prod_n \left\{ \binom{k_n + r_n - 1}{k_n} \left(1 - \frac{\mu_n}{\mu_n + r_n}\right)^{r_n} \left(\frac{\mu_n}{\mu_n + r_n}\right)^{k_n} \right\}$$

$$r_n = \frac{\mu_n}{\alpha + \beta\mu_n}$$

$$\begin{aligned}
\frac{\partial r_n}{\partial \mu_n} &= \frac{\alpha + \beta\mu_n - \mu_n\beta}{(\alpha + \beta\mu)^2} \\
&= \frac{\alpha}{(\alpha + \beta\mu)^2} \\
&= \frac{\alpha r_n^2}{\mu_n^2}
\end{aligned}$$

$$\begin{aligned}
\log(L_{NB}) &= \sum_n \left\{ \log\left( \prod_{i=1}^{k_n} \frac{k_n + r_n - 1 + 1 - i}{i} \right) + r_n \log\left(1 - \frac{\mu_n}{\mu_n + r_n}\right) + k_n \log\left(\frac{\mu_n}{\mu_n + r_n}\right) \right\} \\
&= \sum_n \left\{ \left[ \sum_{i=1}^{k_n} \log\left(\frac{k_n + r_n - i}{i}\right) \right] + r_n \log\left(\frac{r_n}{\mu_n + r_n}\right) + k_n \log\left(\frac{\mu_n}{\mu_n + r_n}\right) \right\} \\
&= \sum_n \left\{ \left[ \sum_{i=1}^{k_n} \log\left(k_n + r_n - (k_n - i + 1)\right) \right] - \left[ \sum_{i=1}^{k_n} \log(i) \right] + r_n \log\left(\frac{r_n}{\mu_n + r_n}\right) + k_n \log\left(\frac{\mu_n}{\mu_n + r_n}\right) \right\} \\
&= \sum_n \left\{ \left[ \sum_{i=1}^{k_n} \log\left(\frac{r_n + i - 1}{i}\right) \right] + r_n \log\left(\frac{r_n}{\mu_n + r_n}\right) + k_n \log\left(\frac{\mu_n}{\mu_n + r_n}\right) \right\} \\
&= \sum_n \left\{ k_n \log\left(\frac{\mu_n}{\mu_n + r_n}\right) + r_n \log\left(\frac{r_n}{\mu_n + r_n}\right) + \left[ \sum_{i=1}^{k_n} \log\left(\frac{r_n + (i-1)}{i}\right) \right] \right\}
\end{aligned}$$

$$
\begin{aligned}
\frac{\partial \log(L_{NB})}{\partial \mu_n} = \quad & \textstyle\sum_n \Bigg\{ \quad \left[ k_n \frac{\mu_n + r_n}{\mu_n} \frac{(\mu_n + r_n) - \mu_n \left(1 + \frac{\partial r_n}{\partial \mu_n}\right)}{(\mu_n + r_n)^2} \right] \\
& \quad + \left[ \frac{\partial r_n}{\partial \mu_n} \log\left(\frac{r_n}{\mu_n + r_n}\right) + r_n \frac{\mu_n + r_n}{r_n} \frac{\frac{\partial r_n}{\partial \mu_n}(\mu_n + r_n) - r_n \left(1 + \frac{\partial r_n}{\partial \mu_n}\right)}{(\mu_n + r_n)^2} \right] \\
& \quad + \left[ \sum_{i=1}^{k_n} \frac{i}{r_n + i - 1} \frac{\frac{\partial r_n}{\partial \mu_n}}{i} \right] \Bigg\} \\
= \quad & \textstyle\sum_n \Bigg\{ \quad \left[ k_n \frac{r_n - \mu_n \frac{\partial r_n}{\partial \mu_n}}{\mu_n(\mu_n + r_n)} \right] + \left[ \frac{\mu_n \frac{\partial r_n}{\partial \mu_n} - r_n}{\mu_n + r_n} + \frac{\partial r_n}{\partial \mu_n} \log\left(\frac{r_n}{\mu_n + r_n}\right) \right] + \left[ \sum_{i=1}^{k_n} \frac{\frac{\partial r_n}{\partial \mu_n}}{r_n + i - 1} \right] \Bigg\} \\
= \quad & \textstyle\sum_n \Bigg\{ \quad \frac{(k_n - \mu_n)\left(r_n - \mu_n \frac{\partial r_n}{\partial \mu_n}\right)}{\mu_n(\mu_n + r_n)} + \frac{\partial r_n}{\partial \mu_n} \log\left(\frac{r_n}{\mu_n + r_n}\right) + \left[ \frac{\partial r_n}{\partial \mu_n} \sum_{i=1}^{k_n} \frac{1}{r_n + i - 1} \right] \Bigg\} \\
= \quad & \textstyle\sum_n \Bigg\{ \quad \frac{(k_n - \mu_n)\left(r_n - \mu_n \frac{\alpha r_n^2}{\mu_n^2}\right)}{\mu_n(\mu_n + r_n)} + \frac{\alpha r_n^2}{\mu_n^2} \log\left(\frac{r_n}{\mu_n + r_n}\right) + \left[ \frac{\alpha r_n^2}{\mu_n^2} \sum_{i=1}^{k_n} \frac{1}{r_n + i - 1} \right] \Bigg\} \\
= \quad & \textstyle\sum_n \Bigg\{ \quad \frac{r_n(k_n - \mu_n)}{\mu_n(\mu_n + r_n)} \left(1 - \frac{\alpha r_n}{\mu_n}\right) + \left[ \log\left(\frac{r_n}{\mu_n + r_n}\right) + \sum_{i=1}^{k_n} \frac{1}{r_n + i - 1} \right] r_n^2 \frac{\alpha}{\mu_n^2} \Bigg\} \\
= \quad & \textstyle\sum_n \Bigg\{ \quad \left( \frac{r_n(k_n - \mu_n)}{\mu_n + r_n} \left(1 - \frac{\alpha r_n}{\mu_n}\right) + \left[ \log\left(\frac{1}{\frac{\mu_n}{r_n} + 1}\right) + \sum_{i=1}^{k_n} \frac{1}{r_n + (i-1)} \right] r_n^2 \frac{\alpha}{\mu_n} \right) \frac{1}{\mu_n} \Bigg\}
\end{aligned}
$$

$$
\begin{aligned}
\frac{\partial \log(L_{NB})}{\partial s_{\tilde{j}}} &= \frac{\partial \log(L_B)}{\partial \mu_n} \frac{\partial \mu_n}{\partial b_{GC}^N} \frac{\partial b_{GC}^N}{\partial b_{GC,spline}} \frac{\partial b_{GC,spline}}{\partial s_{\tilde{j}}} \\
&= \left[ \frac{\partial \log(L_B)}{\partial \mu_n} \mu_n \right] \left[ \frac{1}{b_{GC}^N} \frac{\partial b_{GC}^N}{\partial b_{GC,spline}} \right] \frac{\partial b_{GC,spline}}{\partial s_{\tilde{j}}}
\end{aligned}
$$

$$
\frac{\partial \log(L_{NB})}{\partial b_{f,p}} = \left[ \frac{\partial \log(L_B)}{\partial \mu_n} \mu_n \right] \left[ \frac{1}{\mu_n} \frac{\partial \mu_n}{\partial b_{f,p}} \right]
$$

$$
\begin{aligned}
\frac{\partial \log(L_{NB})}{\partial r_n} = \quad & \textstyle\sum_n \Bigg\{ \quad \left[ k_n \frac{\mu_n + r_n}{\mu_n} \frac{-\mu_n}{(\mu_n + r_n)^2} \right] \\
& \quad + \left[ \log\left(\frac{r_n}{\mu_n + r_n}\right) + r_n \frac{\mu_n + r_n}{r_n} \frac{(\mu_n + r_n) - r_n}{(\mu_n + r_n)^2} \right] \\
& \quad + \left[ \sum_{i=1}^{k_n} \frac{i}{r_n + i - 1} \frac{1}{i} \right] \Bigg\} \\
= \quad & \textstyle\sum_n \Bigg\{ \quad \left[ \frac{-k_n}{\mu_n + r_n} \right] + \left[ \frac{\mu_n}{\mu_n + r_n} + \log\left(\frac{r_n}{\mu_n + r_n}\right) \right] + \left[ \sum_{i=1}^{k_n} \frac{1}{r_n + i - 1} \right] \Bigg\} \\
= \quad & \textstyle\sum_n \Bigg\{ \quad \frac{\mu_n - k_n}{\mu_n + r_n} + \log\left(\frac{1}{\frac{\mu_n}{r_n} + 1}\right) + \left[ \sum_{i=1}^{k_n} \frac{1}{r_n + (i-1)} \right] \Bigg\}
\end{aligned}
$$

$$
\frac{\partial r_n}{\partial \alpha} = \frac{-\mu_n}{(\alpha + \beta\mu)^2} = \frac{-r_n^2}{\mu_n}
$$

$$\begin{aligned}\frac{\partial \log(L_{NB})}{\partial \alpha} &= \frac{\partial \log(L_B)}{\partial r_n}\frac{\partial r_n}{\partial \alpha} \\ &= \left\{\frac{r_n^2}{\mu_n}\frac{k_n-\mu_n}{\mu_n+r_n} - \left[\log\left(\frac{1}{\frac{\mu_n}{r_n}+1}\right) + \sum_{i=1}^{k_n}\frac{1}{r_n+(i-1)}\right]\frac{r_n^2}{\mu_n}\right\}\end{aligned}$$

$$\frac{\partial r_n}{\partial \beta} = \frac{-\mu_n^2}{(\alpha+\beta\mu)^2} = -r_n^2$$

$$\begin{aligned}\frac{\partial \log(L_{NB})}{\partial \beta} &= \frac{\partial \log(L_B)}{\partial r_n}\frac{\partial r_n}{\partial \beta} \\ &= \left\{r_n^2\frac{k_n-\mu_n}{\mu_n+r_n} - \left[\log\left(\frac{1}{\frac{\mu_n}{r_n}+1}\right) + \sum_{i=1}^{k_n}\frac{1}{r_n+(i-1)}\right]r_n^2\right\}\end{aligned}$$
