## Supplementary material for "ReSeq simulates realistic Illumina high-throughput sequencing data": QUAST assembly report

| | sga-contigs |
|---|---|
| # contigs (>= 0 bp) | 61992 |
| # contigs (>= 1000 bp) | 1872 |
| # contigs (>= 5000 bp) | 306 |
| # contigs (>= 10000 bp) | 137 |
| # contigs (>= 25000 bp) | 14 |
| # contigs (>= 50000 bp) | 0 |
| Total length (>= 0 bp) | 16229865 |
| Total length (>= 1000 bp) | 6216155 |
| Total length (>= 5000 bp) | 3477799 |
| Total length (>= 10000 bp) | 2265808 |
| Total length (>= 25000 bp) | 465690 |
| Total length (>= 50000 bp) | 0 |
| # contigs | 61992 |
| Largest contig | 47116 |
| Total length | 16229865 |
| Reference length | 4616724 |
| GC (%) | 48.23 |
| Reference GC (%) | 50.80 |
| N50 | 402 |
| NG50 | 9781 |
| N75 | 127 |
| NG75 | 5017 |
| L50 | 4813 |
| LG50 | 142 |
| L75 | 29805 |
| LG75 | 303 |
| # misassemblies | 3 |
| # misassembled contigs | 3 |
| Misassembled contigs length | 681 |
| # local misassemblies | 4 |
| # unaligned mis. contigs | 0 |
| # unaligned contigs | 8053 + 0 part |
| Unaligned length | 4567968 |
| Genome fraction (%) | 98.928 |
| Duplication ratio | 2.553 |
| # N's per 100 kbp | 0.00 |
| # mismatches per 100 kbp | 6.59 |
| # indels per 100 kbp | 0.26 |
| Largest alignment | 47116 |
| Total aligned length | 11640635 |
| NA50 | 126 |
| NGA50 | 9781 |
| NGA75 | 5017 |
| LA50 | 25821 |
| LGA50 | 142 |
| LGA75 | 303 |

#### Misassemblies report

| | sga-contigs |
|---|---|
| # misassemblies | 3 |
| # relocations | 3 |
| # translocations | 0 |
| # inversions | 0 |
| # misassembled contigs | 3 |
| Misassembled contigs length | 681 |
| # local misassemblies | 4 |
| # unaligned mis. contigs | 0 |
| # mismatches | 301 |
| # indels | 12 |
| # indels (<= 5 bp) | 12 |
| # indels (> 5 bp) | 0 |
| Indels length | 12 |

### Unaligned report

| | sga-contigs |
|---|---|
| # fully unaligned contigs | 8053 |
| Fully unaligned length | 4567968 |
| # partially unaligned contigs | 0 |
| Partially unaligned length | 0 |
| # N's | 0 |

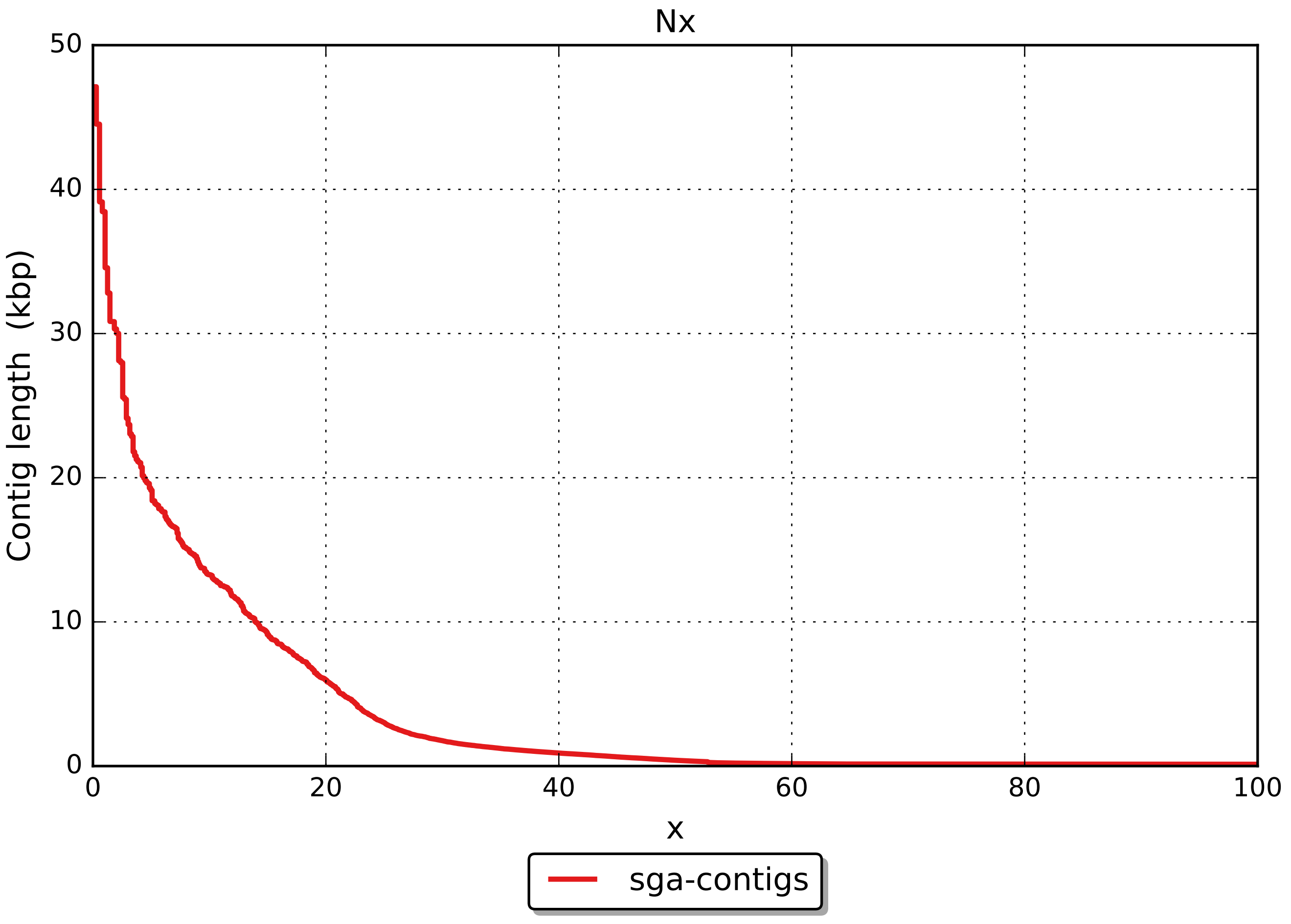
Nx
Contig length (kbp)
50
40
30
20
10
0
0
20
40
60
80
100
x
sga-contigs

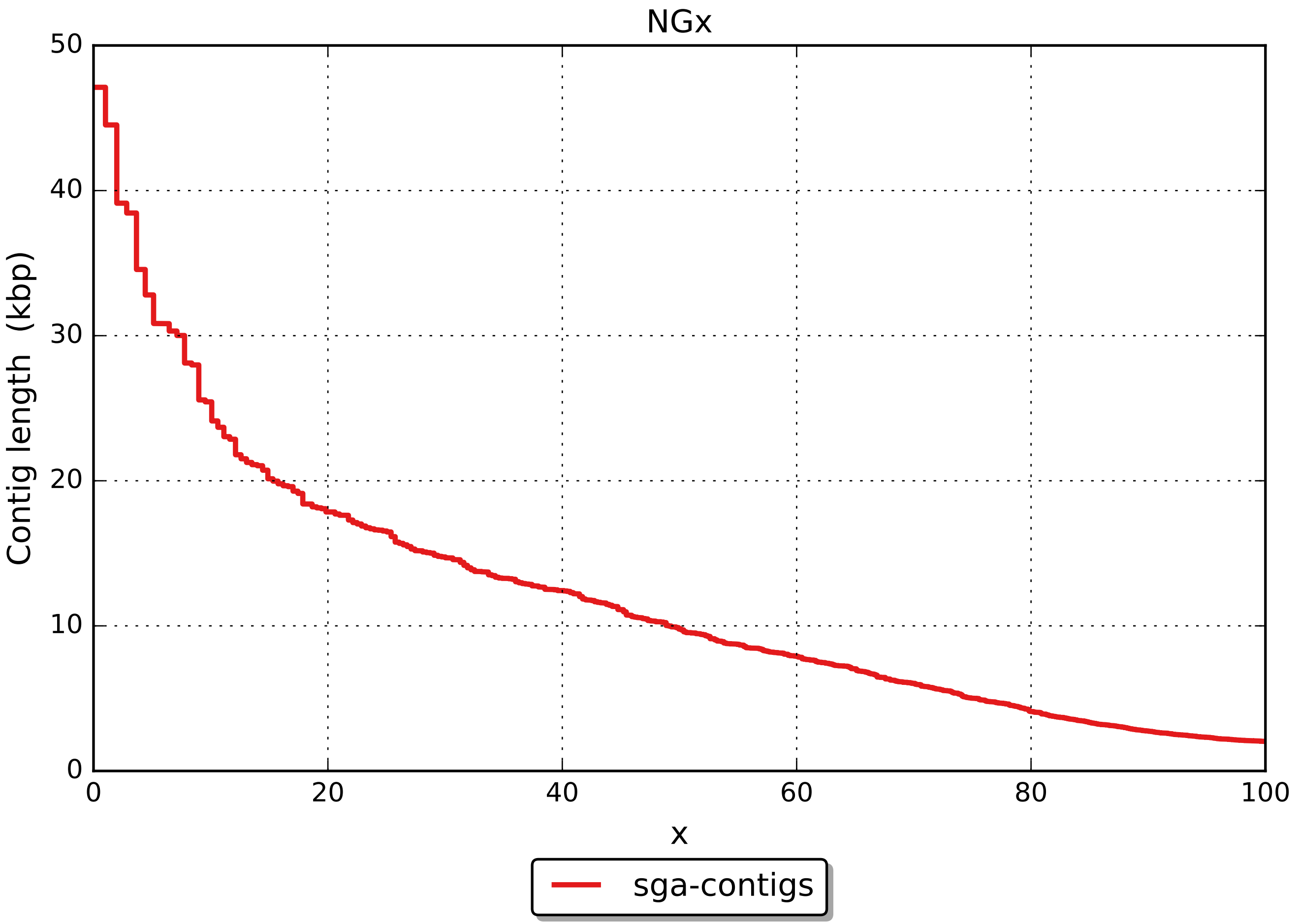
NGx
Contig length (kbp)
0
10
20
30
40
50
0
20
40
60
80
100
x
sga-contigs

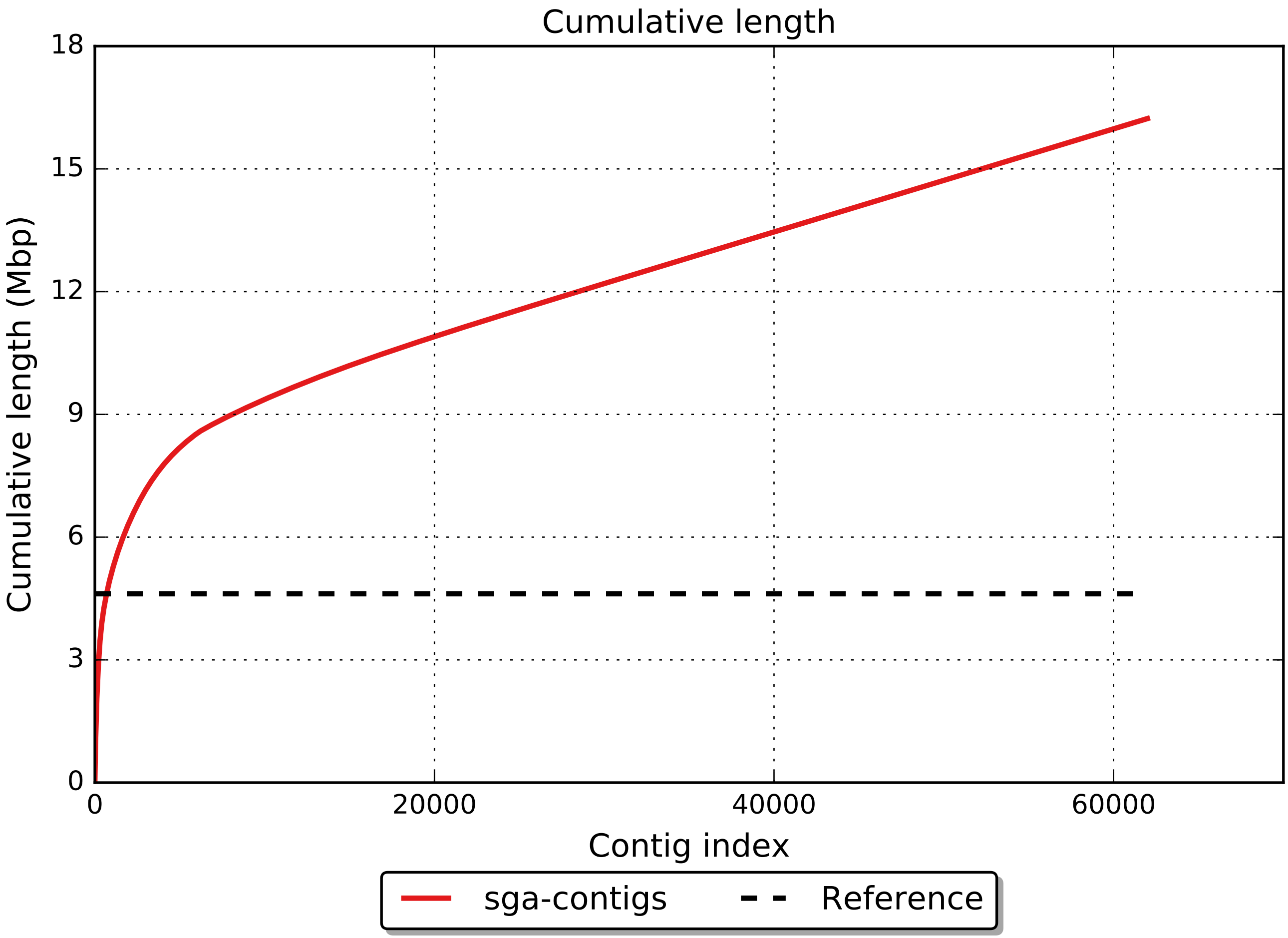

Cumulative length
Cumulative length (Mbp)
0
3
6
9
12
15
18
0
20000
40000
60000
Contig index
sga-contigs
Reference

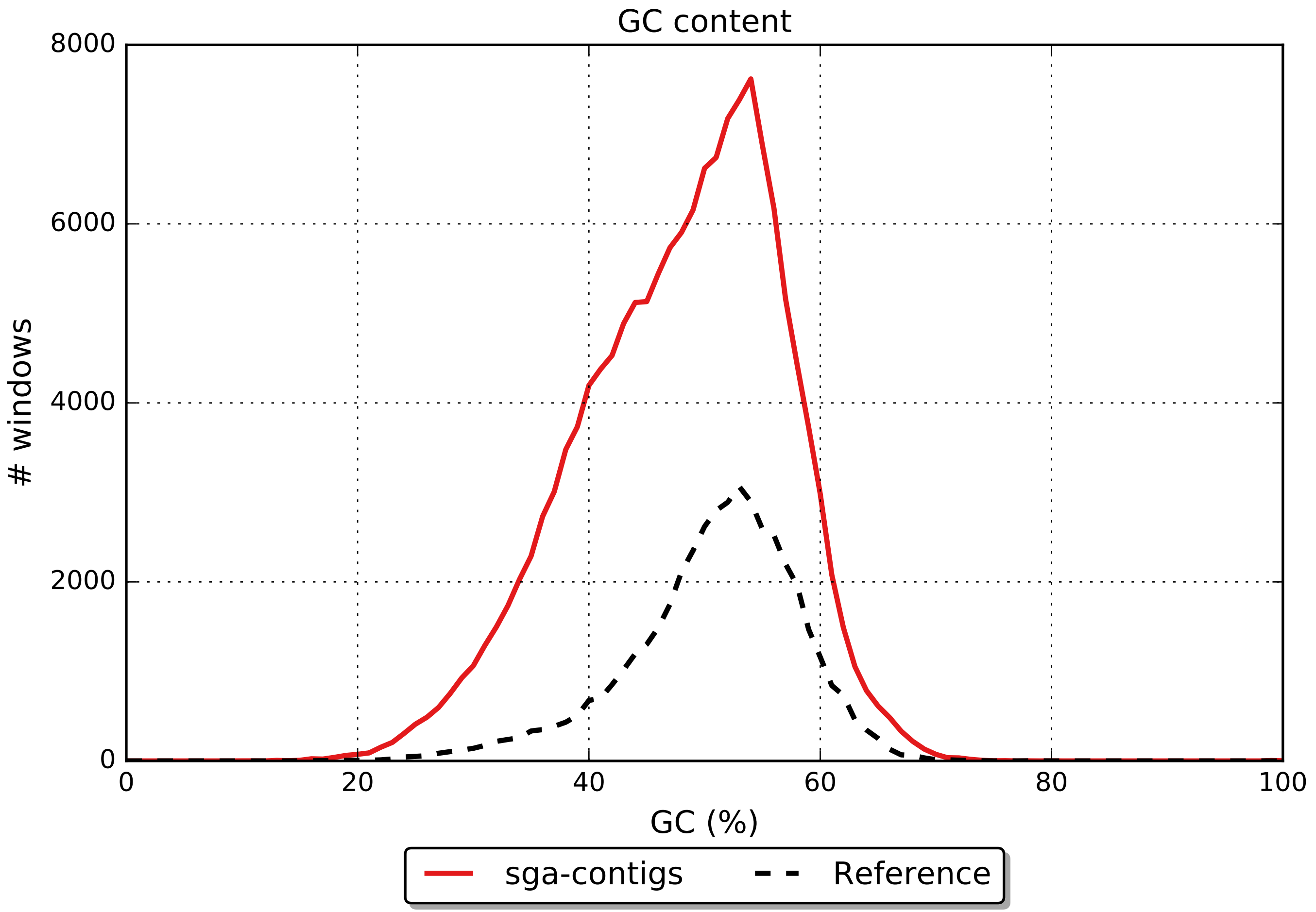
GC content
8000
6000
4000
2000
0
### windows
0
20
40
60
80
100
GC (%)
sga-contigs
Reference

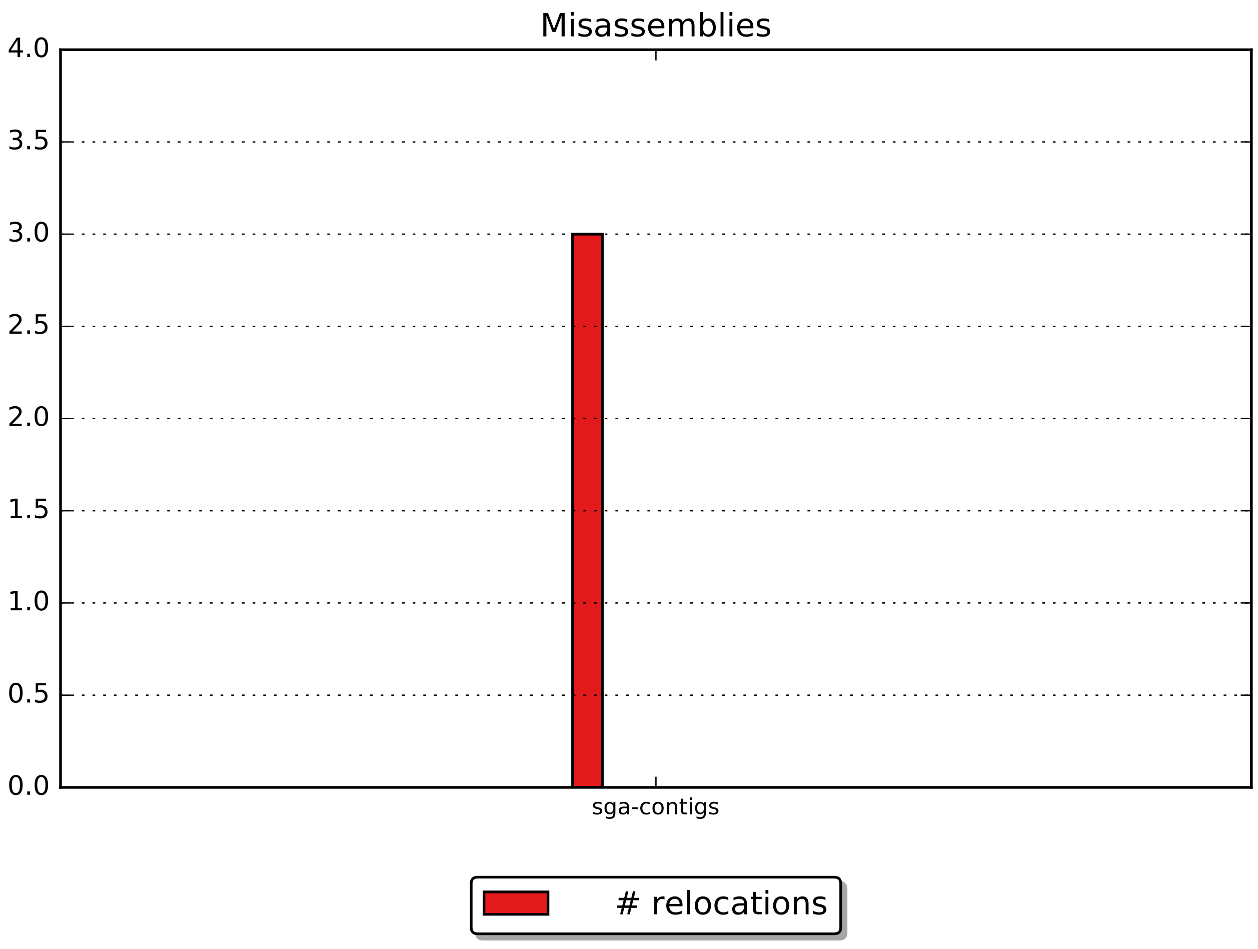
Misassemblies
4.0
3.5
3.0
2.5
2.0
1.5
1.0
0.5
0.0
sga-contigs
### relocations

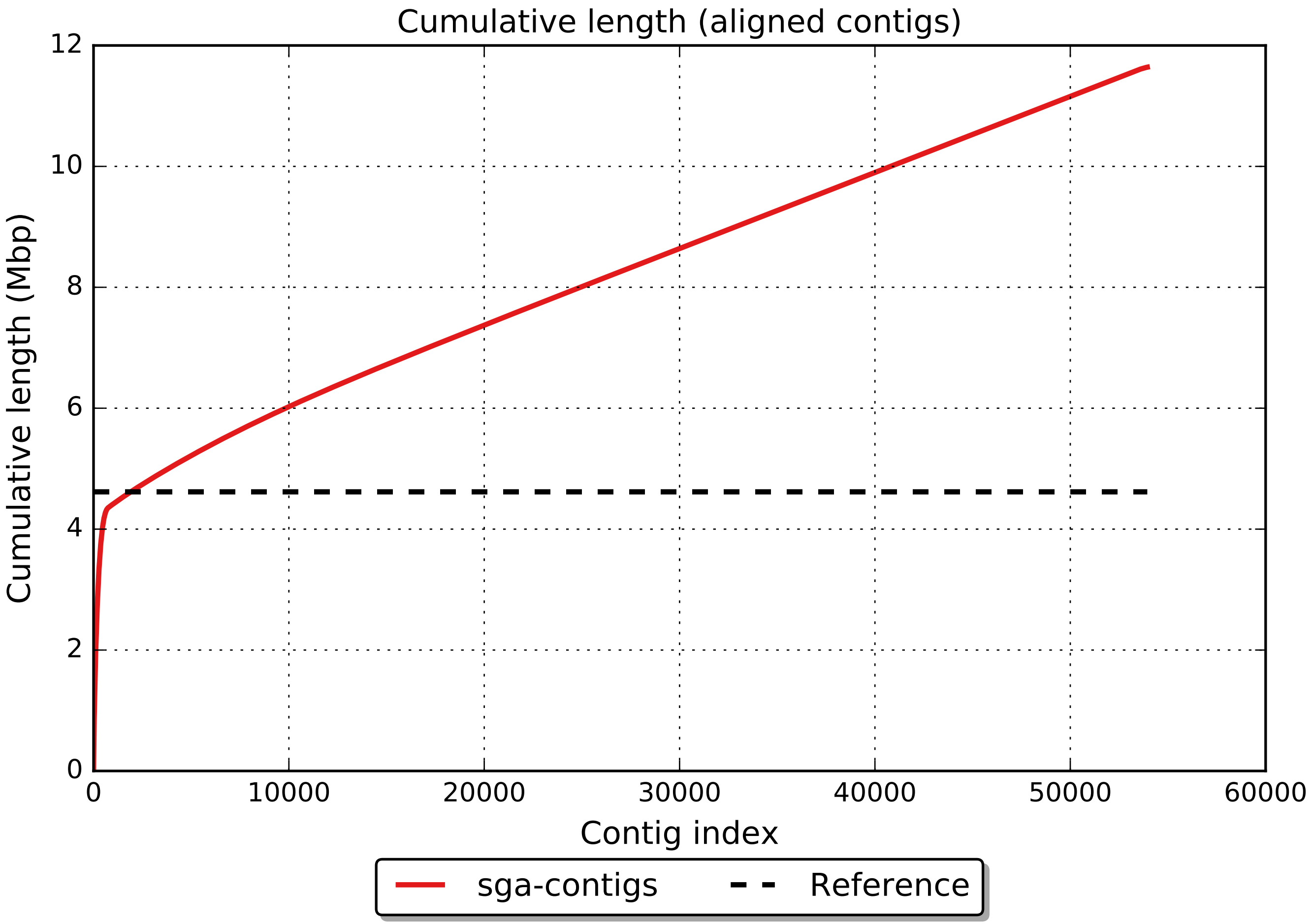

Cumulative length (aligned contigs)
Cumulative length (Mbp)
0
2
4
6
8
10
12
0
10000
20000
30000
40000
50000
60000
Contig index
sga-contigs
Reference

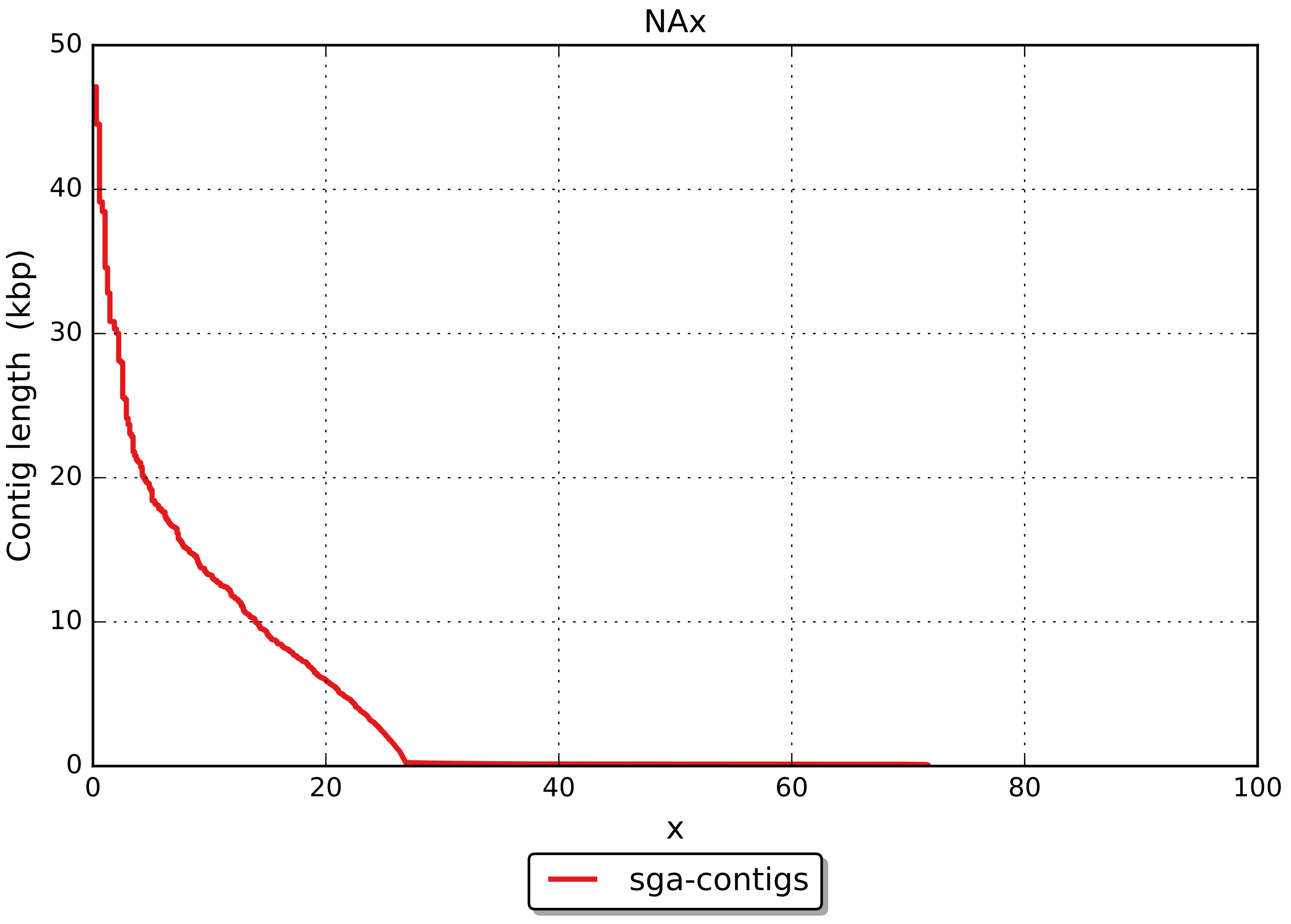

NAx
50
40
30
20
10
0
0
20
40
60
80
100
Contig length (kbp)
x
sga-contigs

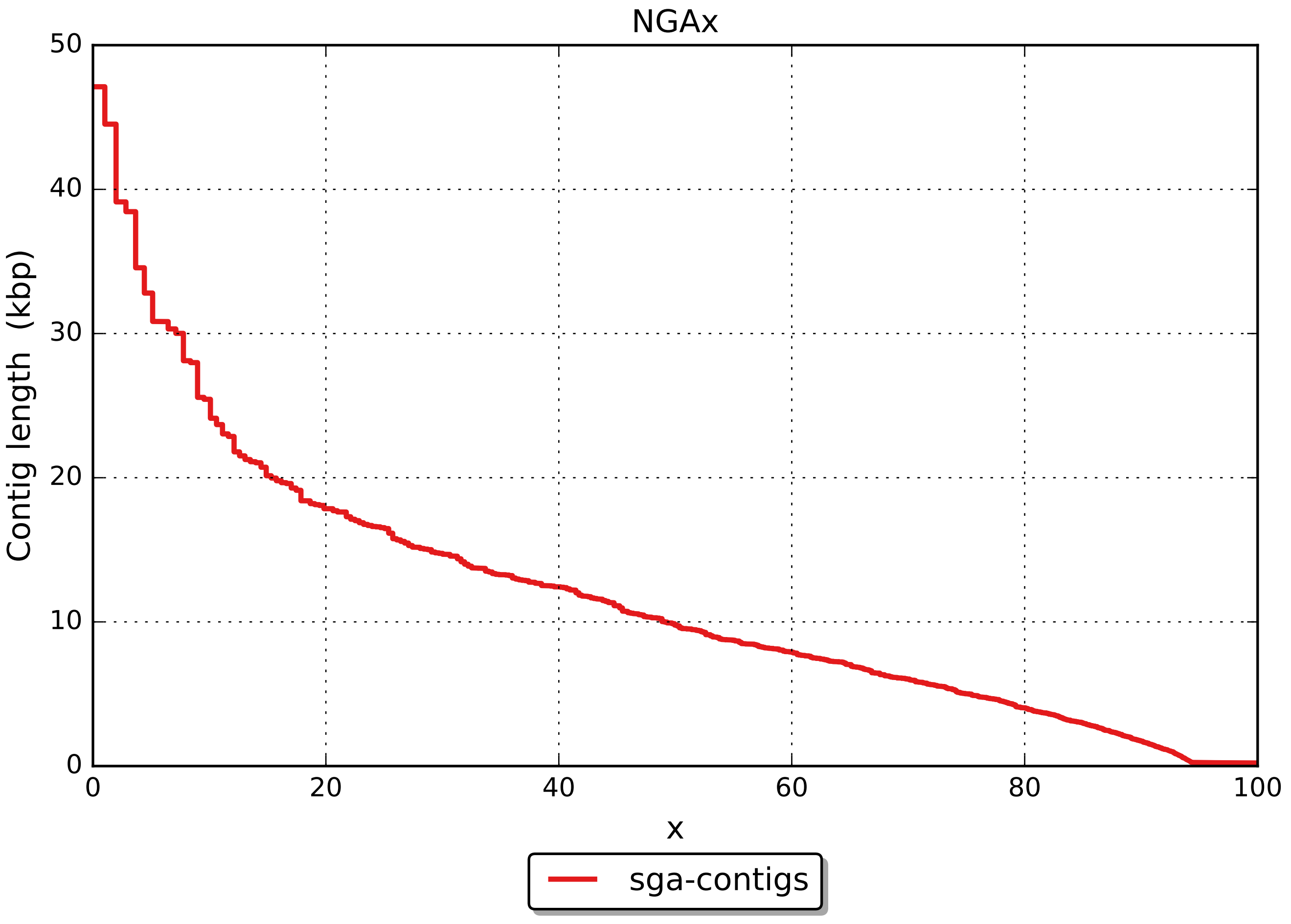
NGAx
Contig length (kbp)
50
40
30
20
10
0
0
20
40
60
80
100
x
sga-contigs
