## Supplementary material for "ReSeq simulates realistic Illumina high-throughput sequencing data": ReSeq Manual

More realistic simulator for genomic DNA sequences from Illumina machines that achieves a similar k-mer spectrum as the original sequences.

### Table of Contents


- Abstract
- Requirements
- Installation
- Quick start examples
- Parameter
- File Formats
- Included libraries
- Publication


### Abstract


Even though sequencing biases and errors have been deeply researched to adequately account for them, comparison studies, e.g. for error correction, assembly or variant calling, face the problem that synthetic datasets resemble the real output of high-throughput sequencers only in very limited ways, resulting in optimistic estimated performance of programs run on simulated data compared to real data. Therefore, comparison studies are often based on real data. However, this approach has its own difficulties, since the ground truth is unknown and can only be estimated, which introduces its own biases and circularity towards easy solutions and the methods used.

**Re**al **Seq**uence Reproducer shortens the gap between simulated and real data evaluations by adequately reproducing key statistics of real data, like the coverage profile, systematic errors and the k-mer spectrum. When these characteristics are translated into new synthetic computational experiments (i.e. simulated data), the performance can be more accurately estimated. Combining our simulator and real data gives two valuable perspectives on the performance of tools to minimize biases.


### Requirements

| Requirement | Ubuntu/Debian | CentOS | Manual installation | Comments | Tested version |
|---|---|---|---|---|---|
| Linux system | | | | | |
| Compiler supporting C++14 | `sudo apt install build-essential` | `sudo yum install gcc gcc-c++ glibc-devel make` | | On CentOs 7 the g++ compiler is too old to support C++14 so you need to additionally to the yum command install a newer version following for example this guide (https://linuxhostsupport.com/blog/how-to-install-gcc-on-centos-7/). If the standard install path `usr/local/` was used, afterwards the `CXX` variable has to be set to `/usr/local/bin/c++`, the `CC` variable to `/usr/local/bin/gcc`, `/usr/local/lib64/` has to be added to your `LD_LIBRARY_PATH` and `/usr/local/bin` to your `PATH` before installing boost. | 7.2.1 20171019 |
| ZLIB | `sudo apt-get install zlib1g-dev` | `sudo yum install zlib-devel` | | | |
| BZip2 | `sudo apt-get install libbz2-dev` | `sudo yum install bzip2-devel` | | | |
| Python 2 | | | | Preinstalled, but through something like Anaconda the default might be python3 | 2.7 |
| Python libraries | `sudo apt-get install python-dev` | `sudo yum install python-devel` | | | |
| Git | `sudo apt-get install git` | `sudo yum install git` | | | |
| CMake | `sudo apt-get install cmake` | (too old version) | https://cmake.org/install/ | The newest versions (starting 3.16) require `sudo apt-get install libssl-dev` or `sudo yum install openssl-devel` | 3.5.1 |

| Requirement | Ubuntu/Debian | CentOS | Manual installation | Comments | Tested version |
|---|---|---|---|---|---|
| Boost C++ libraries | `sudo apt-get install libboost-all-dev` | (version not working) | https://www.boost.org/doc/libs/1_71_0/more/getting_started/unix-variants.html | Only download and extraction in section 1 and library builds in section 5 are strictly needed, if you set a prefix you need to set `BOOST_ROOT` to this `prefix` before the installation process below or you will get boost library errors at the cmake and make step. If you manually installed g++ run `./b2` without sudo so the environment variables `CXX` and `CC` are found. | 1.67.0 |
| SWIG | `sudo apt-get install swig` | (too old version) | http://www.swig.org/Doc4.0/Preface.html | If you set a prefix you need to add prefix/bin to your PATH variable | 3.0.8 |

### Installation

To install to the standard folder `usr/local` or to keep everything in the build folder:

```
cd /where/you/want/to/build/ReSeq
git clone https://github.com/schmeing/ReSeq.git
cd ReSeq
mkdir build
cd build
cmake ..
make
```

To install to a different folder the same steps apply but the `cmake ..` line has to be exchange with:

```
cmake -DCMAKE_INSTALL_PREFIX=/where/you/want/to/install/ReSeq/ ..
```

The executable file will afterwards be `/where/you/want/to/build/ReSeq/ReSeq/build/bin/reseq` and can be added to the PATH variable or copied to the desired place.

Alternatively ReSeq can be install to the standard folder `usr/local` or the previously defined folder by:

```
make install
```

To test the installation run:

```
reseq test
```

Some useful python scripts can be found in `/where/you/want/to/install/ReSeq/ReSeq/python` or after an installation in `usr/local/bin` or `/where/you/want/to/install/ReSeq/bin/`.

### Quick start examples

To create simulated data similar to real data you first need to map the real data to a reference. For example with `bowtie2`:

```
bowtie2-build my_reference.fa my_reference
bowtie2 -p 32 -X 2000 -x my_reference -1 my_data_1.fq -2 my_data_2.fq | samtools sort -m 10G -@ 4 -T _tmp -o my_mappings.bam -
```

To run the full simulation pipeline (Stats creation, Probability estimation, Simulation) execute:

```
reseq illuminaPE -j 32 -r my_reference.fa -b my_mappings.bam -1 my_simulated_data_1.fq -2 my_simulated_data_2.fq
```

The same is done by the following three commands for the three different steps (So you can run for example only the simulation the second time you want to simulate from the same real data):

```
reseq illuminaPE -j 32 -r my_reference.fa -b my_mappings.bam --statsOnly
reseq illuminaPE -j 32 -s my_mappings.bam.reseq --stopAfterEstimation
reseq illuminaPE -j 32 -R my_reference.fa -s my_mappings.bam.reseq --ipfIterations 0 -1 my_simulated_data_1.fq -2 my_simulated_data_2.fq
```

In order to add variation (to simulate diploid genomes or populations) the parameter `-V` needs to be added:

```
reseq illuminaPE -j 32 -r my_reference.fa -b my_mappings.bam -V my_variation.vcf -1 my_simulated_data_1.fq -2 my_simulated_data_2.fq
```

or

```
reseq illuminaPE -j 32 -R my_reference.fa -s my_mappings.bam.reseq -V my_variation.vcf --ipfIterations 0 -1 my_simulated_data_1.fq -2 my_simulated_data_2.fq
```

To run a simulation with tiles the tile information needs to stay in the read names after the mapping. This means there must not be a space before it, like it is often the case for read archive data. To replace the space on the fly with an underscore the reseq-prepare-names.py script is provided. In this case run the mapping like this:

```
bowtie2 -p 32 -X 2000 -x my_reference -1 <(reseq-prepare-names.py my_data_1.fq my_data_2.fq) -2 <(reseq-prepare-names.py my_data_2.fq my_data_1.fq) | samtools sort
```

### Parameter

`reseq getRefSeqBias [options]`

| Parameter | Default | Description |
|---|---|---|
| **General** | | |
| `-h` `--help` | | Prints help information and exits |
| `-j` `--threads` | 0 | Number of threads used (0=auto) |
| `--verbosity` | 4 | Sets the level of verbosity (4=everything, 0=nothing) |
| `--version` | | Prints version info and exits |
| **ReplaceN** | | |
| `-o` `--output` | None | Output file for the reference sequence bias (tsv format) |
| `-r` `--ref` | None | Reference sequences in fasta format (gz and bz2 supported) |
| `-s` `--stats` | None | Reseq statistics file to extract reference sequence bias |

`reseq illuminaPE [options]`

| Parameter | Default | Description |
|---|---|---|
| **General** | | |
| `-h` `--help` | | Prints help information and exits |
| `-j` `--threads` | 0 | Number of threads used (0=auto) |
| `--verbosity` | 4 | Sets the level of verbosity (4=everything, 0=nothing) |
| `--version` | | Prints version info and exits |
| **Stats** | | |
| `--adapterFile` | (AutoDetect) | Fasta file with adapter sequences |
| `--adapterMatrix` | (AutoDetect) | 0/1 matrix with valid adapter pairing (first read in rows, second read in columns) |
| `-b` `--bamIn` | None | Position sorted bam/sam file with reads mapped to `--refIn` |
| `--binSizeBiasFit` | 100000000 | Reference sequences large then this are split for bias fitting to limit memory consumption |
| `--maxFragLen` | 2000 | Maximum fragment length to include pairs into statistics |
| `--minMapQ` | 10 | Minimum mapping quality to include pairs into statistics |
| –noBias | | Do not perform bias fit. Results in uniform coverage if simulated from |
| `--noTiles` | | Ignore tiles for the statistics [default] |
| `-r` `--refIn` | None | Reference sequences in fasta format (gz and bz2 supported) |
| `--statsOnly` | | Only generate the statistics |
| `-s` `--statsIn` | None | Skips statistics generation and reads directly from stats file |
| `-S` `--statsOut` | `--bamIn` .reseq | Stores the real data statistics for reuse in given file |
| `--tiles` | | Use tiles for the statistics |
| `-v` `--vcfIn` | None | Ignore all positions with a listed variant for stats generation |
| **Probabilities** | | |
| `--ipfIterations` | 200 | Maximum number of iterations for iterative proportional fitting |
| `--ipfPrecision` | 5 | Iterative proportional fitting procedure stops after reaching this precision (%) |
| `-p` `--probabilitiesIn` | `--statsIn` .ipf | Loads last estimated probabilities and continues from there if precision is not met |
| `-P` `--probabilitiesOut` | `--probabilitiesIn` | Stores the probabilities estimated by iterative proportional fitting |
| `--stopAfterEstimation` | | Stop after estimating the probabilities |

| Parameter | Default | Description |
|---|---|---|
| **Simulation** | | |
| `-1` `--firstReadsOut` | reseq-R1.fq | Writes the simulated first reads into this file |
| `-2` `--secondReadsOut` | reseq-R2.fq | Writes the simulated second reads into this file |
| `-c` `--coverage` | 0 | Approximate average read depth simulated (0 = Corrected original coverage) |
| `--errorMutliplier` | 1.0 | Divides the original probability of correct base calls(no substitution error) by this value and renormalizes |
| `--methylation` | | Extended bed graph file specifying methylation for regions. Multiple score columns for individual alleles are possible, but must match with vcfSim. C->T conversions for 1-specified value in region. |
| `--numReads` | 0 | Approximate number of read pairs simulated (0 = Use `--coverage` ) |
| `--readSysError` | None | Read systematic errors from file in fastq format (seq=dominant error, qual=error percentage) |
| `--recordBaseIdentifier` | ReseqRead | Base Identifier for the simulated fastq records, followed by a count and other information about the read |
| `--refBias` | keep/no | Way to select the reference biases for simulation (keep [from refIn] / no [biases] / draw [with replacement from original biases] / file) |
| `--refBiasFile` | None | File to read reference biases from: One sequence per file (identifier bias) |
| `-R` `--refSim` | `--refIn` | Reference sequences in fasta format to simulate from |
| `--seed` | None | Seed used for simulation, if none is given random seed will be used |
| `-V` `--vcfSim` | None | Defines genotypes to simulate alleles or populations |
| `--writeSysError` | None | Write the randomly drawn systematic errors to file in fastq format (seq=dominant error, qual=error percentage) |

`reseq replaceN [options]`

| Parameter | Default | Description |
|---|---|---|
| **General** | | |
| `-h` `--help` | | Prints help information and exits |
| `-j` `--threads` | 0 | Number of threads used (0=auto) |
| `--verbosity` | 4 | Sets the level of verbosity (4=everything, 0=nothing) |
| `--version` | | Prints version info and exits |
| **ReplaceN** | | |
| `-r` `--refIn` | None | Reference sequences in fasta format (gz and bz2 supported) |
| `-R` `--refSim` | None | File to where reference sequences in fasta format with N's randomly replace should be written to |
| `--seed` | None | Seed used for replacing N, if none is given random seed will be used |

### File Formats

| File type | Ending | Information |
|---|---|---|
| Reference file | .fa | Standard reference file in fasta format. Everything not an A,C,G,T is randomly replaced by one of those before simulating. To consistently simulate the same base over multiple simulations use the replaceN mode of reseq before starting the simulation to create a reference without N(or other ambiguous bases, which are all treated as N). A single reference sequence is only supported up to a maximum length of 4294967295 bases. The complete reference can be much bigger. |
| Mapping file | .bam | Standard mapping file in bam format. Reference information need to match the provided reference. To include tiles the tile information needs to stay in the QNAME field. Only primary alignments are used. |
| Adapter file | .fa | File in fasta format giving the sequences of possibly used adapters. The shorter this list the less missidentification can happen. Some files to use are in the adapter folder. The direction of the adapters is irrelevant as always the adapter given and its reverse complement are checked, due to sequencing-machine dependence on the direction. |
| Adapter matrix | .mat | 0/1 matrix stating if the adapters can occur in a read pair together. (0:no; 1:yes). The n-th row/column is the n-th entry in the adapter file. Rows represent the adapter in the first read and columns represent the adapter in the second read. Columns are consecutive digits in a row. Number of rows and columns need to match the number of entries in the adapter file. |

| File type | Ending | Information |
|---|---|---|
| Variant file | .vcf | Standard vcf format. The reference information in the header and in the reference column must match the given reference file. Ambiguous bases (e.g. N's) are not supported in the reference and alternative column. Except of this only the CHROM, POS, ALT columns and the genotype information are used. No filtering by quality etc. takes place. All genotypes in the file will be simulated. No distinction is made if genotypes are in a single sample or spread out over multiple samples. All genotype information is considered phased independent of what is encoded in the file. At most 64 genotypes are currently supported. |
| Methylation file | .bed | Extended bed graph format with possibility of multiple score columns for individual alleles. Number of columns must be either 1 or the same number of alleles specified in the variant file. The number of columns need to be the same within each reference sequence. Bisulfite sequencing is simulated, so C->T conversions are inserted with a probability of 1-methylation value specified in this file. |
| Stats file | .reseq | Boost archive: Not recommended to modify or create by hand even though it is in ASCII format |
| Probability file | .reseq.ipf | Boost archive: Not recommended to modify or create by hand even though it is in ASCII format |
| Systematic error file | .fq | Standard fastq format. The sequence represents the dominant error and the quality the error rate in percent at that position. There are two entries per reference sequence. The order of the reference sequences must be kept. The length must match the length of the reference sequence. The first entry per reference sequence is the reverse strand and reverse complemented. So an A in the first position means that a systematic error towards a T is simulated for the last base of the reference sequence in reads on the reverse strand. The second entry is the forward strand taken as is, so not reverse complemented. The error rate in percent is encoded similar to quality values with an offset of 33. Since the fastq format is limited to 94 quality values odd percentages over 86 are omitted. This mean `~` encodes 100% and `}` 98%. |
| Reference bias file | .txt | One line per specified bias. The line starts by a unique identifier of the reference sequence (the part before the first space in the reference sequence name in the reference file). The identifier can be followed by a space and after it some arbitrary information. The line ends with a space or tab separating a floating point number representing the bias for this sequence. It must be positive. All reference sequences in the reference file must have a bias given. However, the order of the sequences doesn't need to be kept and the bias for additional reference sequences could be specified. |

### Included libraries

Googletest (https://github.com/google/googletest.git, BSD 3-Clause license)\ NLopt (https://github.com/stevengj/nlopt.git, MIT license)\ SeqAn (https://github.com/seqan/seqan, BSD 3-Clause license)\ skewer (https://github.com/relipmoc/skewer, MIT license, made slight adaptations to the code to be able to include it)

### Publication

In preparation
